# Tubulin E-hook Hexamers Reveal Charge Dependent Compaction and Transient Secondary Structure Signatures

**DOI:** 10.64898/2026.08.11.744204

**Authors:** Alexander C. Bromley, Nicholas A. Kruse, Connor R. Brower, Madison K. Beam, Nathan I. Hammer, Ryan C. Fortenberry, Dana N. Reinemann

## Abstract

This present work shows that E-hook fragments possess functional structure differences governed by electrostatic interactions and sequence composition. The acidic C-terminal tails of tubulin, known as E-hooks, play a central role in regulating interactions between microtubules and motor proteins, microtubule-associated proteins, and enzymatic modifiers. Despite their functional importance, the intrinsic structural properties of these peptide segments remain poorly characterized due to their intrinsically disordered nature. In this work, we present quantum-mechanically optimized structures of hexamer peptides derived from β-tubulin E-hook sequences. Density functional theory calculations were used to optimize peptide geometries using progressively larger basis sets. From the optimized geometries we calculated theoretical Raman spectra, Ramachandran backbone dihedral distributions, and measured radii of gyration to resolve composition dependent structural tendencies. The combined Raman and conformational analyses provide a systematic computational approach for comparing simulated and experimental Raman spectra of tubulin E-hooks and other intrinsically disordered proteins and offer insight into how E-hooks contribute to the recognition mechanisms underlying the tubulin code.

## 1. Introduction

Microtubules are dynamic cytoskeletal polymers that perform essential cellular functions including intracellular transport, chromosome segregation during mitosis(1), motility(2), and other stages of cellular evolution(3). These polymers are composed of α- and β-tubulin heterodimers(4, 5) that assemble into protofilaments which join to form the hollow cylindrical microtubule lattice(6). Although the structured domains of the tubulin core have been extensively characterized through X-ray crystallography and cryo-electron microscopy(7), the C-terminal tails of tubulin remain largely unresolved in structural studies(8).

Microtubule’s C-terminal tails, commonly referred to as E-hooks, are short peptide segments rich in glutamic acid residues, giving them their namesake, and are characterized by high negative charge density(9). E-hooks extend outward from the microtubule surface, acting like an electronegative brush for motor proteins, microtubule-associated proteins (MAPs), and severing enzymes, affecting their binding capabilities and functionality(10–12). E-hooks are thought to be modulated by two major structural factors that influence MAPs: isotype expression(13, 14) and post-translational modification(15, 16). The functional importance of E-hooks and the regulatory function of their structure have led to the concept of the “tubulin code,” wherein the combinations of the tubulin isotypes and post-translational modifications regulate microtubule behavior and function by modulating MAP interactions(17, 18).

Despite their biological significance, the structural properties of E-hooks remain poorly understood. Because of their intrinsic flexibility from the electrostatic repulsion between their acidic residues, these segments are widely believed to behave as intrinsically disordered peptides, making traditional experimental structure studies, such as X-ray crystallography and cryo-electron microscopy, ineffective(8). However, growing evidence from alternative structural techniques suggests that intrinsically disordered regions may still exhibit transient local secondary structures that may contribute to molecular recognition(19). Novel approaches, such as computational chemistry, must be employed in order to examine how the structures of these molecules influence their environments.

E-hook structures have been studied in the past using molecular dynamics (MD)(19–22). Conformational ensembles were originally prepared by allowing a free E-hook to explore its conformational space. While the E-hooks did portray the predicted characteristics of intrinsically disordered proteins, the MD simulations predict that they also demonstrated a consistent yet transient helical secondary structure(19). Further MD studies with E-hooks attached to tubulin’s globular core have revealed the existence of anchoring points, wherein certain E-hook residues may non-covalently interact with the globular core(21). These specific residues of the E-hook and globular core that interacted as attachment points varied between simulations yet remained in place throughout each simulation, suggesting that E-hooks may have their conformational landscape limited in some capacity(20). Other computational studies have used MD to examine how E-hooks affect MAPs and motor protein binding. These computations have revealed a soft, guided landing functionality of E-hooks(23, 24). By providing a broad brush of negative charge, domains with positive residues are drawn toward the microtubule, but upon nearing the filament, the microtubule binding domain’s (usually along the globular core) begins to control the MAP’s microtubule binding due to its stronger binding affinity (23, 25). While MD can show how E-hooks may change over time and how they may affect dynamic processes, minute differences in intramolecular interactions between the different E-hook isotypes cannot be effectively predicted, preventing deeper understanding for how E-hook structure affects microtubule functionality(26).

Time-independent quantum chemical modeling provides the precision necessary to study intramolecular interactions leading to these transient secondary structures and how they differ across E-hook sequence composition and post-translational modification. However, the increased cost of this approach prevents its usage across a conformational ensemble(27). Density functional theory (DFT) computations can both optimize a given molecular geometry and predict its associated vibrational frequencies, allowing theoretical spectra to be compared to directly to experimental measurements(28). Raman spectroscopy, on the other hand, provides an experimental means to probe peptide structure in flexible, biological systems(29). This technique is particularly useful in the study of E-hooks due to backbone amide and side chain vibrational modes providing detailed information about peptide conformation and hydrogen bonding, allowing for experimental corroboration with computationally derived models(29–33).

In this study, quantum mechanically optimized structures of E-hook hexamer peptides are presented. These are derived from the C-terminal of β-tubulin sequences (**Table 1**) and are used to quantify each simulated geometries’ agreement with experiment as well as map E-hook Raman fingerprints. The C-terminal hexamers of the E-hooks are used as representative peptides due to the computational cost of quantum mechanical calculations(34, 35). In addition, this region of the E-hook hosts the major compositional differences between the isotypes and would likely be the portion closest to MAPs and motor proteins attempting to bind to the microtubule, making their conformational information relevant to functional translation(8, 9, 36). Our approach integrates DFT structural optimizations, theoretical Raman spectra calculations, Ramachandran backbone analysis, and structural compactness measurements through radius of gyration calculations. By combining these methods, the aim of the present work is to characterize intrinsic structural tendencies of E-hooks and examine how electrostatics and sequence composition influence their conformational behavior.

**Table 1:**
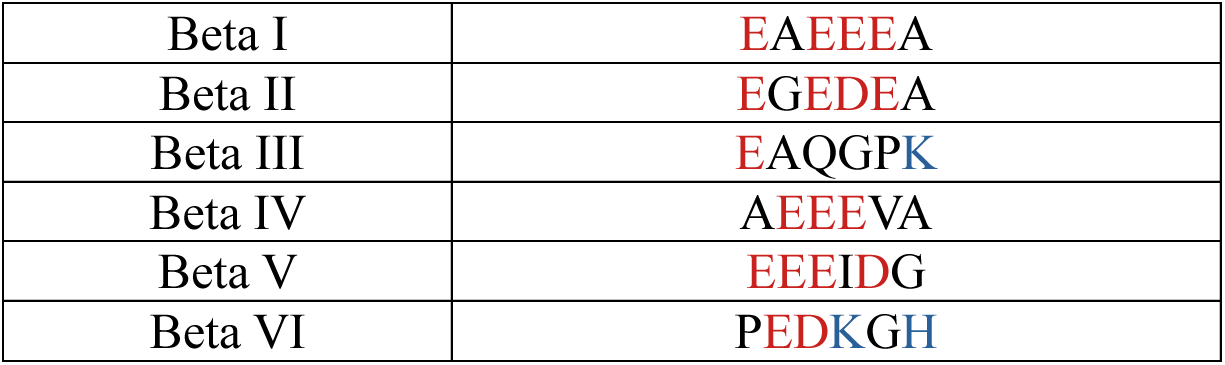
E-hooks and their sequences modelled and characterized in this study. Red denotes a negatively charged residue, and blue denotes a positively charged residue at pH 7. Histidine is partially protonated at this pH; however, its hydrogen-bonding capacity and the highly acidic local environment are expected to increase its effective pKa, giving it a higher protonation and positive charge. Sequences were obtained from the NCBI protein database(37).

## 2. Methods

### 2.1 Peptide Structure Construction

Structures of E-hook hexamers corresponding to C-terminal regions of β-tubulin E-hook sequences are constructed. A stepwise buildup method for constructing E-hooks has been previously developed using quantum chemical geometry optimizations(34, 35) and is described briefly here. Each hexamer sequence is divided into two residue structures. These structures are iteratively optimized in Gaussian16(38) using density functional theory (DFT) with the Becke three-parameter Lee-Yang-Parr (B3LYP) exchange-correlation functional with basis sets increasing in size. Basis sets 3-21G(39), 6-31G(40), 6-31+G(*d*,*p*)(41), and 6-311+G(*2df*,*2pd*)(42) are utilized with the last two incorporating diffuse and polarization functions to improve the description of intramolecular interactions and the diffuse electron environment of E-hooks, respectively. Upon optimization of the structure with the final basis set, the dimer is joined with another structurally optimized dimer, creating a tetramer that is then subjected to the same iterative optimization. Tetramers are formed from both the N-terminus and the C-terminus of the representative hexamer, allowing for the creation of two final hexamers, expanding the conformations available from previous studies. These tetramers are then joined to the final, separately optimized dimer, creating two hexamers which are subsequently subjected to the iterative optimization again (**Figure 1**). These hexamers are named according to their reconstruction strategy: geometries made with the N-terminal tetramer and C-terminal dimer are labelled as 4+2 (e.g. EAEE+EA), and those made with the N-terminal dimer and C-terminal tetramer are labelled as 2+4 (e.g. EA+EEEA). The final hexamers are visualized and analyzed for hydrogen bonding in Jmol(43) based on the Baker-Hubbard definition where the distance between the hydrogen and acceptor heavy atom is less than or equal to 2.5 Å and the hydrogen bond angle is greater than or equal to 120°(44–46), criteria commonly employed to identify geometrically favorable hydrogen bonds while excluding weak or non-directional contacts.

**Figure 1.**
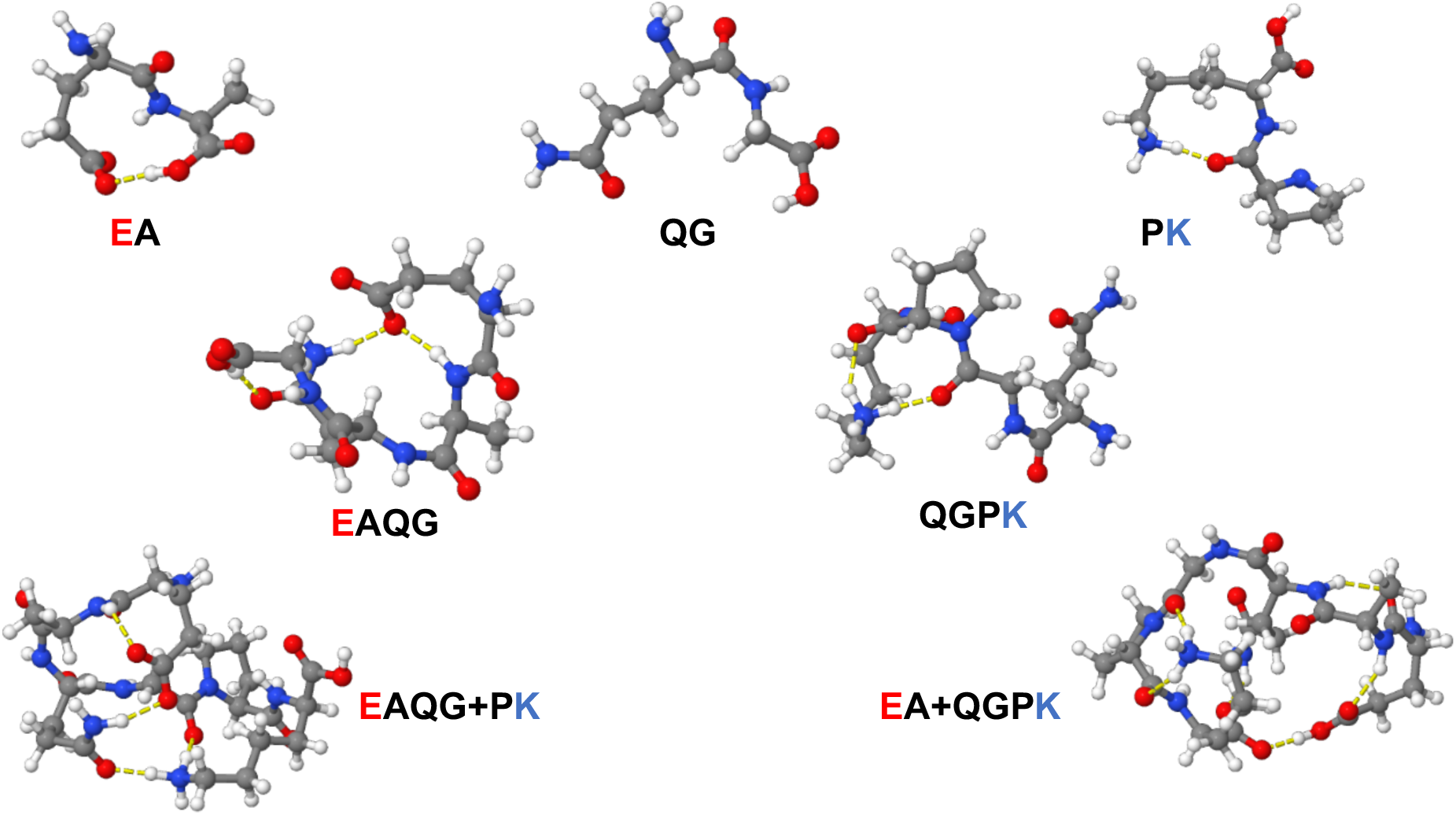
An example of the iterative optimization buildup method used to construct full hexamer E-hooks. Three dimers make two tetramers which, in turn, make the same hexamer but with two different conformations. Shown here is the process to build up to full hexamers of Beta III with hydrogen bonds shown in yellow.

**Figure 2.**
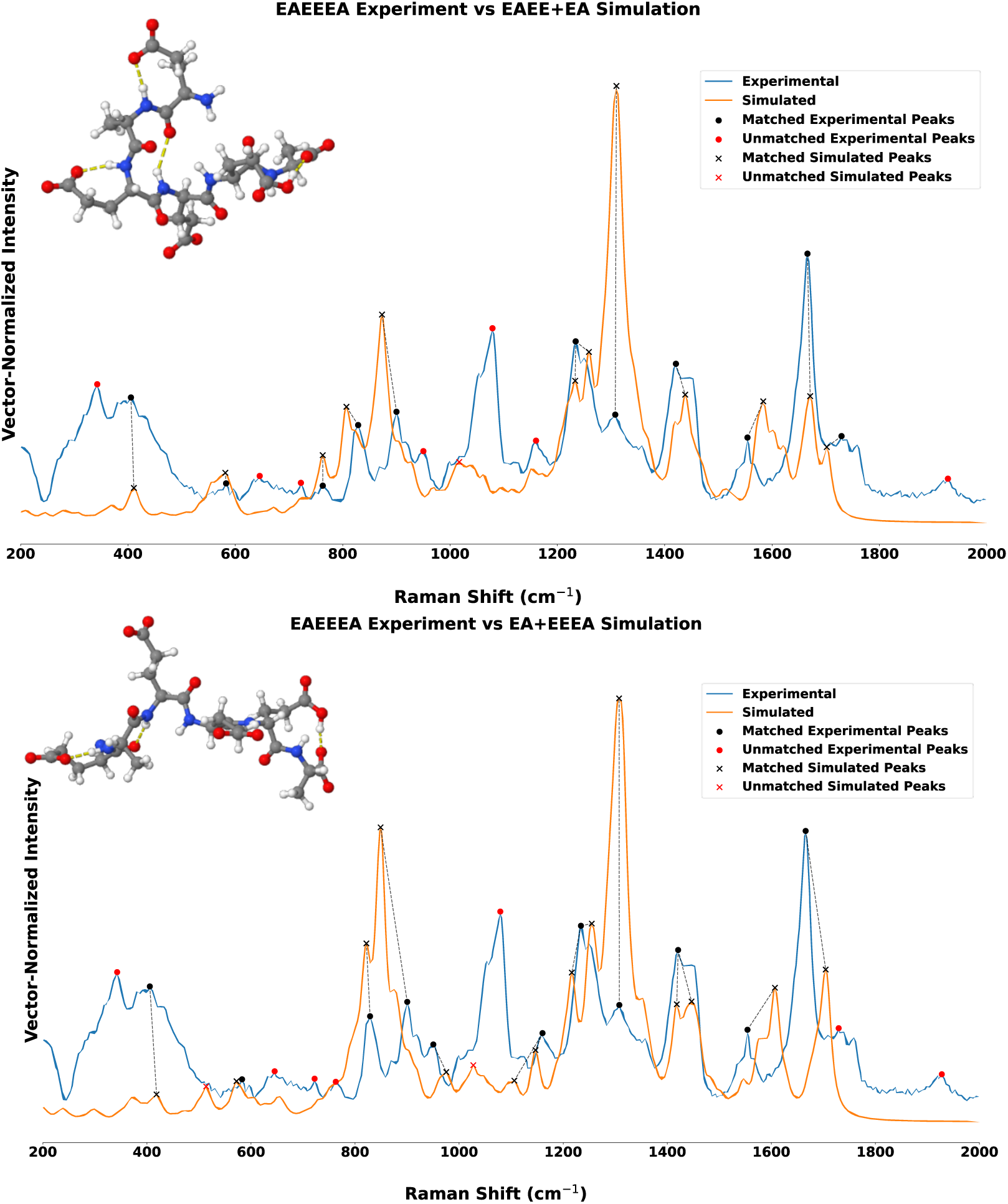
Raman spectral comparison of βI-tubulin E-hook hexamer (EAEEEA) experimental spectra with simulated spectra generated from alternative peptide construction strategies. The EAEE+EA (top) geometry demonstrates improved agreement with experimentally observed vibrational frequencies relative to the EA+EEEA (bottom) reconstruction, particularly through reduced peak-position deviations.

**Figure 3.**
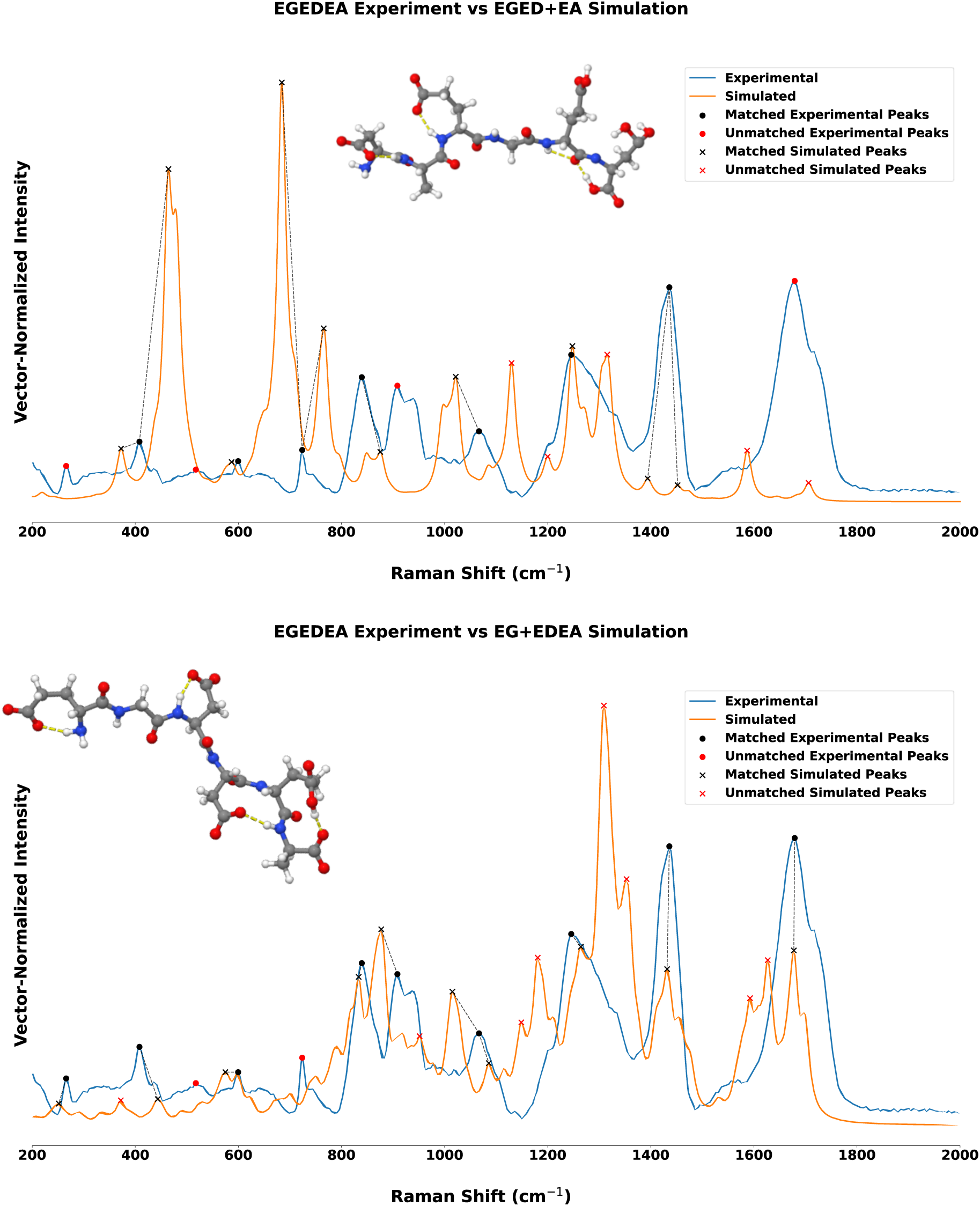
Raman spectral comparison of βII-tubulin E-hook hexamer (EGEDEA) experimental spectra with simulated spectra generated from alternative peptide construction strategies. The EA+EDEA (bottom) geometry demonstrates improved agreement with experimentally observed vibrational frequencies relative to the EGED+EA (top) reconstruction, particularly through reduced peak-position deviations and increased signal intensity similarity.

**Figure 4.**
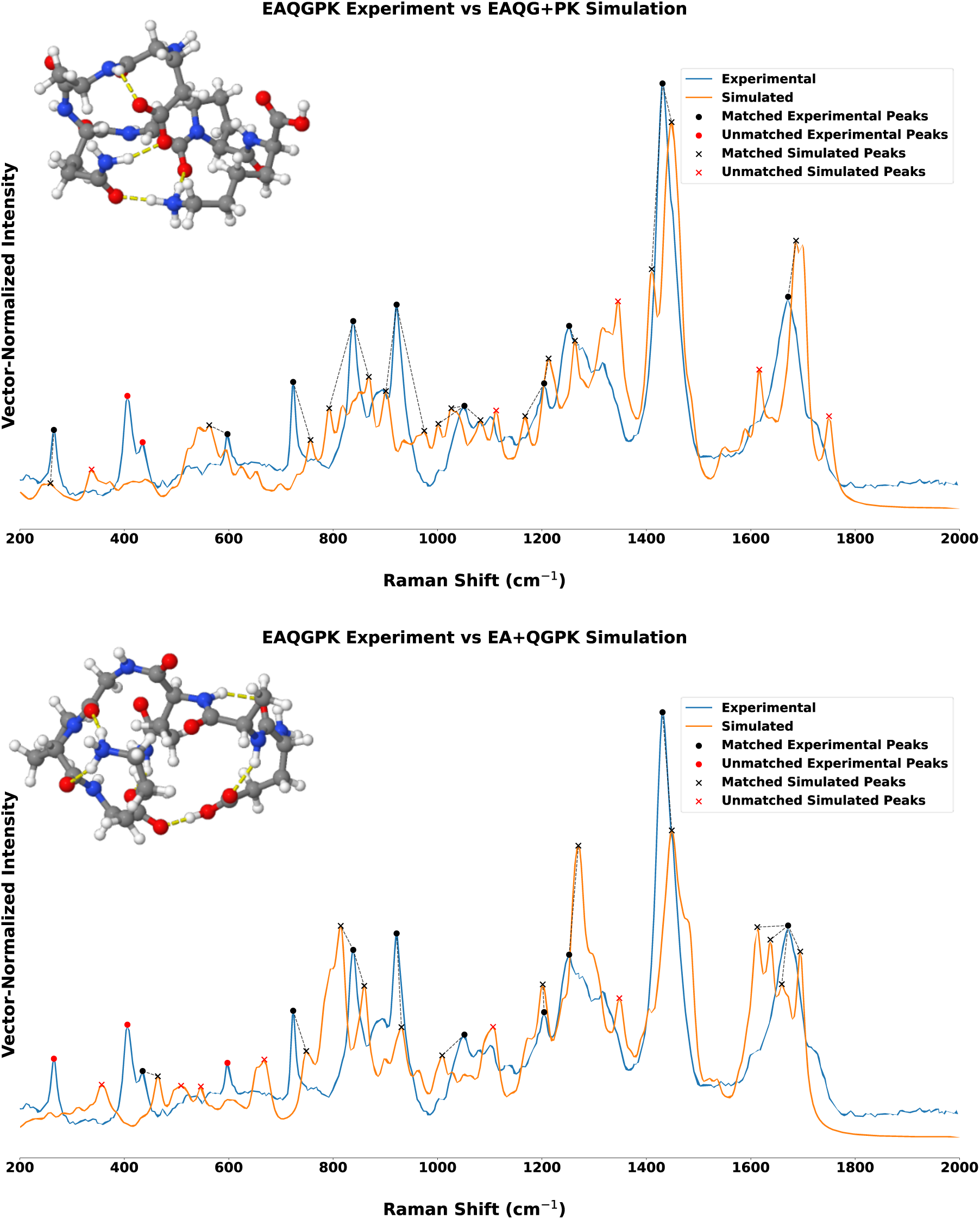
Raman spectral comparison of βIII-tubulin E-hook hexamer (EAQGPK) experimental spectra with simulated spectra generated from alternative peptide construction strategies. The EAQG+PK (top) geometry demonstrates improved agreement with experimentally observed vibrational frequencies relative to the EA+QGPK (bottom) reconstruction, particularly through reduced peak-position deviations.

**Figure 5.**
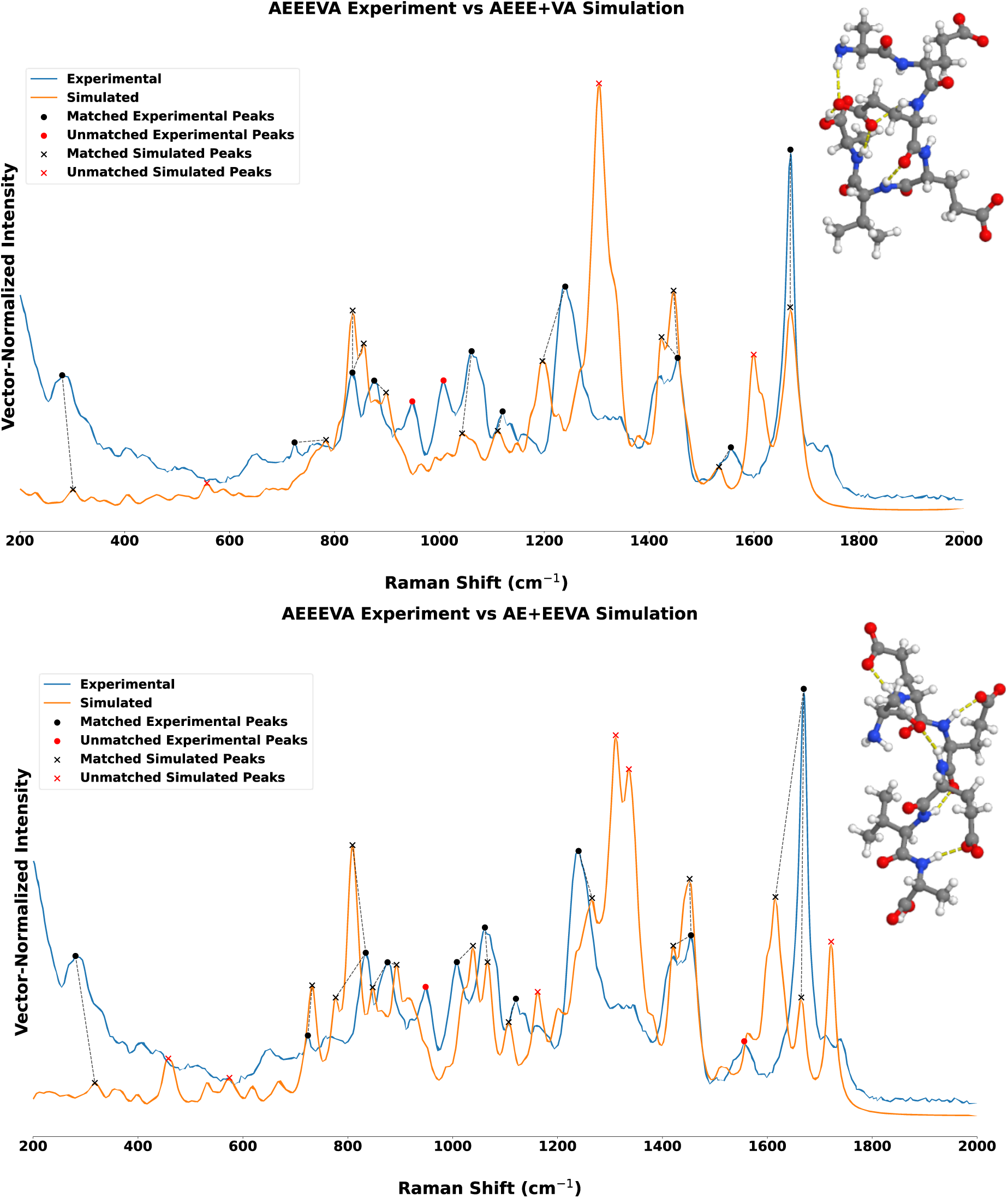
Raman spectral comparison of βIV-tubulin E-hook hexamer (AEEEVA) experimental spectra and its two simulated geometries. AEEE+VA reconstruction (top) shows slightly improved peak shift RMSD and Pearson coefficient compared to the AE+EEVA geometry (bottom).

**Figure 6.**
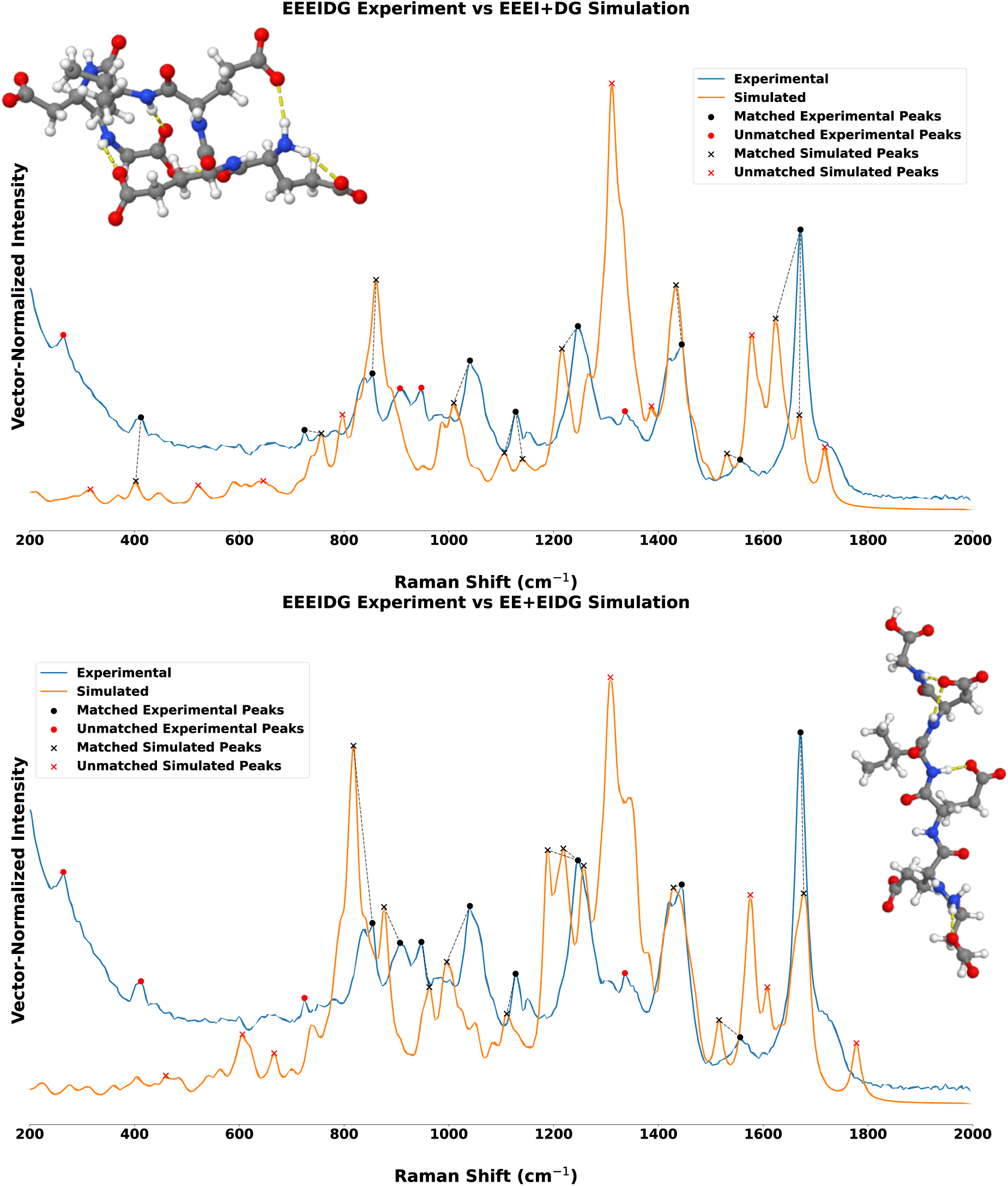
Raman spectral comparison of βV-tubulin E-hook hexamer (EEEIDG). Reconstruction using the EEEI+DG fragmentation strategy (top) substantially improves peak-position agreement despite reduced intensity correlation compared to EE+EIDG (bottom), indicating improved recovery of experimentally observed vibrational frequencies independent of exact intensity reproduction.

**Figure 7.**
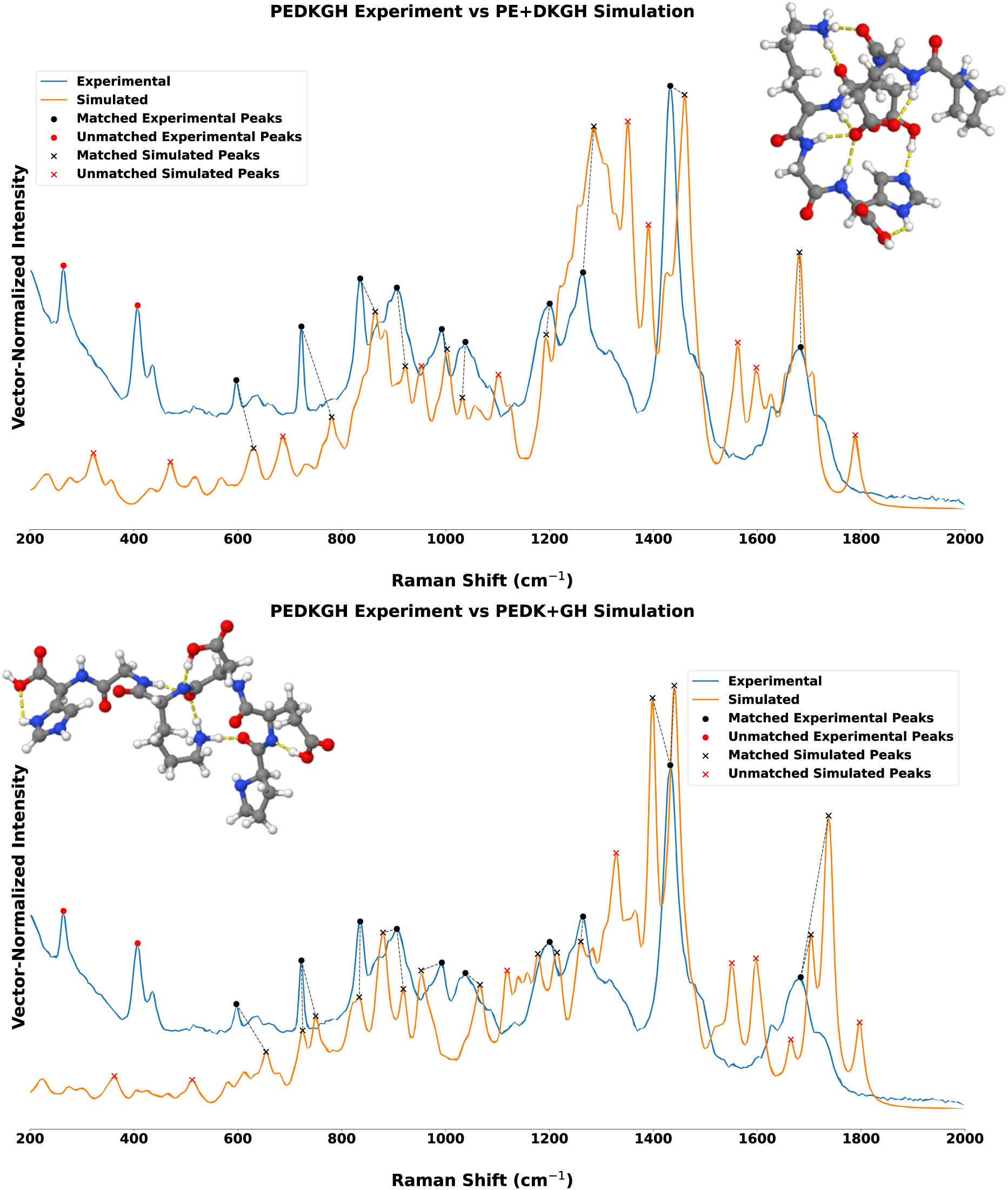
Experimental Raman spectra of the βVI-tubulin E-hook hexamer (PEDKGH) overlaid with simulated spectra. The PE+DKGH reconstruction (bottom) demonstrates improved overall agreement with experimentally observed vibrational frequencies relative to the PEDK+GH (top) geometry, although both structures exhibit comparatively similar spectral behavior.

### 2.2 Vibrational Frequency Analysis and Experimental Raman Spectra

Following each geometry optimization, harmonic vibrational frequency calculations are performed within Gaussian16 to obtain Raman intensities and frequencies. A scaling factor of 0.97 is applied to account for anharmonicity of the fundamental vibrational frequencies, and the spectra are plotted using a Lorentzian line function(47).

For experimental studies, chemically synthesized E-hooks, both full length and hexamers, have been obtained from GenScript (lyophilized, purity ≥ 95%). Experimental Raman spectra are produced using a Horiba LabRAM HR Evolution Spectroscopy system with the E-hooks in a solid state using 600 or 1800 grooves/mm gratings, 532-nm laser excitation and CCD camera detection. These experimental spectra are normalized and vertically offset for direct comparison, allowing for experimental corroboration of the quantum chemical results.

### 2.3 Raman Spectra Analysis

Comparison of computed and experimental Raman spectra is frequently performed through qualitative visual inspection in which corresponding peak are identified manually and the overall agreement is assessed by the investigator(34, 35, 48–50). While this approach can be effective for simple spectra, it becomes increasingly subjective for complex biomolecular systems containing numerous overlapping bands, broadened experimental features, and systematic frequency shifts arising from harmonic quantum chemical approximations. Consequently, different investigators may emphasize certain spectral regions or assign peak correspondences differently, reducing reproducibility and making quantitative comparisons between computational methods difficult. The motivation for this present methodology, therefore, is to replace qualitative visual assessment with a systematic, reproducible, and objective comparison framework. Due to the mixed and shifted nature of these simulated spectra, this approach is used less as a means to exhaustively assign vibrational modes but more as a means to determine which buildup strategy most accurately reproduces experimentally-observed vibrational frequency positions and peak distributions.

Raman spectra are compared using a region-specific dynamic time warping approach. Experimental and simulated spectra are normalized within the fingerprint region (200-2000 cm^-1^), and the simulated spectrum is interpolated onto the experimental wavenumber axis. Both spectra are smoothed using a Gaussian filter. Dynamic time warping (DTW) is then applied to the simulated spectrum with its experimental counterpart as the reference, allowing the simulated spectrum to stretch and warp to maximize overlap between the two spectral shapes while minimizing the cost of warping to achieve the overlap. By warping the spectrum, inherit shifting from the quantum chemical simulation and its isolated conditions can be accounted for, allowing for assignment of local vibrational peaks in the warped form and recapitulation of this assignment in the non-warped spectra. The DTW spectra are used to identify candidate experimental peak positions corresponding to each simulated peak. Peaks are detected using the SciPy Python library’s *find_peaks* function(51), and simulated peaks are assigned to experimental peaks within a frequency tolerance of 40 cm^-1^.

Peak assignments are scored using a weighted function containing a DTW-derived wavenumber distance term, a scaled intensity-difference term, and an occupancy penalty to discourage excessive assignment of multiple simulated peaks to the same experimental peak. The assignment for each simulated peak is selected as the candidate with the lowest matching score. This initial assignment establishes the correspondence between simulated and experimental peaks, while leaving unmatched peaks unassigned. A second calculation of the peaks’ score is then conducted, allowing the occupancy penalty to be applied evenly throughout the spectrum and distributing simulated peaks if many are assigned to a single experimental peak if other experimental matches are available and appropriate. Agreement is assessed using weighted error functions incorporating normalized peak position root mean square difference (RMSD), Pearson correlation error, and peak matching error. Three different weighting schemes have been used to separately emphasize positional agreement (**Equation 1**), spectral-shape agreement (**Equation 2**), and a balanced agreement between peak position and spectral shape (**Equation 3**). These weights provide percentage windows into the different aspects of the scoring methodology, allowing for the quantitative differentiation of the multiple priorities that can be applied to an analytical spectral comparison method without negating and losing the resolution of any facet completely.

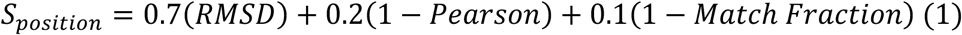

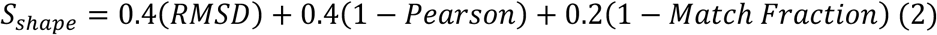

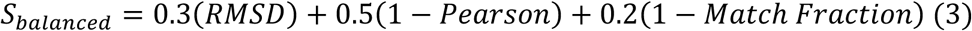

Existing Raman-matching approaches commonly rely upon point similarity scoring(52–54), peak-shift corrections(53, 55), spectral segmentation(56), or unconstrained profile warping(57). Although these methods are effective for spectral library searching and inter-instrument standardization, they are not designed for the comparison of simulated quantum-chemical spectra with experimental peptide spectra. Pointwise metrics are sensitive to frequency displacements(52, 54), whereas unrestricted warping can produce favorable similarity scores through excessive or chemically ambiguous alignment(57). The present methodology, therefore, combines constrained DTW with explicit peak detection, reduced redundant peak assignments, peak-position error, and matched-feature coverage. This framework tolerates expected calculated-to-experimental frequency shifts while retaining the magnitude of those shifts and identifying unmatched bands. It consequently provides both whole-spectrum similarity and mode-level interpretability which are both necessary for evaluating basis-set and conformational effects in tubulin E-hook Raman spectra.

### 2.4 Structural Analysis

Backbone dihedral angles φ and ψ are calculated using the atomic coordinates from the optimized structures. Ramachandran plots are generated to visualize conformational distributions with unique marker shapes for each isotype and unique marker colors for each residue. These plots allow for identification of conformational trends both within and across isotypes.

The radius of gyration (R_g_) is the root-mean-square distance of the peptide’s atoms from its center of mass and is calculated for each peptide using atomic coordinates from the electronically optimized structures. The radius of gyration is defined as

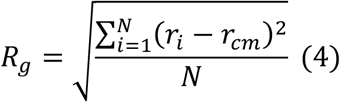

where *r_i_* represents the position of atom *i* and *r_cm_* is the peptide’s center of mass. R_g_ provides a quantitative measure for peptide compactness(58).

## 3. Results

### 3.1 Raman Spectra of E-hook Peptides

Across all β-tubulin isotypes, both the 4+2 and 2+4 buildup methods reproduce the majority of experimentally observed spectral features. However, neither the 4+2 nor the 2+4 reconstruction strategy consistently outperform the other across all sequences. Instead, the preferred reconstruction strategy is strongly dependent upon peptide composition, indicating that sequence-specific structural interactions may influence the preservation of experimentally observed vibrational behavior (**Table 2**).

**Table 2.**
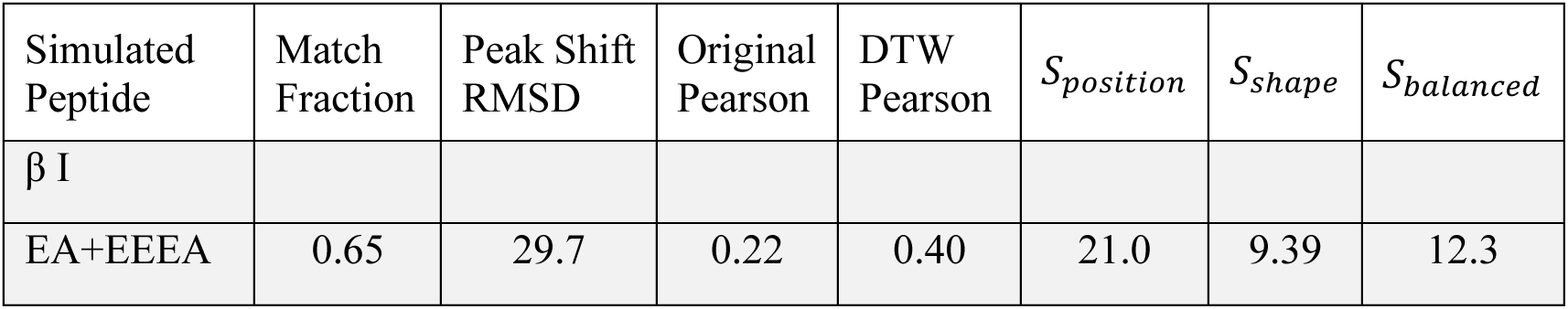
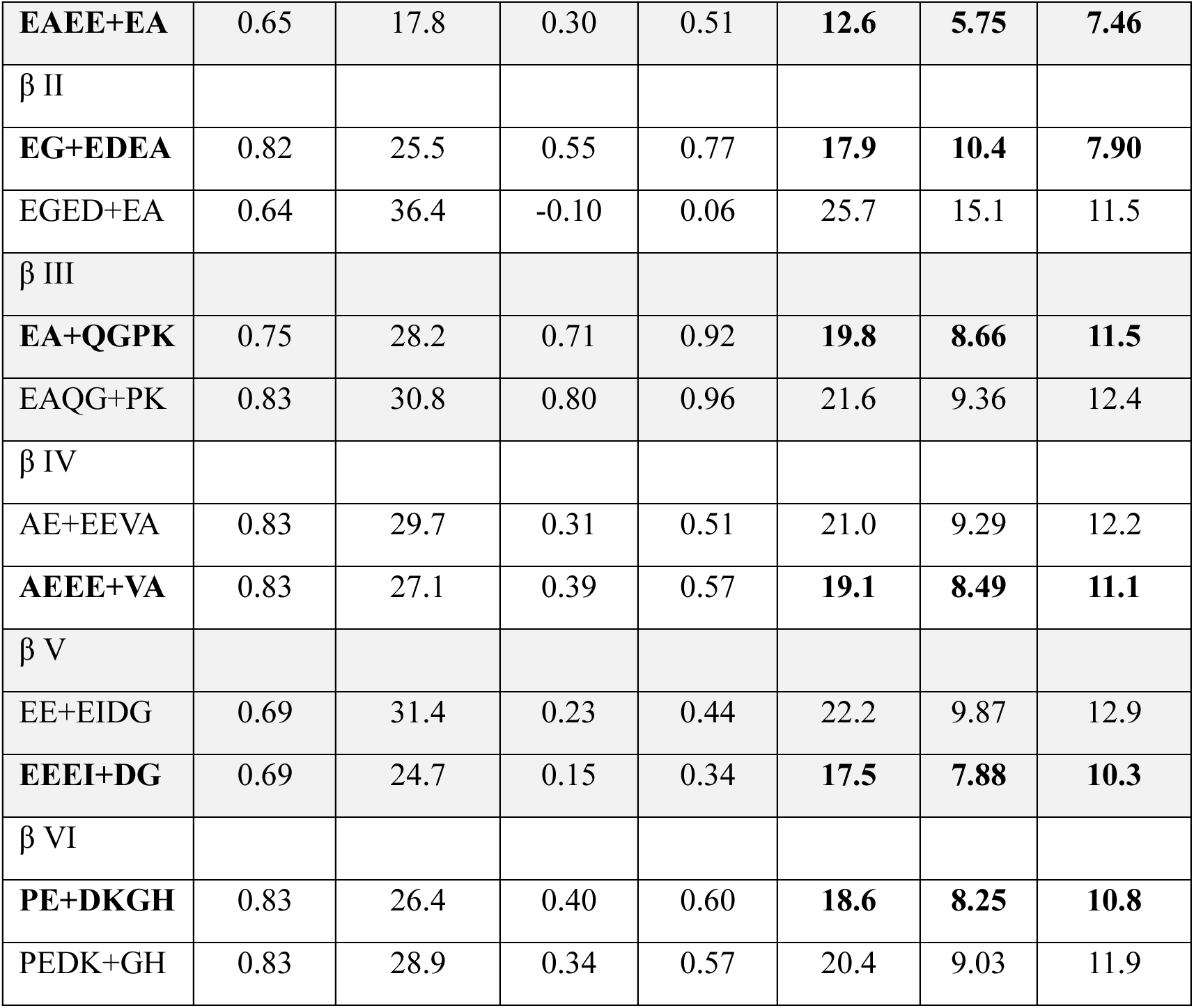
Quantitative analysis performed on all simulated β-tubulin E-hook hexamers. The three metrics used for composite scoring and ranking are shown on the left: match fraction, peak shift RMSD, and the original Pearson coefficient. The Pearson coefficient of the warped spectra is also shown for reference. Each reconstruction with the lowest composite scores (highest agreement) across each isotype is bolded.

#### 3.1.1 βI – EAEEEA

βI E-hooks (EAEEEA) demonstrate the strongest sensitivity to reconstruction strategy. The EAEE+EA (4+2) buildup reduces peak-shift RMSD from 29.7 cm⁻¹ to 17.8 cm⁻¹ while simultaneously improving the Pearson correlation from 0.215 to 0.297. The peak-matching fraction remains identical between both geometries, indicating that the improvement arises from more accurate reproduction of experimentally observed vibrational frequencies rather than recovery of additional spectral features. In the EAEE+EA spectrum, the amide III region near 1250–1260 cm⁻¹ closely reproduces the experimental feature and is assigned primarily to amide III-like C–N stretching coupled with N–H bending, reflecting improved preservation of backbone vibrational modes. Likewise, the amide I band near 1650–1660 cm⁻¹, assigned to backbone carbonyl stretching, shows close frequency agreement with experiment. Despite these improvements, the simulation continues to overestimate the intensity of the 1290–1310 cm⁻¹ feature, which is assigned largely to amide III-like and mixed collective vibrations, indicating that vibrational coupling among the glutamate-rich residues remains incompletely captured.

The alternative EA+EEEA (2+4) reconstruction predicts effectively the same principal amide III and amide I assignments but exhibits greater splitting and intensity distortion throughout the 780–820 cm⁻¹ region. This span of features corresponds primarily to skeletal bending/torsional and mixed collective vibrations. Most notably, both reconstructions fail to reproduce the broad experimental band between approximately 1050 and 1100 cm⁻¹, despite reproducing neighboring features. This region is associated with mixed collective backbone and side-chain motions, suggesting that the experimentally observed vibration likely depends on longer-range coupling or conformational averaging that is disrupted by fragmentation.

Consequently, although the EAEE+EA (4+2) reconstruction substantially improved peak-position agreement, the persistent absence of the 1050–1100 cm⁻¹ feature explains why the overall Pearson correlation remains comparatively low.

#### 3.1.2 βII – EGEDEA

The EG+EDEA (2+4) reconstruction provides the more useful description of the βII E-hooks (EGEDA) even though it produces lower composite scores, but it maintains strong agreement with the experimental Raman spectrum. Although both reconstruction pathways reproduce the dominant experimental features, the 2+4 model more consistently preserves peak positions throughout the fingerprint region, indicating that retaining the larger C-terminal fragment better maintains the local vibrational environment. In the EG+EDEA (2+4) spectrum, the amide III band near 1240–1250 cm⁻¹ closely matches the experimental feature and is assigned primarily to C–N stretching coupled with N–H bending from the vectors of the quantum chemically computed normal modes. Such an assignment is consistent with preservation of backbone amide vibrations. Likewise, the dominant 1450 cm⁻¹ feature, assigned to CH₂/CH₃ deformation (scissoring/bending), reproduces both the position and relative intensity of the experimental maximum. The amide I region near 1660 cm⁻¹, arising predominantly from backbone carbonyl stretching, also shows excellent positional agreement, indicating that the principal peptide backbone vibrations are retained despite intensity differences.

The alternative EGED+EA (4+2) reconstruction reproduces the same principal vibrational assignments but exhibits greater splitting within the 780–820 cm⁻¹ region, where mixed collective and skeletal bending/torsional vibrations generate additional simulated peaks that have not been observed experimentally. Additionally, EGED+EA is predicted to exhibit exaggerated relative intensity compared to experiment. Similarly, the 1300–1350 cm⁻¹ region contains several partially unmatched bands, suggesting increased perturbation of coupled backbone vibrations which is not present in the experimental spectrum potentially due to the broadening effect of vibrational mode mixing.

#### 3.1.3 βIII – EAQGPK

βIII E-hooks (EAQGPK) exhibit the strongest overall agreement with experiment and favor the fragmentation strategy of EA+QGPK (2+4). The EA+QGPK reconstruction produces superior composite scores and stronger global spectral agreement relative to EAQG+PK. Both simulations more closely agree with experiment than those of the other isotypes, indicating that βIII’s structural behavior is comparatively robust to fragmentation while simultaneously demonstrating a sequence-specific preference for preserving vibrational information through the 2+4 reconstruction. In the EA+QGPK spectrum, the dominant feature near 1445–1455 cm⁻¹ agrees strongly with the experimental maximum and was assigned primarily to CH₂/CH₃ deformation (scissoring/bending), reproducing both its position and relative prominence. The amide I region between 1650 and 1700 cm⁻¹ also closely matches the experimental spectrum and is attributed to backbone carbonyl stretching. A remaining discrepancy occurs in the 780–820 cm⁻¹ region, where skeletal bending/torsion and mixed collective vibrations are somewhat over-resolved, producing several closely spaced simulated peaks that exceed the complexity observed experimentally.

The EAQG+PK (4+2) reconstruction likewise reproduces the dominant amide I and CH₂ deformation bands, consistent with the overall robustness of βIII to fragmentation. However, the 1240–1260 cm⁻¹ amide III region, assigned predominantly to C–N stretching coupled with N–H bending, exhibits broader splitting and slightly poorer positional agreement than in the 2+4 reconstruction. Additional unmatched features between 1300 and 1350 cm⁻¹ suggest increased fragmentation-induced coupling among neighboring backbone modes despite preservation of the underlying vibrational assignments. These comparatively subtle differences explain why βIII maintains high spectral agreement for both reconstruction pathways while still exhibiting a measurable preference for the EA+QGPK (2+4) strategy.

#### 3.1.4 βIV – AEEEVA

βIV E-hooks (AEEEVA) favors the AEEE+VA (4+2) geometry. The peak-matching fraction remains identical between both structural models, while peak-shift RMSD, Pearson correlation, and composite scoring metrics all improve under the 4+2 reconstruction. These improvements indicate moderately enhanced reproduction of vibrational frequencies without significant changes in spectral coverage. In the AEEE+VA spectrum, the amide III region near 1240–1260 cm⁻¹ exhibits close agreement with the experimental spectrum and is assigned primarily to amide III-like C–N stretching coupled with N–H bending, with dominant contributions from the contiguous glutamate residues within the AEEE fragment. The strong amide I band near 1660–1670 cm⁻¹ is also accurately reproduced and corresponds predominantly to backbone and glutamate carbonyl stretching, indicating preservation of the principal peptide vibrational framework. A minor discrepancy remained near 1450–1470 cm⁻¹, where several calculated CH₂/CH₃ deformation modes are more distinctly resolved than in the experimental spectrum, producing slight intensity redistribution despite good positional agreement.

In contrast, the AE+EEVA (2+4) reconstruction generates a broader cluster of unmatched features between 1300 and 1360 cm⁻¹, reflecting increased splitting of modes associated with amide III and mixed collective backbone vibrations. The 800–850 cm⁻¹ region, assigned primarily to skeletal bending/torsional and mixed collective vibrations, also appears more intense than observed experimentally, reflecting altered vibrational coupling after fragmentation. Although both reconstructions reproduce the dominant amide I feature well, the improved agreement throughout the fingerprint region demonstrates that preserving the contiguous AEEE segment more effectively maintains the experimentally observed vibrational behavior.

#### 3.1.5 βV – EEEIDG

βV E-hooks (EEEIDG) exhibits substantial improvement under the EEEI+DG (4+2) reconstruction. The RMSD decreases from 31.4 cm⁻¹ to 24.7 cm⁻¹, while position-focused composite scoring improves significantly. Although the Pearson correlation decreases from 0.228 to 0.146, improved peak-position agreement is considered more significant than intensity disagreement because Raman intensities are highly sensitive to conformational averaging and local environmental effects. In the EEEI+DG spectrum, the amide III region near 1250–1300 cm⁻¹ show notably improved agreement with the experimental spectrum and is assigned primarily to amide III-like C–N stretching coupled with N–H bending involving the glutamate-rich EEEI fragment as shown in the quantum chemical computations. Likewise, the strong feature near 1625–1635 cm⁻¹ closely reproduces the experimental amide I band and is dominated by carbonyl stretching from glutamate carboxylate groups, indicating preservation of the backbone and acidic side-chain vibrational environment. A remaining discrepancy occurs near 1300–1330 cm⁻¹, where the quantum chemical computation produces an overly intense, partially unmatched band despite the correct underlying amide III assignment, suggesting altered coupling between neighboring glutamate residues following fragmentation.

In contrast, the EE+EIDG (2+4) reconstruction generates an overly pronounced feature near 800–850 cm⁻¹, corresponding to mixed collective CH₂-dominated vibrations, that substantially exceeded the experimental intensity. The amide III region around 1250–1300 cm⁻¹ is also reproduced but exhibits broader splitting and greater intensity distortion than observed experimentally. Although the amide I feature near 1700 cm⁻¹, assigned to carbonyl stretching, remains well positioned in both reconstructions, the superior agreement throughout the fingerprint region demonstrates that preserving the contiguous glutamate-rich EEEI segment in EEEI+DG reconstruction more effectively retains the experimentally observed vibrational behavior.

#### 3.1.6 βVI – PEDKGH

βVI E-hooks (PEDKGH) favored the PE+DKGH (2+4) reconstruction strategy. While both structures produce similar peak-matching fractions, lower composite scores, and improved correlation metrics consistently favor the 2+4 reconstruction. In the PE+DKGH spectrum, the strong simulated feature near 1280–1300 cm⁻¹ aligns well with the corresponding experimental band and is assigned primarily to amide III-like C–N stretching/N–H bending, with substantial contributions from the histidine-containing DKGH fragment. Likewise, the simulated carbonyl-region feature near 1680–1700 cm⁻¹ closely reproduces the experimental peak and is associated predominantly with aspartic acid carboxylic acid C=O stretching, indicating that the 2+4 reconstruction retains residue-localized acidic-group vibrations. A less satisfactory feature occurs near 1350–1400 cm⁻¹, where the simulation predicts several intense, partially unmatched peaks arising from mixed histidine N–H/C–N motions and backbone deformation, indicative of over-resolution or altered coupling of closely spaced modes.

The PEDK+GH spectrum also reproduces the dominant experimental band near 1430–1450 cm⁻¹, but the simulation resolved it into two exceptionally intense peaks. The PEDK mode assignments associate these features primarily with histidine-localized N–H and C–N motions. The relatively intense simulated feature near 1730–1750 cm⁻¹, assigned largely to amide-I/carbonyl stretching, is also stronger and more sharply defined than the corresponding experimental region. Compared with βI and βV, however, βVI demonstrates comparatively limited sensitivity to fragmentation, which may indicate more localized preservation of vibrational behavior across both reconstruction pathways.

### 3.2 Structural Compactness Corresponds with Raman Reconstruction Preferences and Sequence

To characterize the global structural compactness across β-tubulin E-hook isotypes, the R_g_ (**Equation 4**) is calculated for optimized peptide structures across four progressively larger basis sets (**Table 3**). R_g_ values vary substantially between isotypes and reconstruction strategy, indicating significant sequence-dependent and differences in structural compactness.

**Table 3.**
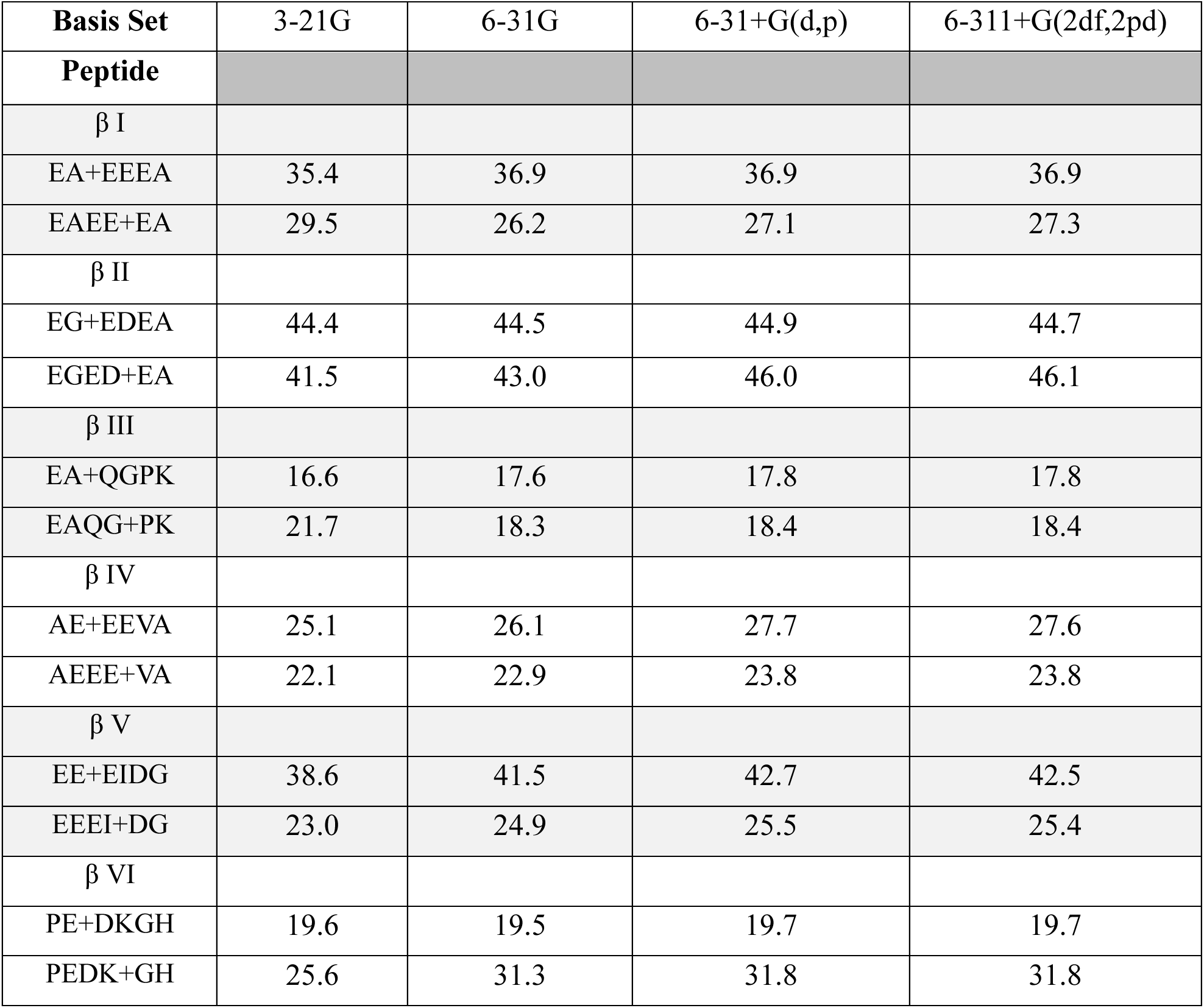
Average R_g_ of each β-tubulin E-hook isotype studied by basis set. βIII and βVI have lower R_g_ values due to their basic residues.

βIII E-hooks and βVI’s 2+4 reconstruction exhibit the lowest radii of gyration across all basis sets with values remaining approximately between 17-22 Å and 19-20 Å, respectively. In contrast, βI, βII, and βV E-hooks produce substantially larger R_g_ values ranging from approximately 26-46 Å, indicating significantly more extended conformational behavior.

Interestingly, the more compact βIII and βVI isotypes favor the 2+4 reconstruction strategy during Raman spectral analysis, whereas the more extended βI and βV structures strongly favor the 4+2 reconstruction strategy. This suggests that global structural compactness may influence how vibrational coupling is preserved during peptide fragmentation.

Several isotypes exhibit basis set-dependent structural expansion. βVI’s 4+2 demonstrates the largest increase in Rg, increasing from 25.6 Å under the minimal 3-21G basis set to 31.8 Å via 6-311+G(2df,2pd). βII’s 4+2 geometry similarly increases from 41.5 Å to approximately 46.1 Å. In contrast, βIII’s and βVI’s 2+4 structures shows minimal variation across basis sets, indicating comparatively rigid structural behavior. The consistently lower Rg values observed in βIII’s and βVI’s 2+4 geometries may reflect increased intramolecular stabilization arising from charged and polar residue interactions, particularly involving lysine and histidine residues.

Conversely, glutamate-rich sequences such as βI and βV appear to adopt more extended conformations, likely due to electrostatic repulsion among negatively charged sidechains. These results suggest that sequence-dependent structural compactness strongly influences vibrational behavior and may partially explain the distinct reconstruction preferences observed during Raman spectral analysis.

Additionally, the change in Rg primarily happens in the progression from 3-21G to 6-31G, and most isotypes exhibit a more minor change in the transition from the 6-31+G(d,p) to 6-311+G(2df,2pd) calculation. This may allow extensions of these E-hook structures, which are 18-24 amino acids at full length, to proceed by limiting the ballooning computational cost as basis set size increases.

### 3.3 Preferred Reconstructions Occupy Favorable Ramachandran Conformational Space

To determine whether Raman-derived structural preferences correspond to physically meaningful conformational differences, backbone dihedral angle distributions have been examined using Ramachandran analysis (**Figure 8**). The favored reconstruction strategies of each isotype consistently display more constrained φ and ψ angle distributions, indicating reduced conformational disorder. This relationship is most pronounced in βI and βV, which also demonstrates the largest Raman-derived differences between competing structural models. In both isotypes, the reconstruction that produces superior Raman agreement also generates noticeably tighter clustering of backbone φ and ψ angles, whereas the alternative reconstruction exhibits broader distributions and increased sampling across multiple dihedral regions. These structural differences parallel the Raman discrepancies observed between the competing reconstruction strategies, suggesting that conformational restriction is associated with improved spectral agreement in these isotypes. In contrast, βIII and βVI structures exhibit substantial overlap between the φ-ψ distributions generated by the two reconstruction strategies. Although preferred buildup methods for these two isotypes appear to be present, both of their 4+2 and 2+4 geometries show high agreement with experimental Raman spectra. This behavior is mimicked here in that both 2+4 and 4+2 geometries occupy similar backbone conformational space instead of sampling distinct conformations and backbone space.

**Figure 8.**
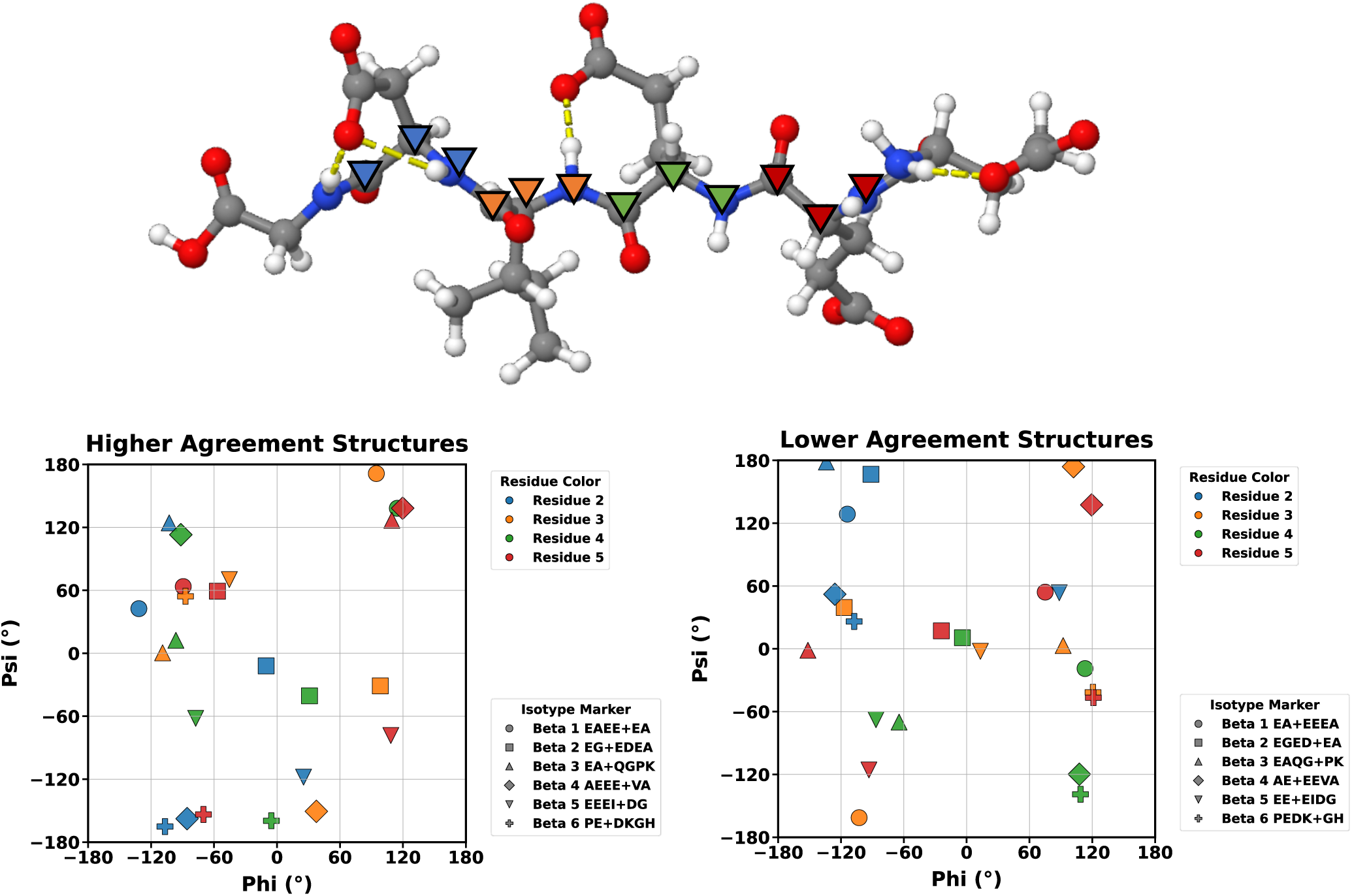
Ramachandran plots of higher agreement simulated geometries (left) and lower agreement simulated structures (right). Each structure is plotted by its isotype and residue number using unique marker shapes and marker colors, respectively. An example of the backbone labelling system is shown above, βV EE+EIDG.

The favored reconstruction methods that result in structures with better computational to experimental agreement tend to cluster within favorable areas of the Ramachandran chart. Several residues exist within the β-sheet/Extended area of the graph, lying within φ −150° to −90° and ψ 90° to 180°. The higher agreement dihedrals cluster around the other β extended/Polyproline II conformational basin which wraps across the ψ ±180° boundary. Only one dihedral resides in the right-handed α-helix portion, sitting between φ −90° to −30° and ψ - 70° to 0°. Interestingly, unfavored reconstruction methods’ dihedrals localize around the left-handed α-helix favored area of the chart, φ 30° to 90° and ψ 0° to 90°(59, 60). Otherwise, unfavored reconstruction strategies frequently exhibit broader conformational sampling and increased occupation of less favorable dihedral regions (any area outside of the listed areas of favored secondary structure), supporting the interpretation that Raman spectral agreement reflects physically meaningful structural differences rather than numerical fitting artifacts.

### 3.4 Residue Propensities for Hydrogen Bonding May Drive Conformational Differences

Intraresidue hydrogen bonds are hydrogen bonds wherein the acceptor and donor atoms are a part of the same residue. For example, a glutamic acid’s sidechain carboxylic acid group doubling back and interacting with its own backbone nitrogen or oxygen would be an intraresidue interaction.

Hydrogen bonding analysis has been performed to characterize local structural stabilization within each peptide fragment and determine whether hydrogen bonding patterns correlate with experimentally favored structural reconstructions (**Table 4**). Hydrogen bonding capacity varies substantially between E-hook isotypes. PE+DKGH are predicted to possess the largest total hydrogen bonding networks with seven hydrogen bonds, followed by EEEI+DG and EA+QGPK with six hydrogen bonds (**Figure 9**). EA+EEEA, EGED+EA, and PEDK+GH showcase the weakest overall hydrogen bonding capacities with only three total hydrogen bonds each.

**Figure 9.**
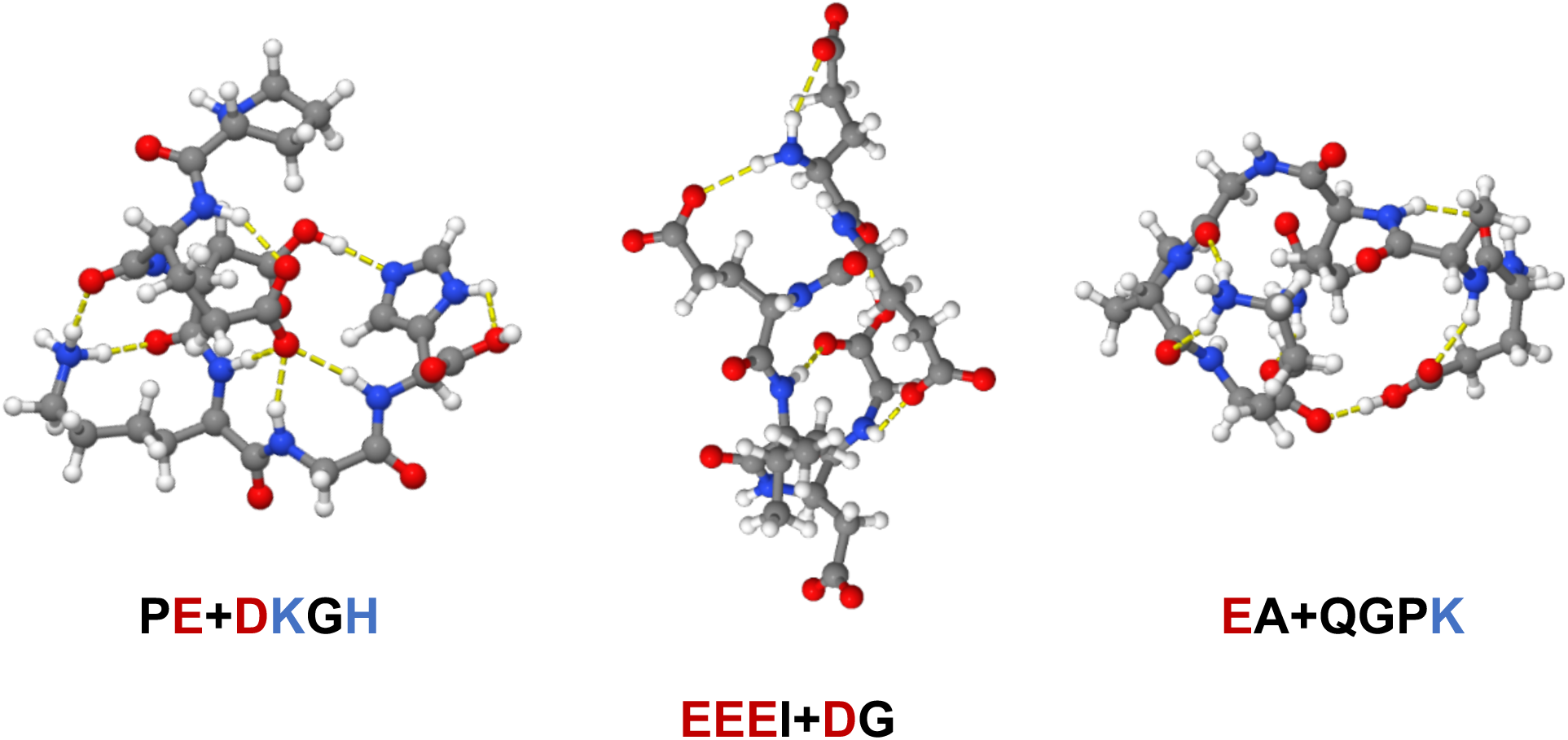
The three structures with the highest capacity for hydrogen bonding. Notice how the βVI (left) and βIII (right) structures obtain their high intramolecular interaction rate by adopting more closed conformers while the βV (middle) structure has landed in a more sheet-like orientation.

**Table 4.**
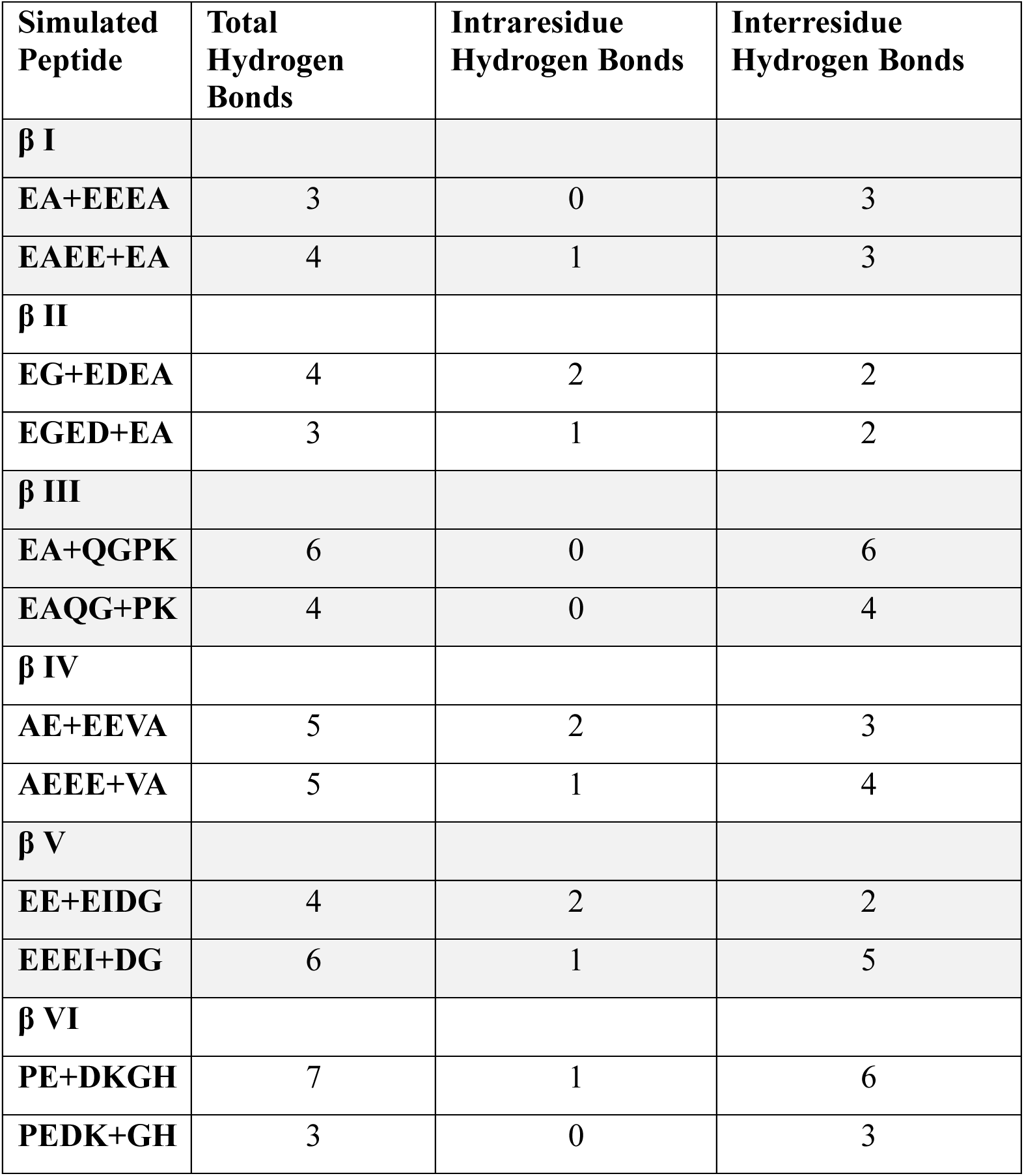
Hydrogen bonds for each simulated geometry wherein the hydrogen’s distance to the heavy acceptor atom is less than or equal to 2.5 Å and their bond angle is greater than or equal to 120°.

Across nearly all peptide reconstructions, interresidue bonding interactions (hydrogen bonds wherein the donor and acceptor atoms are a part of two different residues) substantially exceed intraresidue hydrogen bons (hydrogen bonds wherein the donor and acceptor atoms are a part of the same residue), indicating that structural stabilization primarily arises through cooperative residue-residue interactions rather than isolated local stabilization. For example, EAEE+EA forms six total hydrogen bonds but only one intraresidue hydrogen bond, suggesting that distributed intermolecular interactions dominate structural organization.

Residue-level analysis demonstrates strong dependence on amino acid identity (**Table 5**). Glutamate contributes the largest total hydrogen bonding contribution with 54 total hydrogen bonds and 1.4 interresidue bonds per residue across all 6-311+G(2df,2pd) basis set hexamer structures, reflecting its dominant structural role within glutamate-rich E-hook sequences. Lysine exhibits the strongest per-residue hydrogen bonding capacity with 2.8 total hydrogen bonds per residue and each being an interresidue bond, pointing to a strong local stabilizing potential. Glutamine residues similarly demonstrate strong intermolecular stabilization, averaging 2.0 total hydrogen bonds per residue and each being an interresidue interaction.

**Table 5.**
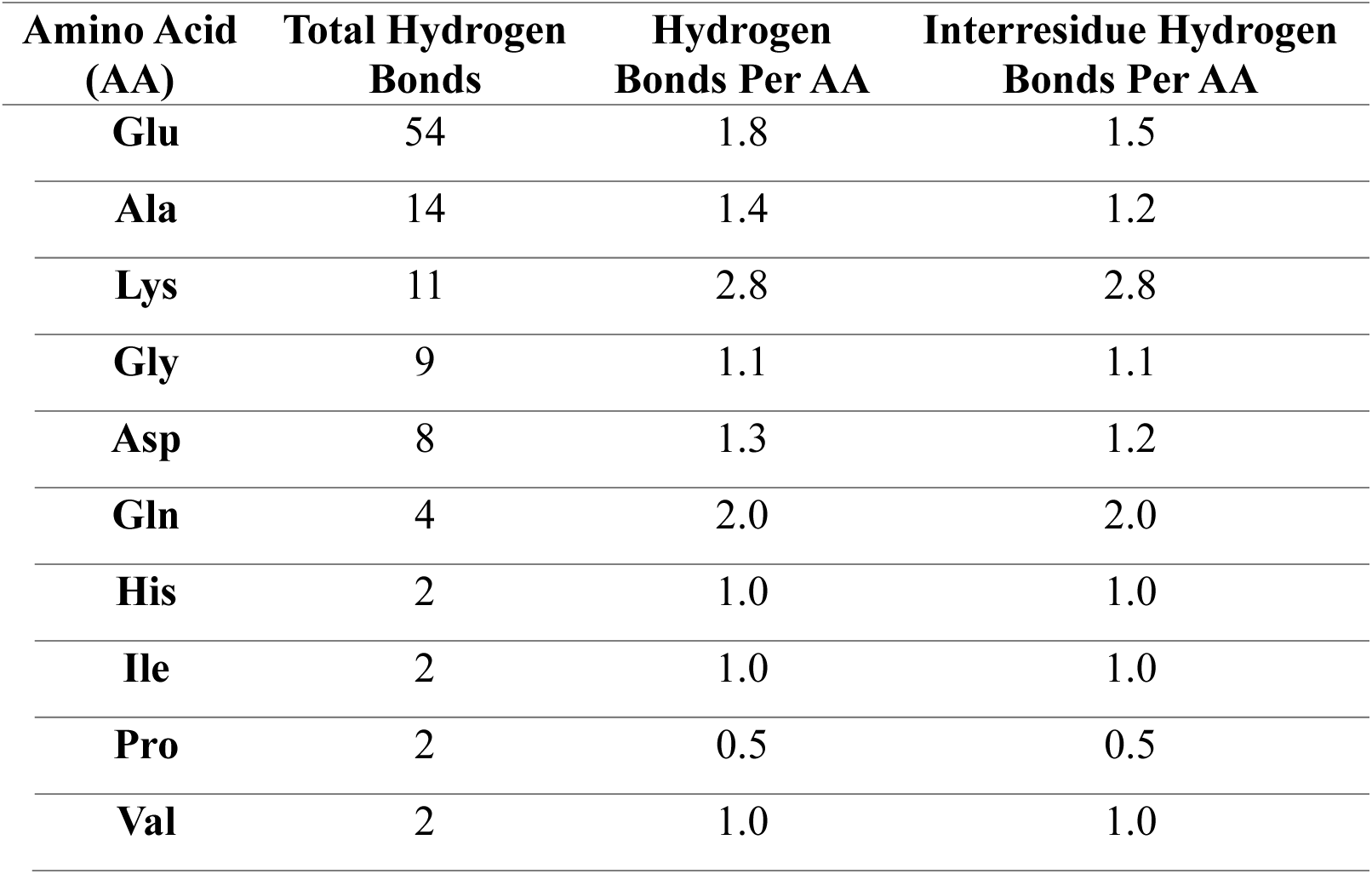
Hydrogen bonds formed by each amino acid throughout all simulation steps towards hexamer construction.

| **Amino Acid (AA)** | **Total Hydrogen Bonds** | **Hydrogen Bonds Per AA** | **Interresidue Hydrogen Bonds Per AA** |
| --- | --- | --- | --- |
| **Glu** | 54 | 1.8 | 1.5 |
| **Ala** | 14 | 1.4 | 1.2 |
| **Lys** | 11 | 2.8 | 2.8 |
| **Gly** | 9 | 1.1 | 1.1 |
| **Asp** | 8 | 1.3 | 1.2 |
| **Gln** | 4 | 2.0 | 2.0 |
| **His** | 2 | 1.0 | 1.0 |
| **Ile** | 2 | 1.0 | 1.0 |
| **Pro** | 2 | 0.5 | 0.5 |
| **Val** | 2 | 1.0 | 1.0 |

Differences in residue-specific hydrogen bonding behavior likely contribute to the observed isotype-dependent fragmentation preferences. Peptides rich in strongly interacting charged and polar residues may preserve structural integrity differently depending on fragmentation boundaries, explaining why preferred reconstruction strategy varies across

## 4. Conclusion

The results herein support a model in which β-tubulin E-hooks behave as electrostatically structured yet conformationally flexible peptides, capable of transient structural organization depending on sequence composition and interaction partners. Raman spectroscopy appears to provide an effective experimental framework for evaluating and differentiating computationally reconstructed β-tubulin E-hook peptides. By comparing experimentally obtained Raman spectra against quantum chemically simulated spectra, structural models have been assessed according to their ability to reproduce observed vibrational frequencies and overall spectral behavior.

Rather than supporting a single universally preferred fragmentation strategy, the present results indicate that local amino acid composition determines how structural interactions and vibrational coupling are preserved during peptide reconstruction. β1, β4, and β5 favor 4+2 fragmentation strategies, while β3, β2, β6 favor 2+4 fragmentation strategies. Ramachandran analysis indicates that experimentally favored structures generally occupy more conformationally-constrained regions of backbone conformational space, while hydrogen bonding analysis reveals substantial sequence-dependent differences in local stabilization behavior driven primarily by polar and charged residues.

The level of the quantum chemical theory employed also influences the predicted peptide geometry, although the magnitude of this effect is sequence- and reconstruction-dependent. Across the basis sets examined, changes in structural compactness are minimal for some E-hook structures, such as EA+QGPK (16.6-17.8 Å), but considerably larger for others, such as PEDK+GH (25.6-31.8 Å). The pronounced differences in compactness observed for the 2+4 reconstructions for the positively charged isotypes suggest that electrostatic interactions between acidic and basic residues influence E-hook conformational organization. In the context of the microtubule, these electrostatic interactions may influence how β-tubulin E-hooks engage positively charged regions of motor proteins, including kinesin and other microtubule-associated proteins.

Other, recent studies have shown that α-tubulin E-hooks are largely sequestered against the microtubule through interactions with basic residues in the tubulin core and become exposed only in the presence of specific MAPs(21). In contrast, β-tubulin E-hooks are expected to access the surrounding cytoplasmic environment to a greater extent, thereby increasing their opportunity to interact with MAPs, motor proteins, and other binding partners. Understanding how the conformations of β-tubulin E-hooks vary with isotype-specific compositions therefore provides insight into the molecular basis of the tubulin code and how subtle sequence differences may give rise to distinct functional behaviors.

## Author Contributions

A.C.B. was involved in all aspects of the work, including performing experiments, calculations, data analysis, and preparation of the manuscript. A.C.B., N.A.K., and C.R.B. performed the spectroscopic experiments and analysis. A.C.B., M.K.B., N.I.H., R.C.F., and D.N.R. designed the research and computational work and performed data analysis. A.B., N.I.H., R.C.F., and D.N.R. prepared the manuscript. All authors have read and agreed to the published version of the manuscript.

## Acknowledgements

The authors would like to thank the Mississippi Center for Supercomputing Research for lending resources during this study.

