## Supplemental Information for "Tubulin E-hook Hexamers Reveal Charge Dependent Compaction and Transient Secondary Structure Signatures"

**Table of Contents:**

1. Table S1: EA 6-311+G(*2df,2pd*) Cartesian Coordinates
2. Table S2: EE 6-311+G(*2df,2pd*) Cartesian Coordinates
3. Table S3: EG 6-311+G(*2df,2pd*) Cartesian Coordinates
4. Table S4: ED 6-311+G(*2df,2pd*) Cartesian Coordinates
5. Table S5: QG 6-311+G(*2df,2pd*) Cartesian Coordinates
6. Table S6: PK 6-311+G(*2df,2pd*) Cartesian Coordinates
7. Table S7: AE 6-311+G(*2df,2pd*) Cartesian Coordinates
8. Table S8: VA 6-311+G(*2df,2pd*) Cartesian Coordinates
9. Table S9: EI 6-311+G(*2df,2pd*) Cartesian Coordinates
10. Table S10: DG 6-311+G(*2df,2pd*) Cartesian Coordinates
11. Table S11: PE 6-311+G(*2df,2pd*) Cartesian Coordinates
12. Table S12: DK 6-311+G(*2df,2pd*) Cartesian Coordinates
13. Table S13: GH 6-311+G(*2df,2pd*) Cartesian Coordinates
14. Table S14: EAEE 6-311+G(*2df,2pd*) Cartesian Coordinates
15. Table S15: EEEA 6-311+G(*2df,2pd*) Cartesian Coordinates
16. Table S16: EGED 6-311+G(*2df,2pd*) Cartesian Coordinates
17. Table S17: EDEA 6-311+G(*2df,2pd*) Cartesian Coordinates
18. Table S18: EAQG 6-311+G(*2df,2pd*) Cartesian Coordinates
19. Table S19: QGPK 6-311+G(*2df,2pd*) Cartesian Coordinates
20. Table S20: AEEE 6-311+G(*2df,2pd*) Cartesian Coordinates
21. Table S21: EEVA 6-311+G(*2df,2pd*) Cartesian Coordinates
22. Table S22: EEEI 6-311+G(*2df,2pd*) Cartesian Coordinates
23. Table S23: EIDG 6-311+G(*2df,2pd*) Cartesian Coordinates
24. Table S24: PEDK 6-311+G(*2df,2pd*) Cartesian Coordinates
25. Table S25: DKGH 6-311+G(*2df,2pd*) Cartesian Coordinates
26. Table S26: EAEE+EA 6-311+G(*2df,2pd*) Cartesian Coordinates
27. Table S27: EA+EEEA 6-311+G(*2df,2pd*) Cartesian Coordinates
28. Table S28: EGED+EA 6-311+G(*2df,2pd*) Cartesian Coordinates
29. Table S29: EG+EDEA 6-311+G(*2df,2pd*) Cartesian Coordinates
30. Table S30: EAQG+PK 6-311+G(*2df,2pd*) Cartesian Coordinates
31. Table S31: EA+QGPK 6-311+G(*2df,2pd*) Cartesian Coordinates
32. Table S32: AEEE+VA 6-311+G(*2df,2pd*) Cartesian Coordinates
33. Table S33: AE+EEVA 6-311+G(*2df,2pd*) Cartesian Coordinates
34. Table S34: EEEI+DG 6-311+G(*2df,2pd*) Cartesian Coordinates
35. Table S35: EE+EIDG 6-311+G(*2df,2pd*) Cartesian Coordinates
36. Table S36: PEDK+GH 6-311+G(*2df,2pd*) Cartesian Coordinates
37. Table S37: PE+DKGH 6-311+G(*2df,2pd*) Cartesian Coordinates
38. Figure S1: Structures of βI Buildup
39. Figure S2: Structures of βII Buildup
40. Figure S3: Structures of βIV Buildup
41. Figure S4: Structures of βV Buildup
42. Figure S5: Structures of βVI Buildup
43. Dynamic Time Warping Analysis Code

1. **Table S1:** Cartesian Coordinates for the low-energy geometry of the **EA** dipeptide calculated using the B3LYP method and the 6-311+G(*2df,2pd*) level of theory.

| **Standard Orientation** | | | | | |
| --- | --- | --- | --- | --- | --- |
| Center Number | Atomic Number | Atomic Type | Coordinates (Angstroms) | | |
|  |  |  | X | Y | Z |
| 1 | 6 | 0 | -1.672265 | 1.534958 | -0.184027 |
| 2 | 7 | 0 | -2.106169 | 2.132056 | 1.081917 |
| 3 | 1 | 0 | -2.842953 | 2.806596 | 0.925962 |
| 4 | 1 | 0 | -1.313245 | 2.610039 | 1.497326 |
| 5 | 1 | 0 | -1.713214 | 2.271015 | -1.004739 |
| 6 | 6 | 0 | -0.154902 | 1.254024 | -0.065274 |
| 7 | 8 | 0 | 0.537720 | 1.969371 | 0.652477 |
| 8 | 7 | 0 | 0.356305 | 0.273272 | -0.835901 |
| 9 | 6 | 0 | 1.768476 | -0.096429 | -0.836706 |
| 10 | 1 | 0 | 1.801957 | -1.035156 | -1.396550 |
| 11 | 6 | 0 | 2.354139 | -0.458224 | 0.553591 |
| 12 | 8 | 0 | 3.477934 | -0.116002 | 0.851205 |
| 13 | 8 | 0 | 1.638440 | -1.225460 | 1.351496 |
| 14 | 1 | 0 | 0.736821 | -1.559993 | 1.035473 |
| 15 | 6 | 0 | 2.634387 | 0.937577 | -1.554595 |
| 16 | 1 | 0 | 2.252192 | 1.094486 | -2.564470 |
| 17 | 1 | 0 | 3.665522 | 0.595606 | -1.610094 |
| 18 | 1 | 0 | 2.618778 | 1.882392 | -1.015958 |
| 19 | 1 | 0 | -0.269425 | -0.396477 | -1.276595 |
| 20 | 6 | 0 | -2.615709 | 0.387207 | -0.598425 |
| 21 | 6 | 0 | -2.598853 | -0.879151 | 0.270007 |
| 22 | 1 | 0 | -3.522313 | -1.435250 | 0.084665 |
| 23 | 1 | 0 | -2.569293 | -0.622738 | 1.327185 |
| 24 | 6 | 0 | -1.423575 | -1.818622 | -0.079764 |
| 25 | 8 | 0 | -1.239370 | -2.056253 | -1.297697 |
| 26 | 8 | 0 | -0.702653 | -2.213611 | 0.876006 |
| 27 | 1 | 0 | -2.421476 | 0.099347 | -1.633694 |
| 28 | 1 | 0 | -3.621063 | 0.820427 | -0.587375 |

2. **Table S2:** Cartesian Coordinates for the low-energy geometry of the **EE** dipeptide calculated using the B3LYP method and the 6-311+G(*2df,2pd*) level of theory.

| Center Number | Atomic Number | Atomic Type | Coordinates (Angstroms) | | |
| --- | --- | --- | --- | --- | --- |
|  |  |  | X | Y | Z |
| 1 | 6 | 0 | 2.165583 | 0.532504 | 0.072056 |
| 2 | 6 | 0 | 2.751942 | -0.522109 | -0.886738 |
| 3 | 6 | 0 | 4.267913 | -0.433359 | -1.116486 |
| 4 | 6 | 0 | 5.253053 | -0.885267 | 0.008081 |
| 5 | 8 | 0 | 6.462177 | -0.792742 | -0.292660 |
| 6 | 8 | 0 | 4.759254 | -1.304437 | 1.089960 |
| 7 | 1 | 0 | 4.524627 | -1.038649 | -1.990897 |
| 8 | 1 | 0 | 4.541103 | 0.591114 | -1.388275 |
| 9 | 1 | 0 | 2.489926 | -1.511834 | -0.504330 |
| 10 | 1 | 0 | 2.250570 | -0.401618 | -1.849429 |
| 11 | 7 | 0 | 2.348349 | 0.125338 | 1.469772 |
| 12 | 1 | 0 | 2.479767 | 0.936191 | 2.061695 |
| 13 | 1 | 0 | 3.198723 | -0.453499 | 1.528517 |
| 14 | 6 | 0 | 0.685814 | 0.805700 | -0.260402 |
| 15 | 7 | 0 | -0.191248 | 0.347452 | 0.655357 |
| 16 | 6 | 0 | -1.635236 | 0.566321 | 0.642993 |
| 17 | 6 | 0 | -2.453884 | -0.537339 | -0.044753 |
| 18 | 6 | 0 | -3.940308 | -0.500848 | 0.315752 |
| 19 | 6 | 0 | -4.775659 | -1.768570 | -0.077931 |
| 20 | 8 | 0 | -6.017986 | -1.588810 | -0.084350 |
| 21 | 8 | 0 | -4.136325 | -2.819230 | -0.308202 |
| 22 | 1 | 0 | -4.047457 | -0.404257 | 1.404388 |
| 23 | 1 | 0 | -4.436132 | 0.368359 | -0.115312 |
| 24 | 1 | 0 | -2.050615 | -1.499878 | 0.270613 |
| 25 | 1 | 0 | -2.312912 | -0.484418 | -1.124362 |
| 26 | 6 | 0 | -1.914243 | 2.004932 | 0.181482 |
| 27 | 8 | 0 | -2.657860 | 2.101348 | -0.936089 |
| 28 | 1 | 0 | -2.738368 | 3.046926 | -1.119315 |
| 29 | 8 | 0 | -1.565631 | 2.984974 | 0.798185 |
| 30 | 1 | 0 | -1.931150 | 0.584706 | 1.696041 |
| 31 | 1 | 0 | 0.247942 | -0.096369 | 1.452827 |
| 32 | 8 | 0 | 0.355878 | 1.418268 | -1.272812 |
| 33 | 1 | 0 | 2.658345 | 1.486933 | -0.154655 |

3. **Table S3:** Cartesian Coordinates for the low-energy geometry of the **EG** dipeptide calculated using the B3LYP method and the 6-311+G(*2df,2pd*) level of theory.

| **Standard Orientation** | | | | | |
| --- | --- | --- | --- | --- | --- |
| Center Number | Atomic Number | Atomic Type | Coordinates (Angstroms) | | |
|  |  |  | X | Y | Z |
| 1 | 6 | 0 | -1.383917 | -1.357391 | 0.152249 |
| 2 | 7 | 0 | -1.383350 | -1.222938 | 1.617498 |
| 3 | 1 | 0 | -0.605629 | -1.733533 | 2.018541 |
| 4 | 1 | 0 | -1.264296 | -0.240351 | 1.847312 |
| 5 | 1 | 0 | -1.632567 | -2.393933 | -0.082424 |
| 6 | 6 | 0 | -2.478641 | -0.461506 | -0.454030 |
| 7 | 6 | 0 | -2.524446 | 1.052728 | -0.072236 |
| 8 | 8 | 0 | -3.568351 | 1.643532 | -0.373321 |
| 9 | 8 | 0 | -1.499116 | 1.545854 | 0.501414 |
| 10 | 1 | 0 | -2.428855 | -0.515192 | -1.545098 |
| 11 | 1 | 0 | -3.447260 | -0.876510 | -0.172819 |
| 12 | 6 | 0 | 0.011808 | -1.174933 | -0.477961 |
| 13 | 8 | 0 | 0.613837 | -2.124139 | -0.984811 |
| 14 | 7 | 0 | 0.518138 | 0.075308 | -0.401086 |
| 15 | 1 | 0 | -0.130882 | 0.799591 | -0.011882 |
| 16 | 6 | 0 | 1.798920 | 0.403170 | -0.944027 |
| 17 | 1 | 0 | 2.105014 | -0.389283 | -1.632420 |
| 18 | 1 | 0 | 1.759932 | 1.334850 | -1.511438 |
| 19 | 6 | 0 | 2.947286 | 0.560536 | 0.037984 |
| 20 | 8 | 0 | 4.020060 | 1.032757 | -0.257588 |
| 21 | 8 | 0 | 2.682031 | 0.109521 | 1.280104 |
| 22 | 1 | 0 | 3.487291 | 0.251950 | 1.797090 |

4. **Table S4:** Cartesian Coordinates for the low-energy geometry of the **ED** dipeptide calculated using the B3LYP method and the 6-311+G(*2df,2pd*) level of theory.

| **Standard Orientation** | | | | | |
| --- | --- | --- | --- | --- | --- |
| Center Number | Atomic Number | Atomic Type | Coordinates (Angstroms) | | |
|  |  |  | X | Y | Z |
| 1 | 6 | 0 | 2.204382 | 0.529791 | -0.320830 |
| 2 | 7 | 0 | 2.099997 | 1.200413 | 0.984306 |
| 3 | 1 | 0 | 1.837159 | 2.170957 | 0.851618 |
| 4 | 1 | 0 | 3.055537 | 1.187396 | 1.355390 |
| 5 | 1 | 0 | 2.693253 | 1.161268 | -1.073231 |
| 6 | 6 | 0 | 3.046252 | -0.749873 | -0.179322 |
| 7 | 6 | 0 | 4.536318 | -0.548845 | 0.256061 |
| 8 | 8 | 0 | 4.803291 | 0.480709 | 0.937072 |
| 9 | 8 | 0 | 5.320238 | -1.456699 | -0.089625 |
| 10 | 1 | 0 | 2.577854 | -1.395833 | 0.571148 |
| 11 | 1 | 0 | 3.035286 | -1.293486 | -1.121786 |
| 12 | 6 | 0 | 0.824474 | 0.212714 | -0.923033 |
| 13 | 8 | 0 | 0.693943 | -0.039924 | -2.116868 |
| 14 | 7 | 0 | -0.196581 | 0.237988 | -0.031909 |
| 15 | 1 | 0 | 0.077668 | 0.445446 | 0.919703 |
| 16 | 6 | 0 | -1.606879 | 0.010766 | -0.347741 |
| 17 | 1 | 0 | -1.665650 | -0.145141 | -1.420816 |
| 18 | 6 | 0 | -2.352127 | 1.314309 | -0.070036 |
| 19 | 8 | 0 | -2.610273 | 2.175618 | -0.874723 |
| 20 | 8 | 0 | -2.520122 | 1.530139 | 1.268001 |
| 21 | 1 | 0 | -2.985086 | 2.373728 | 1.333977 |
| 22 | 6 | 0 | -2.186577 | -1.187972 | 0.405299 |
| 23 | 6 | 0 | -3.725812 | -1.349278 | 0.239517 |
| 24 | 8 | 0 | -4.206774 | -2.445464 | 0.597838 |
| 25 | 8 | 0 | -4.321203 | -0.341355 | -0.220035 |
| 26 | 1 | 0 | -1.684890 | -2.096311 | 0.070423 |
| 27 | 1 | 0 | -1.978034 | -1.090703 | 1.474026 |

5. **Table S5:** Cartesian Coordinates for the low-energy geometry of the **QG** dipeptide calculated using the B3LYP method and the 6-311+G(*2df,2pd*) level of theory.

| **Standard Orientation** | | | | | |
| --- | --- | --- | --- | --- | --- |
| Center Number | Atomic Number | Atomic Type | Coordinates (Angstroms) | | |
|  |  |  | X | Y | Z |
| 1 | 6 | 0 | -0.412220 | 1.377533 | 0.530833 |
| 2 | 7 | 0 | -1.125142 | 2.585617 | 0.927396 |
| 3 | 1 | 0 | -0.625965 | 3.063662 | 1.667744 |
| 4 | 1 | 0 | -1.128985 | 3.227417 | 0.139967 |
| 5 | 1 | 0 | -0.120439 | 0.850912 | 1.443413 |
| 6 | 6 | 0 | -1.333426 | 0.453130 | -0.282921 |
| 7 | 6 | 0 | -2.538741 | -0.046126 | 0.517684 |
| 8 | 6 | 0 | -3.181000 | -1.257521 | -0.142401 |
| 9 | 7 | 0 | -4.532281 | -1.209516 | -0.290433 |
| 10 | 1 | 0 | -5.071507 | -0.407888 | -0.020352 |
| 11 | 1 | 0 | -5.000875 | -1.995950 | -0.707872 |
| 12 | 8 | 0 | -2.524559 | -2.222126 | -0.496290 |
| 13 | 1 | 0 | -2.207499 | -0.381952 | 1.504720 |
| 14 | 1 | 0 | -3.250685 | 0.760078 | 0.684303 |
| 15 | 1 | 0 | -0.780795 | -0.413334 | -0.645510 |
| 16 | 1 | 0 | -1.671121 | 0.997037 | -1.168666 |
| 17 | 6 | 0 | 0.845763 | 1.690056 | -0.303627 |
| 18 | 8 | 0 | 0.926648 | 2.723153 | -0.945597 |
| 19 | 7 | 0 | 1.847822 | 0.765676 | -0.322060 |
| 20 | 1 | 0 | 2.654002 | 0.987437 | -0.892786 |
| 21 | 6 | 0 | 1.912918 | -0.491546 | 0.377671 |
| 22 | 1 | 0 | 1.857671 | -0.364346 | 1.465059 |
| 23 | 1 | 0 | 1.102316 | -1.170064 | 0.091969 |
| 24 | 6 | 0 | 3.236485 | -1.178342 | 0.066892 |
| 25 | 8 | 0 | 4.101842 | -0.690103 | -0.598637 |
| 26 | 8 | 0 | 3.388098 | -2.403745 | 0.611109 |
| 27 | 1 | 0 | 2.596176 | -2.685993 | 1.084224 |

6. **Table S6:** Cartesian Coordinates for the low-energy geometry of the **PK** dipeptide calculated using the B3LYP method and the 6-311+G(*2df,2pd*) level of theory.

| **Standard Orientation** | | | | | |
| --- | --- | --- | --- | --- | --- |
| Center Number | Atomic Number | Atomic Type | Coordinates (Angstroms) | | |
|  |  |  | X | Y | Z |
| 1 | 6 | 0 | 2.062380 | -1.114524 | -0.475830 |
| 2 | 6 | 0 | 3.083226 | -1.613153 | 0.582565 |
| 3 | 6 | 0 | 4.031437 | -0.419337 | 0.757401 |
| 4 | 6 | 0 | 4.098730 | 0.168400 | -0.650293 |
| 5 | 7 | 0 | 2.711470 | 0.017755 | -1.148921 |
| 6 | 1 | 0 | 2.669068 | -0.051343 | -2.157354 |
| 7 | 1 | 0 | 4.812638 | -0.392621 | -1.264497 |
| 8 | 1 | 0 | 4.392595 | 1.218446 | -0.663567 |
| 9 | 1 | 0 | 3.611846 | 0.311363 | 1.453701 |
| 10 | 1 | 0 | 5.010563 | -0.712749 | 1.134720 |
| 11 | 1 | 0 | 3.625785 | -2.468140 | 0.175346 |
| 12 | 1 | 0 | 2.602997 | -1.937760 | 1.504508 |
| 13 | 1 | 0 | 1.805799 | -1.925356 | -1.164854 |
| 14 | 6 | 0 | 0.737287 | -0.710569 | 0.181739 |
| 15 | 8 | 0 | 0.031591 | -1.589975 | 0.724984 |
| 16 | 7 | 0 | 0.444452 | 0.582507 | 0.116979 |
| 17 | 1 | 0 | 1.140010 | 1.126393 | -0.403436 |
| 18 | 6 | 0 | -0.651814 | 1.310383 | 0.742019 |
| 19 | 6 | 0 | -2.026304 | 0.614795 | 0.686583 |
| 20 | 1 | 0 | -1.970579 | -0.236517 | 1.362410 |
| 21 | 1 | 0 | -2.752867 | 1.301358 | 1.123775 |
| 22 | 6 | 0 | -2.489850 | 0.192211 | -0.715417 |
| 23 | 1 | 0 | -2.868464 | 1.070389 | -1.242815 |
| 24 | 1 | 0 | -1.643096 | -0.165592 | -1.308270 |
| 25 | 6 | 0 | -3.590919 | -0.884163 | -0.703187 |
| 26 | 1 | 0 | -4.282308 | -0.718936 | 0.130826 |
| 27 | 6 | 0 | -3.104626 | -2.330523 | -0.696138 |
| 28 | 1 | 0 | -2.439681 | -2.524913 | -1.538223 |
| 29 | 1 | 0 | -3.946153 | -3.020485 | -0.758610 |
| 30 | 7 | 0 | -2.320376 | -2.701167 | 0.541647 |
| 31 | 1 | 0 | -2.865555 | -2.529050 | 1.387727 |
| 32 | 1 | 0 | -2.083863 | -3.694792 | 0.532671 |
| 33 | 1 | 0 | -1.389506 | -2.163730 | 0.616540 |
| 34 | 1 | 0 | -4.198142 | -0.793710 | -1.607390 |
| 35 | 1 | 0 | -0.419598 | 1.468636 | 1.802301 |
| 36 | 6 | 0 | -0.666243 | 2.705786 | 0.113788 |
| 37 | 8 | 0 | 0.109829 | 3.082762 | -0.722580 |
| 38 | 8 | 0 | -1.639047 | 3.467069 | 0.639715 |
| 39 | 1 | 0 | -1.579108 | 4.350758 | 0.240218 |

7. **Table S7:** Cartesian Coordinates for the low-energy geometry of the **AE** dipeptide calculated using the B3LYP method and the 6-311+G(*2df,2pd*) level of theory.

| **Standard Orientation** | | | | | |
| --- | --- | --- | --- | --- | --- |
| Center Number | Atomic Number | Atomic Type | Coordinates (Angstroms) | | |
|  |  |  | X | Y | Z |
| 1 | 6 | 0 | -2.466550 | -1.643044 | -0.197501 |
| 2 | 7 | 0 | -2.604576 | -2.528303 | 0.959797 |
| 3 | 1 | 0 | -2.883349 | -1.963758 | 1.757018 |
| 4 | 1 | 0 | -1.702845 | -2.923951 | 1.202161 |
| 5 | 1 | 0 | -1.963039 | -2.208586 | -0.985418 |
| 6 | 6 | 0 | -1.644623 | -0.384510 | 0.138121 |
| 7 | 8 | 0 | -1.863302 | 0.247797 | 1.162260 |
| 8 | 7 | 0 | -0.722030 | -0.019756 | -0.782174 |
| 9 | 1 | 0 | -0.489973 | -0.691597 | -1.494063 |
| 10 | 6 | 0 | 0.278871 | 1.031875 | -0.571089 |
| 11 | 6 | 0 | 1.519873 | 0.542986 | 0.177171 |
| 12 | 1 | 0 | 1.239209 | 0.265090 | 1.194664 |
| 13 | 6 | 0 | 2.208115 | -0.620354 | -0.534225 |
| 14 | 1 | 0 | 1.626137 | -1.541793 | -0.469455 |
| 15 | 1 | 0 | 2.316531 | -0.387512 | -1.602746 |
| 16 | 6 | 0 | 3.661370 | -0.938384 | -0.023930 |
| 17 | 8 | 0 | 4.013347 | -2.129712 | -0.166304 |
| 18 | 8 | 0 | 4.303564 | 0.035195 | 0.425536 |
| 19 | 1 | 0 | 2.233110 | 1.361378 | 0.267123 |
| 20 | 6 | 0 | -0.356860 | 2.286944 | 0.018286 |
| 21 | 8 | 0 | 0.105973 | 2.968352 | 0.890912 |
| 22 | 8 | 0 | -1.482099 | 2.649817 | -0.657185 |
| 23 | 1 | 0 | -1.809083 | 3.445715 | -0.216329 |
| 24 | 1 | 0 | 0.589432 | 1.338713 | -1.576425 |
| 25 | 6 | 0 | -3.850692 | -1.221748 | -0.695038 |
| 26 | 1 | 0 | -4.371717 | -0.666008 | 0.085635 |
| 27 | 1 | 0 | -4.437691 | -2.105185 | -0.943988 |
| 28 | 1 | 0 | -3.777379 | -0.580266 | -1.574050 |

8. **Table S8:** Cartesian Coordinates for the low-energy geometry of the **VA** dipeptide calculated using the B3LYP method and the 6-311+G(*2df,2pd*) level of theory.

| **Standard Orientation** | | | | | |
| --- | --- | --- | --- | --- | --- |
| Center Number | Atomic Number | Atomic Type | Coordinates (Angstroms) | | |
|  |  |  | X | Y | Z |
| 1 | 6 | 0 | -1.816384 | -0.399750 | 0.470839 |
| 2 | 7 | 0 | -2.548873 | -1.635168 | 0.757304 |
| 3 | 1 | 0 | -2.059779 | -2.194486 | 1.445514 |
| 4 | 1 | 0 | -2.600984 | -2.191436 | -0.089963 |
| 5 | 1 | 0 | -1.601488 | 0.085686 | 1.428246 |
| 6 | 6 | 0 | -0.498822 | -0.709402 | -0.254786 |
| 7 | 8 | 0 | -0.479198 | -1.367635 | -1.284518 |
| 8 | 7 | 0 | 0.635206 | -0.232082 | 0.325715 |
| 9 | 1 | 0 | 0.570906 | 0.317763 | 1.165037 |
| 10 | 6 | 0 | 1.947143 | -0.440684 | -0.265042 |
| 11 | 6 | 0 | 2.591775 | -1.766780 | 0.174695 |
| 12 | 1 | 0 | 2.733849 | -1.791027 | 1.255610 |
| 13 | 1 | 0 | 3.557191 | -1.897077 | -0.311308 |
| 14 | 1 | 0 | 1.938136 | -2.587320 | -0.113434 |
| 15 | 6 | 0 | 2.899726 | 0.704018 | 0.030296 |
| 16 | 8 | 0 | 2.523106 | 1.454493 | 1.095098 |
| 17 | 1 | 0 | 3.201881 | 2.132908 | 1.224376 |
| 18 | 8 | 0 | 3.912879 | 0.909709 | -0.579783 |
| 19 | 1 | 0 | 1.818620 | -0.468432 | -1.346435 |
| 20 | 6 | 0 | -2.689075 | 0.558352 | -0.371899 |
| 21 | 6 | 0 | -3.971223 | 0.929240 | 0.377087 |
| 22 | 1 | 0 | -4.623425 | 1.530296 | -0.258290 |
| 23 | 1 | 0 | -3.742779 | 1.521386 | 1.267622 |
| 24 | 1 | 0 | -4.515613 | 0.042647 | 0.693301 |
| 25 | 6 | 0 | -1.924208 | 1.816414 | -0.794559 |
| 26 | 1 | 0 | -1.583516 | 2.383012 | 0.076340 |
| 27 | 1 | 0 | -2.570951 | 2.473837 | -1.376399 |
| 28 | 1 | 0 | -1.054789 | 1.585004 | -1.409828 |
| 29 | 1 | 0 | -2.959479 | 0.007005 | -1.277683 |

9. **Table S9:** Cartesian Coordinates for the low-energy geometry of the **EI** dipeptide calculated using the B3LYP method and the 6-311+G(*2df,2pd*) level of theory.

| **Standard Orientation** | | | | | |
| --- | --- | --- | --- | --- | --- |
| Center Number | Atomic Number | Atomic Type | Coordinates (Angstroms) | | |
|  |  |  | X | Y | Z |
| 1 | 6 | 0 | -1.435883 | -1.114700 | 1.155853 |
| 2 | 7 | 0 | -1.630989 | -2.442053 | 1.743164 |
| 3 | 1 | 0 | -2.480643 | -2.457773 | 2.291201 |
| 4 | 1 | 0 | -0.844736 | -2.676799 | 2.336646 |
| 5 | 1 | 0 | -1.530560 | -0.311428 | 1.908334 |
| 6 | 6 | 0 | -2.505659 | -0.847254 | 0.085290 |
| 7 | 1 | 0 | -2.347626 | -1.531547 | -0.752221 |
| 8 | 1 | 0 | -3.470457 | -1.115735 | 0.522277 |
| 9 | 6 | 0 | -2.673690 | 0.597158 | -0.397496 |
| 10 | 1 | 0 | -2.315428 | 1.301640 | 0.361827 |
| 11 | 1 | 0 | -2.121042 | 0.815506 | -1.316724 |
| 12 | 6 | 0 | -4.166645 | 1.030478 | -0.685277 |
| 13 | 8 | 0 | -5.050640 | 0.383886 | -0.084210 |
| 14 | 8 | 0 | -4.275354 | 2.004163 | -1.459509 |
| 15 | 6 | 0 | 0.032182 | -1.030148 | 0.724134 |
| 16 | 8 | 0 | 0.918746 | -1.644711 | 1.309493 |
| 17 | 7 | 0 | 0.320733 | -0.199494 | -0.306384 |
| 18 | 1 | 0 | -0.441123 | 0.271230 | -0.769037 |
| 19 | 6 | 0 | 1.677502 | 0.040817 | -0.762601 |
| 20 | 6 | 0 | 2.528086 | 0.992850 | 0.121464 |
| 21 | 6 | 0 | 1.678483 | 2.140410 | 0.690275 |
| 22 | 6 | 0 | 2.422874 | 3.039583 | 1.678759 |
| 23 | 1 | 0 | 1.744247 | 3.776985 | 2.109865 |
| 24 | 1 | 0 | 3.242563 | 3.585734 | 1.210628 |
| 25 | 1 | 0 | 2.840403 | 2.454014 | 2.501089 |
| 26 | 1 | 0 | 0.807125 | 1.723610 | 1.193172 |
| 27 | 1 | 0 | 1.288630 | 2.749002 | -0.133117 |
| 28 | 6 | 0 | 3.718341 | 1.525838 | -0.687172 |
| 29 | 1 | 0 | 4.304351 | 0.715585 | -1.114149 |
| 30 | 1 | 0 | 3.372555 | 2.172333 | -1.499674 |
| 31 | 1 | 0 | 4.386041 | 2.116031 | -0.060177 |
| 32 | 1 | 0 | 2.908748 | 0.400983 | 0.956077 |
| 33 | 1 | 0 | 1.573232 | 0.503818 | -1.749453 |
| 34 | 6 | 0 | 2.401000 | -1.288650 | -1.025612 |
| 35 | 8 | 0 | 1.624970 | -2.140618 | -1.729790 |
| 36 | 1 | 0 | 2.133094 | -2.956760 | -1.833093 |
| 37 | 8 | 0 | 3.540137 | -1.548207 | -0.736815 |

10. **Table S10:** Cartesian Coordinates for the low-energy geometry of the **DG** dipeptide calculated using the B3LYP method and the 6-311+G(*2df,2pd*) level of theory.

| **Standard Orientation** | | | | | |
| --- | --- | --- | --- | --- | --- |
| Center Number | Atomic Number | Atomic Type | Coordinates (Angstroms) | | |
|  |  |  | X | Y | Z |
| 1 | 6 | 0 | 1.156288 | -0.127405 | -0.605401 |
| 2 | 1 | 0 | 1.481796 | -0.036260 | -1.647906 |
| 3 | 7 | 0 | 1.022830 | -1.523751 | -0.167823 |
| 4 | 1 | 0 | 0.666650 | -2.093177 | -0.927242 |
| 5 | 1 | 0 | 1.989808 | -1.824381 | 0.021085 |
| 6 | 6 | 0 | -0.178814 | 0.609175 | -0.520360 |
| 7 | 8 | 0 | -0.422064 | 1.644784 | -1.128073 |
| 8 | 7 | 0 | -1.069319 | 0.028645 | 0.335090 |
| 9 | 1 | 0 | -0.764098 | -0.859323 | 0.717705 |
| 10 | 6 | 0 | -2.346727 | 0.598083 | 0.605778 |
| 11 | 1 | 0 | -2.491678 | 0.818991 | 1.668808 |
| 12 | 1 | 0 | -2.402599 | 1.552298 | 0.075740 |
| 13 | 6 | 0 | -3.518181 | -0.252266 | 0.161245 |
| 14 | 8 | 0 | -4.695615 | 0.344861 | 0.497098 |
| 15 | 1 | 0 | -5.399077 | -0.241476 | 0.186162 |
| 16 | 8 | 0 | -3.479092 | -1.314365 | -0.397869 |
| 17 | 6 | 0 | 2.187337 | 0.603753 | 0.277894 |
| 18 | 6 | 0 | 3.639935 | 0.013167 | 0.267725 |
| 19 | 8 | 0 | 4.544557 | 0.838472 | 0.482986 |
| 20 | 8 | 0 | 3.742985 | -1.231244 | 0.085049 |
| 21 | 1 | 0 | 1.841202 | 0.572410 | 1.316800 |
| 22 | 1 | 0 | 2.238218 | 1.649572 | -0.016821 |

11. **Table S12:** Cartesian Coordinates for the low-energy geometry of the **PE** dipeptide calculated using the B3LYP method and the 6-311+G(*2df,2pd*) level of theory.

| **Standard Orientation** | | | | | |
| --- | --- | --- | --- | --- | --- |
| Center Number | Atomic Number | Atomic Type | Coordinates (Angstroms) | | |
|  |  |  | X | Y | Z |
| 1 | 6 | 0 | -4.452961 | -0.235661 | 0.055020 |
| 2 | 7 | 0 | -3.413280 | 0.775920 | -0.066789 |
| 3 | 1 | 0 | -3.633044 | 1.560538 | -0.664002 |
| 4 | 6 | 0 | -2.223543 | 0.052609 | -0.499462 |
| 5 | 1 | 0 | -2.344208 | -0.344262 | -1.526177 |
| 6 | 6 | 0 | -2.206867 | -1.138787 | 0.471442 |
| 7 | 6 | 0 | -3.712379 | -1.420918 | 0.720527 |
| 8 | 1 | 0 | -4.026977 | -2.371094 | 0.288520 |
| 9 | 1 | 0 | -3.926844 | -1.464363 | 1.788095 |
| 10 | 1 | 0 | -1.660484 | -1.992271 | 0.078404 |
| 11 | 1 | 0 | -1.715565 | -0.839724 | 1.396115 |
| 12 | 6 | 0 | -0.997507 | 0.960923 | -0.563055 |
| 13 | 8 | 0 | -1.124302 | 2.172121 | -0.754366 |
| 14 | 7 | 0 | 0.186764 | 0.336640 | -0.471762 |
| 15 | 1 | 0 | 0.298972 | -0.708445 | -0.384848 |
| 16 | 6 | 0 | 1.422415 | 1.051541 | -0.669664 |
| 17 | 6 | 0 | 2.572092 | 0.080166 | -1.022506 |
| 18 | 6 | 0 | 2.930517 | -1.002414 | 0.011582 |
| 19 | 6 | 0 | 2.092567 | -2.317314 | -0.006979 |
| 20 | 8 | 0 | 0.873195 | -2.214824 | -0.350388 |
| 21 | 8 | 0 | 2.689893 | -3.347945 | 0.334633 |
| 22 | 1 | 0 | 2.865344 | -0.597866 | 1.026771 |
| 23 | 1 | 0 | 3.971481 | -1.294444 | -0.126306 |
| 24 | 1 | 0 | 3.449357 | 0.700887 | -1.213838 |
| 25 | 1 | 0 | 2.304482 | -0.397388 | -1.967077 |
| 26 | 1 | 0 | 1.306568 | 1.758090 | -1.494937 |
| 27 | 6 | 0 | 1.837423 | 1.938415 | 0.502071 |
| 28 | 8 | 0 | 1.242683 | 1.634215 | 1.673291 |
| 29 | 1 | 0 | 1.589531 | 2.262104 | 2.322399 |
| 30 | 8 | 0 | 2.653740 | 2.825098 | 0.412464 |
| 31 | 1 | 0 | -4.854834 | -0.554609 | -0.922036 |
| 32 | 1 | 0 | -5.290366 | 0.134252 | 0.649842 |

12. **Table S11:** Cartesian Coordinates for the low-energy geometry of the **DK** dipeptide calculated using the B3LYP method and the 6-311+G(*2df,2pd*) level of theory.

| **Standard Orientation** | | | | | |
| --- | --- | --- | --- | --- | --- |
| Center Number | Atomic Number | Atomic Type | Coordinates (Angstroms) | | |
|  |  |  | X | Y | Z |
| 1 | 6 | 0 | 0.530434 | 1.225753 | -0.657763 |
| 2 | 7 | 0 | -0.669302 | 0.421649 | -0.463102 |
| 3 | 6 | 0 | -0.756831 | -0.674071 | 0.180057 |
| 4 | 6 | 0 | -2.109766 | -1.369940 | 0.329601 |
| 5 | 7 | 0 | -1.916949 | -2.810545 | 0.123206 |
| 6 | 1 | 0 | -1.203148 | -3.167250 | 0.744287 |
| 7 | 1 | 0 | -2.777404 | -3.306176 | 0.318010 |
| 8 | 1 | 0 | -2.436954 | -1.136530 | 1.355291 |
| 9 | 6 | 0 | -3.197148 | -0.897160 | -0.650820 |
| 10 | 6 | 0 | -3.899307 | 0.416371 | -0.313267 |
| 11 | 8 | 0 | -5.074183 | 0.469065 | -0.050019 |
| 12 | 8 | 0 | -3.135022 | 1.510385 | -0.352701 |
| 13 | 1 | 0 | -3.978753 | -1.652966 | -0.669998 |
| 14 | 1 | 0 | -2.768990 | -0.835504 | -1.652414 |
| 15 | 8 | 0 | 0.180720 | -1.368862 | 0.830463 |
| 16 | 1 | 0 | -2.192709 | 1.228064 | -0.513197 |
| 17 | 6 | 0 | 1.734615 | 0.441755 | -1.280844 |
| 18 | 1 | 0 | 1.733466 | 0.635894 | -2.353422 |
| 19 | 1 | 0 | 1.514339 | -0.617266 | -1.181649 |
| 20 | 6 | 0 | 3.126250 | 0.736386 | -0.705239 |
| 21 | 1 | 0 | 3.531047 | 1.649802 | -1.141920 |
| 22 | 6 | 0 | 4.126555 | -0.409206 | -0.898762 |
| 23 | 1 | 0 | 4.352358 | -0.553661 | -1.958101 |
| 24 | 6 | 0 | 3.666480 | -1.754704 | -0.323023 |
| 25 | 1 | 0 | 2.955477 | -2.233094 | -0.998172 |
| 26 | 7 | 0 | 2.982889 | -1.592703 | 0.976877 |
| 27 | 1 | 0 | 3.576643 | -1.098804 | 1.634598 |
| 28 | 1 | 0 | 2.797835 | -2.501225 | 1.387179 |
| 29 | 1 | 0 | 1.118338 | -1.060695 | 0.819590 |
| 30 | 1 | 0 | 4.529850 | -2.424462 | -0.257877 |
| 31 | 1 | 0 | 5.069534 | -0.119635 | -0.426755 |
| 32 | 1 | 0 | 3.052061 | 0.934882 | 0.363579 |
| 33 | 6 | 0 | 0.919392 | 2.047317 | 0.573624 |
| 34 | 8 | 0 | 1.426088 | 3.135523 | 0.507269 |
| 35 | 8 | 0 | 0.647075 | 1.441019 | 1.748328 |
| 36 | 1 | 0 | 0.898657 | 2.056678 | 2.452563 |
| 37 | 1 | 0 | 0.250430 | 2.001097 | -1.368556 |

13. **Table S13:** Cartesian Coordinates for the low-energy geometry of the **GH** dipeptide calculated using the B3LYP method and the 6-311+G(*2df,2pd*) level of theory.

| **Standard Orientation** | | | | | |
| --- | --- | --- | --- | --- | --- |
| Center Number | Atomic Number | Atomic Type | Coordinates (Angstroms) | | |
|  |  |  | X | Y | Z |
| 1 | 6 | 0 | 3.353503 | -0.090710 | -0.402754 |
| 2 | 7 | 0 | 4.376717 | 0.780769 | 0.109195 |
| 3 | 1 | 0 | 4.146179 | 1.755717 | -0.028663 |
| 4 | 1 | 0 | 4.526631 | 0.644091 | 1.099945 |
| 5 | 1 | 0 | 3.645352 | -1.131465 | -0.235350 |
| 6 | 1 | 0 | 3.283072 | 0.023948 | -1.488797 |
| 7 | 6 | 0 | 1.950325 | 0.106252 | 0.177185 |
| 8 | 8 | 0 | 1.659979 | 0.965433 | 0.988214 |
| 9 | 7 | 0 | 0.981069 | -0.750767 | -0.299851 |
| 10 | 1 | 0 | 1.222264 | -1.579688 | -0.825686 |
| 11 | 6 | 0 | -0.304679 | -0.763583 | 0.344073 |
| 12 | 1 | 0 | -0.181503 | -0.761545 | 1.433924 |
| 13 | 6 | 0 | -1.119074 | 0.452968 | -0.006876 |
| 14 | 6 | 0 | -1.143764 | 1.214964 | -1.133137 |
| 15 | 7 | 0 | -2.050701 | 2.230457 | -0.908382 |
| 16 | 6 | 0 | -2.573065 | 2.109406 | 0.307872 |
| 17 | 7 | 0 | -2.020023 | 1.035210 | 0.864524 |
| 18 | 1 | 0 | -2.229293 | 0.701018 | 1.794585 |
| 19 | 1 | 0 | -3.302794 | 2.758271 | 0.758154 |
| 20 | 1 | 0 | -2.280414 | 2.967507 | -1.559083 |
| 21 | 1 | 0 | -0.584543 | 1.131796 | -2.045784 |
| 22 | 6 | 0 | -1.038875 | -2.062891 | -0.017094 |
| 23 | 8 | 0 | -0.577654 | -2.923259 | -0.706795 |
| 24 | 8 | 0 | -2.247400 | -2.097081 | 0.565283 |
| 25 | 1 | 0 | -2.680017 | -2.938502 | 0.349130 |

14. **Table S14:** Cartesian Coordinates for the low-energy geometry of the **EAEE** tetrapeptide calculated using the B3LYP method and the 6-311+G(*2df,2pd*) level of theory.

| **Standard Orientation** | | | | | |
| --- | --- | --- | --- | --- | --- |
| Center Number | Atomic Number | Atomic Type | Coordinates (Angstroms) | | |
|  |  |  | X | Y | Z |
| 1 | 6 | 0 | 2.669508 | -2.764386 | -0.806914 |
| 2 | 7 | 0 | 2.005286 | -3.610446 | 0.195716 |
| 3 | 1 | 0 | 1.973600 | -4.570776 | -0.123867 |
| 4 | 1 | 0 | 1.048016 | -3.291344 | 0.301356 |
| 5 | 1 | 0 | 2.330754 | -3.021048 | -1.824464 |
| 6 | 6 | 0 | 2.111601 | -1.334718 | -0.646360 |
| 7 | 8 | 0 | 0.880326 | -1.204981 | -0.692265 |
| 8 | 7 | 0 | 2.964714 | -0.316524 | -0.527176 |
| 9 | 6 | 0 | 2.516409 | 1.077826 | -0.525236 |
| 10 | 1 | 0 | 3.356264 | 1.641180 | -0.113272 |
| 11 | 6 | 0 | 1.372292 | 1.290655 | 0.498970 |
| 12 | 8 | 0 | 1.487638 | 0.864683 | 1.640222 |
| 13 | 7 | 0 | 0.284512 | 1.977058 | 0.066506 |
| 14 | 6 | 0 | -0.810744 | 2.304497 | 0.963197 |
| 15 | 6 | 0 | -1.521112 | 3.592062 | 0.507018 |
| 16 | 6 | 0 | -0.659233 | 4.863863 | 0.499346 |
| 17 | 6 | 0 | 0.102449 | 5.201382 | -0.817741 |
| 18 | 8 | 0 | 0.447024 | 6.395177 | -0.952518 |
| 19 | 8 | 0 | 0.310537 | 4.246816 | -1.617622 |
| 20 | 1 | 0 | -1.281506 | 5.731565 | 0.728280 |
| 21 | 1 | 0 | 0.090579 | 4.818495 | 1.296937 |
| 22 | 1 | 0 | -1.929182 | 3.430759 | -0.493688 |
| 23 | 1 | 0 | -2.367891 | 3.720636 | 1.180343 |
| 24 | 6 | 0 | -1.867192 | 1.188990 | 1.138325 |
| 25 | 7 | 0 | -1.712503 | 0.056762 | 0.426633 |
| 26 | 6 | 0 | -2.629458 | -1.084164 | 0.504818 |
| 27 | 6 | 0 | -3.779118 | -1.050316 | -0.522017 |
| 28 | 6 | 0 | -4.432389 | -2.420075 | -0.731369 |
| 29 | 6 | 0 | -5.638821 | -2.492241 | -1.727317 |
| 30 | 8 | 0 | -6.144161 | -3.639360 | -1.842046 |
| 31 | 8 | 0 | -5.982025 | -1.434805 | -2.301511 |
| 32 | 1 | 0 | -3.685069 | -3.134858 | -1.093187 |
| 33 | 1 | 0 | -4.793963 | -2.824362 | 0.216326 |
| 34 | 1 | 0 | -3.360715 | -0.703488 | -1.467282 |
| 35 | 1 | 0 | -4.523855 | -0.312650 | -0.229601 |
| 36 | 6 | 0 | -3.063608 | -1.310401 | 1.961473 |
| 37 | 8 | 0 | -4.395926 | -1.208427 | 2.173856 |
| 38 | 1 | 0 | -4.512451 | -1.360326 | 3.120722 |
| 39 | 8 | 0 | -2.315040 | -1.642218 | 2.848623 |
| 40 | 1 | 0 | -2.004478 | -1.953026 | 0.286971 |
| 41 | 1 | 0 | -0.839933 | -0.096779 | -0.065968 |
| 42 | 8 | 0 | -2.809944 | 1.367917 | 1.908279 |
| 43 | 1 | 0 | -0.408226 | 2.452256 | 1.969146 |
| 44 | 1 | 0 | 0.342898 | 2.582484 | -0.760529 |
| 45 | 6 | 0 | 2.253952 | 1.565485 | -1.953124 |
| 46 | 1 | 0 | 3.157629 | 1.414730 | -2.545892 |
| 47 | 1 | 0 | 1.990734 | 2.622650 | -1.984255 |
| 48 | 1 | 0 | 1.444269 | 0.996528 | -2.409393 |
| 49 | 1 | 0 | 3.990316 | -0.468707 | -0.445811 |
| 50 | 6 | 0 | 4.188374 | -2.970999 | -0.798479 |
| 51 | 6 | 0 | 4.951804 | -2.686566 | 0.513816 |
| 52 | 1 | 0 | 5.373165 | -3.602103 | 0.927236 |
| 53 | 1 | 0 | 4.259343 | -2.294831 | 1.262412 |
| 54 | 6 | 0 | 6.086572 | -1.644559 | 0.333565 |
| 55 | 8 | 0 | 5.740661 | -0.583278 | -0.266090 |
| 56 | 8 | 0 | 7.215789 | -1.925442 | 0.785554 |
| 57 | 1 | 0 | 4.611677 | -2.358777 | -1.597177 |
| 58 | 1 | 0 | 4.357251 | -4.012828 | -1.094781 |

15. **Table S15:** Cartesian Coordinates for the low-energy geometry of the **EEEA** tetrapeptide calculated using the B3LYP method and the 6-311+G(*2df,2pd*) level of theory.

| **Standard Orientation** | | | | | |
| --- | --- | --- | --- | --- | --- |
| Center Number | Atomic Number | Atomic Type | Coordinates (Angstroms) | | |
|  |  |  | X | Y | Z |
| 1 | 6 | 0 | 4.591312 | -0.954277 | -0.079638 |
| 2 | 6 | 0 | 5.866910 | -0.269510 | 0.448095 |
| 3 | 6 | 0 | 7.096610 | -1.176489 | 0.623999 |
| 4 | 6 | 0 | 7.938639 | -1.612041 | -0.616876 |
| 5 | 8 | 0 | 9.001658 | -2.214702 | -0.339081 |
| 6 | 8 | 0 | 7.496327 | -1.330993 | -1.761886 |
| 7 | 1 | 0 | 7.802548 | -0.683159 | 1.298355 |
| 8 | 1 | 0 | 6.802877 | -2.095156 | 1.143045 |
| 9 | 1 | 0 | 6.109032 | 0.560745 | -0.220522 |
| 10 | 1 | 0 | 5.620297 | 0.159343 | 1.421600 |
| 11 | 7 | 0 | 4.638755 | -1.082390 | -1.539777 |
| 12 | 1 | 0 | 4.109900 | -1.890013 | -1.844254 |
| 13 | 1 | 0 | 5.617270 | -1.195868 | -1.828790 |
| 14 | 6 | 0 | 3.328111 | -0.204579 | 0.403439 |
| 15 | 8 | 0 | 2.975245 | -0.261752 | 1.577342 |
| 16 | 7 | 0 | 2.685436 | 0.490914 | -0.558461 |
| 17 | 6 | 0 | 1.367254 | 1.096598 | -0.425041 |
| 18 | 6 | 0 | 1.359828 | 2.298462 | 0.551696 |
| 19 | 6 | 0 | 0.236311 | 3.331017 | 0.375504 |
| 20 | 6 | 0 | 0.613073 | 4.814252 | 0.755330 |
| 21 | 8 | 0 | 1.809201 | 5.145664 | 0.605866 |
| 22 | 8 | 0 | -0.355475 | 5.518981 | 1.129089 |
| 23 | 1 | 0 | -0.075865 | 3.372891 | -0.674998 |
| 24 | 1 | 0 | -0.662241 | 3.100524 | 0.954752 |
| 25 | 1 | 0 | 2.290101 | 2.839852 | 0.379765 |
| 26 | 1 | 0 | 1.401170 | 1.931994 | 1.577627 |
| 27 | 6 | 0 | 0.303210 | -0.035788 | -0.249027 |
| 28 | 8 | 0 | 0.517751 | -1.158125 | -0.671126 |
| 29 | 7 | 0 | -0.862500 | 0.359003 | 0.337465 |
| 30 | 6 | 0 | -2.111423 | -0.382445 | 0.520576 |
| 31 | 1 | 0 | -2.572622 | 0.068357 | 1.403620 |
| 32 | 6 | 0 | -3.077468 | -0.187264 | -0.678137 |
| 33 | 8 | 0 | -2.800528 | -0.579356 | -1.798952 |
| 34 | 7 | 0 | -4.238082 | 0.442430 | -0.346089 |
| 35 | 6 | 0 | -5.437161 | 0.559071 | -1.179872 |
| 36 | 1 | 0 | -6.081166 | 1.256442 | -0.634233 |
| 37 | 6 | 0 | -6.298500 | -0.754662 | -1.255257 |
| 38 | 8 | 0 | -7.143385 | -0.801818 | -2.157847 |
| 39 | 8 | 0 | -6.110937 | -1.622089 | -0.347044 |
| 40 | 6 | 0 | -5.155323 | 1.177552 | -2.548522 |
| 41 | 1 | 0 | -4.647946 | 2.137488 | -2.432885 |
| 42 | 1 | 0 | -6.100521 | 1.324512 | -3.066371 |
| 43 | 1 | 0 | -4.523286 | 0.526508 | -3.145481 |
| 44 | 1 | 0 | -4.389118 | 0.578372 | 0.638528 |
| 45 | 6 | 0 | -1.922906 | -1.878970 | 0.813844 |
| 46 | 6 | 0 | -3.245867 | -2.633417 | 1.047665 |
| 47 | 1 | 0 | -2.995867 | -3.675390 | 1.255722 |
| 48 | 1 | 0 | -3.862476 | -2.611332 | 0.150277 |
| 49 | 6 | 0 | -4.040517 | -2.133799 | 2.250131 |
| 50 | 8 | 0 | -3.627107 | -2.260693 | 3.387189 |
| 51 | 8 | 0 | -5.201358 | -1.543029 | 2.003768 |
| 52 | 1 | 0 | -5.496261 | -1.552614 | 1.010937 |
| 53 | 1 | 0 | -1.281416 | -1.975134 | 1.689848 |
| 54 | 1 | 0 | -1.401647 | -2.332434 | -0.023355 |
| 55 | 1 | 0 | -0.905640 | 1.329401 | 0.603087 |
| 56 | 1 | 0 | 1.139249 | 1.502077 | -1.416765 |
| 57 | 1 | 0 | 3.062307 | 0.320594 | -1.481376 |
| 58 | 1 | 0 | 4.512389 | -1.926652 | 0.423918 |

16. **Table S16:** Cartesian Coordinates for the low-energy geometry of the **EGED** tetrapeptide calculated using the B3LYP method and the 6-311+G(*2df,2pd*) level of theory.

| **Standard Orientation** | | | | | |
| --- | --- | --- | --- | --- | --- |
| Center Number | Atomic Number | Atomic Type | Coordinates (Angstroms) | | |
|  |  |  | X | Y | Z |
| 1 | 6 | 0 | 5.636921 | 0.053937 | 0.514722 |
| 2 | 7 | 0 | 5.511749 | 0.767783 | -0.755230 |
| 3 | 1 | 0 | 6.406666 | 0.702624 | -1.253399 |
| 4 | 1 | 0 | 5.298026 | 1.745832 | -0.605783 |
| 5 | 1 | 0 | 6.036259 | 0.686252 | 1.320872 |
| 6 | 6 | 0 | 6.596007 | -1.146072 | 0.372263 |
| 7 | 6 | 0 | 8.091779 | -0.832138 | 0.529508 |
| 8 | 6 | 0 | 8.865714 | -0.117808 | -0.623238 |
| 9 | 8 | 0 | 10.095383 | 0.008029 | -0.422443 |
| 10 | 8 | 0 | 8.209849 | 0.253806 | -1.631248 |
| 11 | 1 | 0 | 8.249663 | -0.237592 | 1.435261 |
| 12 | 1 | 0 | 8.629281 | -1.768583 | 0.707168 |
| 13 | 1 | 0 | 6.413654 | -1.608438 | -0.601508 |
| 14 | 1 | 0 | 6.319386 | -1.871925 | 1.139311 |
| 15 | 6 | 0 | 4.277530 | -0.412644 | 1.070747 |
| 16 | 8 | 0 | 4.189903 | -0.874359 | 2.210400 |
| 17 | 7 | 0 | 3.216625 | -0.265355 | 0.256429 |
| 18 | 1 | 0 | 3.363635 | 0.116330 | -0.670255 |
| 19 | 6 | 0 | 1.875410 | -0.618961 | 0.653596 |
| 20 | 6 | 0 | 0.845424 | -0.076269 | -0.348011 |
| 21 | 8 | 0 | 1.204239 | 0.445959 | -1.401996 |
| 22 | 7 | 0 | -0.430166 | -0.238670 | 0.033185 |
| 23 | 6 | 0 | -1.606894 | 0.028637 | -0.794287 |
| 24 | 6 | 0 | -2.768196 | -0.853315 | -0.252974 |
| 25 | 7 | 0 | -3.781097 | -0.960344 | -1.046045 |
| 26 | 6 | 0 | -5.023242 | -1.669097 | -0.691024 |
| 27 | 6 | 0 | -4.907281 | -3.120760 | -0.183217 |
| 28 | 8 | 0 | -5.642475 | -3.997439 | -0.608029 |
| 29 | 8 | 0 | -4.011428 | -3.394681 | 0.757601 |
| 30 | 1 | 0 | -3.429650 | -2.566063 | 0.940568 |
| 31 | 1 | 0 | -5.579023 | -1.773561 | -1.622818 |
| 32 | 6 | 0 | -5.960913 | -0.858356 | 0.250565 |
| 33 | 1 | 0 | -6.749300 | -1.524853 | 0.605257 |
| 34 | 1 | 0 | -5.384362 | -0.521907 | 1.144601 |
| 35 | 6 | 0 | -6.655587 | 0.337786 | -0.396133 |
| 36 | 8 | 0 | -7.450871 | 0.964757 | 0.594754 |
| 37 | 1 | 0 | -6.853903 | 1.238323 | 1.338831 |
| 38 | 8 | 0 | -5.620292 | 1.332597 | -0.708593 |
| 39 | 1 | 0 | -4.926056 | 0.835682 | -1.165579 |
| 40 | 8 | 0 | -2.557330 | -1.369688 | 0.913239 |
| 41 | 1 | 0 | -1.389458 | -0.287065 | -1.813839 |
| 42 | 6 | 0 | -1.944281 | 1.530281 | -0.847235 |
| 43 | 1 | 0 | -1.037010 | 2.059176 | -1.144220 |
| 44 | 1 | 0 | -2.679115 | 1.681212 | -1.636181 |
| 45 | 6 | 0 | -2.494546 | 2.091947 | 0.471735 |
| 46 | 1 | 0 | -1.964761 | 1.697102 | 1.336700 |
| 47 | 1 | 0 | -3.551521 | 1.795493 | 0.571295 |
| 48 | 6 | 0 | -2.477288 | 3.581887 | 0.589630 |
| 49 | 8 | 0 | -2.312099 | 4.221663 | 1.607510 |
| 50 | 8 | 0 | -2.716651 | 4.224437 | -0.587380 |
| 51 | 1 | 0 | -2.755186 | 5.164118 | -0.364245 |
| 52 | 1 | 0 | -0.675114 | -0.782016 | 0.856935 |
| 53 | 1 | 0 | 1.758629 | -1.706163 | 0.718675 |
| 54 | 1 | 0 | 1.670315 | -0.232859 | 1.653586 |

17. **Table S17:** Cartesian Coordinates for the low-energy geometry of the **EDEA** tetrapeptide calculated using the B3LYP method and the 6-311+G(*2df,2pd*) level of theory.

| **Standard Orientation** | | | | | |
| --- | --- | --- | --- | --- | --- |
| Center Number | Atomic Number | Atomic Type | Coordinates (Angstroms) | | |
|  |  |  | X | Y | Z |
| 1 | 6 | 0 | 5.025402 | 0.662809 | 0.205199 |
| 2 | 7 | 0 | 5.128694 | 0.175579 | 1.587971 |
| 3 | 1 | 0 | 4.621063 | 0.792091 | 2.213008 |
| 4 | 1 | 0 | 6.121665 | 0.250360 | 1.816256 |
| 5 | 1 | 0 | 5.218939 | 1.740910 | 0.131980 |
| 6 | 6 | 0 | 6.074437 | -0.044865 | -0.667338 |
| 7 | 6 | 0 | 7.574236 | 0.211447 | -0.300771 |
| 8 | 8 | 0 | 7.825633 | 0.667032 | 0.849057 |
| 9 | 8 | 0 | 8.397600 | -0.091734 | -1.192918 |
| 10 | 1 | 0 | 5.911667 | -1.126058 | -0.608569 |
| 11 | 1 | 0 | 5.927335 | 0.240641 | -1.707608 |
| 12 | 6 | 0 | 3.608337 | 0.495343 | -0.377441 |
| 13 | 8 | 0 | 3.273684 | 1.103686 | -1.391572 |
| 14 | 7 | 0 | 2.792522 | -0.328364 | 0.315521 |
| 15 | 1 | 0 | 3.188439 | -0.798258 | 1.118566 |
| 16 | 6 | 0 | 1.391709 | -0.612451 | -0.003568 |
| 17 | 1 | 0 | 0.996322 | 0.231249 | -0.561893 |
| 18 | 6 | 0 | 0.694898 | -0.779673 | 1.357872 |
| 19 | 8 | 0 | 1.246346 | -1.441183 | 2.239275 |
| 20 | 7 | 0 | -0.491203 | -0.174105 | 1.615825 |
| 21 | 6 | 0 | -1.462005 | 0.497227 | 0.751926 |
| 22 | 1 | 0 | -1.126718 | 0.406912 | -0.276268 |
| 23 | 6 | 0 | -2.811415 | -0.230581 | 0.918855 |
| 24 | 8 | 0 | -3.367433 | -0.236971 | 2.021049 |
| 25 | 7 | 0 | -3.291806 | -0.767449 | -0.208179 |
| 26 | 6 | 0 | -4.614113 | -1.366270 | -0.348379 |
| 27 | 1 | 0 | -4.657653 | -1.690982 | -1.391527 |
| 28 | 6 | 0 | -5.794871 | -0.335814 | -0.224722 |
| 29 | 8 | 0 | -6.919071 | -0.819822 | -0.013914 |
| 30 | 8 | 0 | -5.531135 | 0.888708 | -0.416734 |
| 31 | 6 | 0 | -4.804966 | -2.604844 | 0.532927 |
| 32 | 1 | 0 | -4.025605 | -3.338391 | 0.317177 |
| 33 | 1 | 0 | -5.783368 | -3.039682 | 0.337088 |
| 34 | 1 | 0 | -4.747998 | -2.332584 | 1.585019 |
| 35 | 1 | 0 | -2.607635 | -0.907372 | -0.959235 |
| 36 | 6 | 0 | -1.622983 | 1.973443 | 1.178065 |
| 37 | 6 | 0 | -2.897762 | 2.690154 | 0.690624 |
| 38 | 1 | 0 | -2.941933 | 3.655063 | 1.200974 |
| 39 | 1 | 0 | -3.778447 | 2.119165 | 0.978461 |
| 40 | 6 | 0 | -2.935938 | 3.021149 | -0.799026 |
| 41 | 8 | 0 | -2.213809 | 3.883671 | -1.268045 |
| 42 | 8 | 0 | -3.803842 | 2.366118 | -1.544236 |
| 43 | 1 | 0 | -4.406206 | 1.700028 | -1.041791 |
| 44 | 1 | 0 | -0.738093 | 2.526165 | 0.857523 |
| 45 | 1 | 0 | -1.644245 | 1.993621 | 2.269844 |
| 46 | 1 | 0 | -0.855408 | -0.411232 | 2.529797 |
| 47 | 6 | 0 | 1.261063 | -1.888919 | -0.863027 |
| 48 | 6 | 0 | -0.126965 | -2.216347 | -1.486015 |
| 49 | 8 | 0 | -0.318599 | -3.411997 | -1.781106 |
| 50 | 8 | 0 | -0.907902 | -1.240267 | -1.674798 |
| 51 | 1 | 0 | 1.954352 | -1.770035 | -1.699188 |
| 52 | 1 | 0 | 1.599917 | -2.750011 | -0.287120 |

18. **Table S18:** Cartesian Coordinates for the low-energy geometry of the **EAQG** tetrapeptide calculated using the B3LYP method and the 6-311+G(*2df,2pd*) level of theory.

| **Standard Orientation** | | | | | |
| --- | --- | --- | --- | --- | --- |
| Center Number | Atomic Number | Atomic Type | Coordinates (Angstroms) | | |
|  |  |  | X | Y | Z |
| 1 | 6 | 0 | 3.385515 | 1.963532 | 0.676394 |
| 2 | 7 | 0 | 3.276916 | 2.207009 | 2.117392 |
| 3 | 1 | 0 | 3.747331 | 3.066105 | 2.372396 |
| 4 | 1 | 0 | 3.726346 | 1.441442 | 2.607753 |
| 5 | 1 | 0 | 4.417720 | 2.114989 | 0.318164 |
| 6 | 6 | 0 | 3.202366 | 0.447037 | 0.457753 |
| 7 | 8 | 0 | 3.835397 | -0.325267 | 1.171608 |
| 8 | 7 | 0 | 2.412290 | 0.048737 | -0.557543 |
| 9 | 6 | 0 | 2.373966 | -1.325342 | -1.052365 |
| 10 | 1 | 0 | 1.577701 | -1.326218 | -1.798460 |
| 11 | 6 | 0 | 1.990890 | -2.383570 | 0.007175 |
| 12 | 8 | 0 | 2.726495 | -3.329719 | 0.238590 |
| 13 | 7 | 0 | 0.798822 | -2.269781 | 0.658483 |
| 14 | 6 | 0 | -0.407235 | -1.497392 | 0.407808 |
| 15 | 1 | 0 | -0.144170 | -0.499678 | 0.060807 |
| 16 | 6 | 0 | -1.364170 | -2.202539 | -0.617912 |
| 17 | 6 | 0 | -1.395459 | -1.616943 | -2.043415 |
| 18 | 6 | 0 | -2.376765 | -0.462557 | -2.184251 |
| 19 | 7 | 0 | -1.898944 | 0.698996 | -2.624342 |
| 20 | 1 | 0 | -0.902449 | 0.960800 | -2.494891 |
| 21 | 1 | 0 | -2.521386 | 1.493542 | -2.576055 |
| 22 | 8 | 0 | -3.572726 | -0.641780 | -1.887808 |
| 23 | 1 | 0 | -0.404978 | -1.320007 | -2.381996 |
| 24 | 1 | 0 | -1.753283 | -2.392421 | -2.724199 |
| 25 | 1 | 0 | -2.388849 | -2.212567 | -0.244929 |
| 26 | 1 | 0 | -1.047759 | -3.243549 | -0.663964 |
| 27 | 6 | 0 | -1.125013 | -1.409111 | 1.776932 |
| 28 | 8 | 0 | -0.905182 | -2.249849 | 2.639347 |
| 29 | 7 | 0 | -2.018250 | -0.414195 | 1.990486 |
| 30 | 1 | 0 | -2.470615 | -0.451286 | 2.893619 |
| 31 | 6 | 0 | -2.532036 | 0.596371 | 1.080400 |
| 32 | 1 | 0 | -2.149900 | 1.589853 | 1.327236 |
| 33 | 1 | 0 | -2.223500 | 0.389822 | 0.067013 |
| 34 | 6 | 0 | -4.058274 | 0.662402 | 1.224800 |
| 35 | 8 | 0 | -4.552407 | 0.830062 | 2.314711 |
| 36 | 8 | 0 | -4.799302 | 0.555235 | 0.130325 |
| 37 | 1 | 0 | -4.305782 | 0.212074 | -0.662993 |
| 38 | 1 | 0 | 0.678239 | -2.948431 | 1.400574 |
| 39 | 6 | 0 | 3.677431 | -1.716413 | -1.749077 |
| 40 | 1 | 0 | 3.890393 | -0.999399 | -2.542550 |
| 41 | 1 | 0 | 3.598896 | -2.713290 | -2.179712 |
| 42 | 1 | 0 | 4.497426 | -1.723492 | -1.035670 |
| 43 | 1 | 0 | 1.813752 | 0.727460 | -1.053839 |
| 44 | 6 | 0 | 2.514878 | 2.945018 | -0.120634 |
| 45 | 6 | 0 | 1.008644 | 2.968238 | 0.197946 |
| 46 | 1 | 0 | 0.687564 | 3.959218 | 0.513043 |
| 47 | 1 | 0 | 0.806759 | 2.314076 | 1.050979 |
| 48 | 6 | 0 | 0.070948 | 2.529874 | -0.939721 |
| 49 | 8 | 0 | 0.520060 | 1.638291 | -1.747809 |
| 50 | 8 | 0 | -1.066645 | 3.019929 | -0.978589 |
| 51 | 1 | 0 | 2.664053 | 2.759675 | -1.185299 |
| 52 | 1 | 0 | 2.931015 | 3.939090 | 0.067627 |

19. **Table S19:** Cartesian Coordinates for the low-energy geometry of the **QGPK** tetrapeptide calculated using the B3LYP method and the 6-311+G(*2df,2pd*) level of theory.

| **Standard Orientation** | | | | | |
| --- | --- | --- | --- | --- | --- |
| Center Number | Atomic Number | Atomic Type | Coordinates (Angstroms) | | |
|  |  |  | X | Y | Z |
| 1 | 6 | 0 | 3.792555 | 1.461862 | 0.046739 |
| 2 | 7 | 0 | 5.066165 | 2.086983 | 0.372405 |
| 3 | 1 | 0 | 5.573394 | 2.336366 | -0.467393 |
| 4 | 1 | 0 | 4.894562 | 2.953870 | 0.871992 |
| 5 | 1 | 0 | 3.972051 | 0.742369 | -0.752177 |
| 6 | 6 | 0 | 3.253469 | 0.710896 | 1.283304 |
| 7 | 6 | 0 | 4.094146 | -0.506230 | 1.649722 |
| 8 | 6 | 0 | 3.939624 | -1.663249 | 0.674073 |
| 9 | 7 | 0 | 4.847327 | -2.656585 | 0.802366 |
| 10 | 1 | 0 | 5.602634 | -2.605326 | 1.462361 |
| 11 | 1 | 0 | 4.788546 | -3.457632 | 0.196494 |
| 12 | 8 | 0 | 3.045107 | -1.715567 | -0.166068 |
| 13 | 1 | 0 | 5.145549 | -0.221491 | 1.717547 |
| 14 | 1 | 0 | 3.804311 | -0.878238 | 2.636267 |
| 15 | 1 | 0 | 2.219662 | 0.401761 | 1.127641 |
| 16 | 1 | 0 | 3.251794 | 1.411583 | 2.120021 |
| 17 | 6 | 0 | 2.736273 | 2.482532 | -0.411717 |
| 18 | 8 | 0 | 2.761222 | 3.633673 | -0.012965 |
| 19 | 7 | 0 | 1.753056 | 2.066614 | -1.267668 |
| 20 | 1 | 0 | 1.067836 | 2.774593 | -1.486406 |
| 21 | 6 | 0 | 1.489889 | 0.739405 | -1.761182 |
| 22 | 1 | 0 | 1.832965 | 0.610369 | -2.797430 |
| 23 | 1 | 0 | 2.009809 | -0.011460 | -1.168151 |
| 24 | 6 | 0 | -0.002340 | 0.445961 | -1.752141 |
| 25 | 8 | 0 | -0.823854 | 1.362996 | -1.577659 |
| 26 | 7 | 0 | -0.391273 | -0.820061 | -1.965242 |
| 27 | 6 | 0 | -1.816929 | -1.141370 | -2.123266 |
| 28 | 6 | 0 | -1.811496 | -2.601157 | -2.634414 |
| 29 | 6 | 0 | -0.455220 | -3.161990 | -2.186825 |
| 30 | 6 | 0 | 0.489610 | -1.966866 | -2.294893 |
| 31 | 1 | 0 | 0.867661 | -1.855132 | -3.314292 |
| 32 | 1 | 0 | 1.332954 | -2.022285 | -1.612014 |
| 33 | 1 | 0 | -0.499391 | -3.513488 | -1.155269 |
| 34 | 1 | 0 | -0.126963 | -3.997488 | -2.800800 |
| 35 | 1 | 0 | -1.878897 | -2.592385 | -3.721921 |
| 36 | 1 | 0 | -2.660168 | -3.169565 | -2.260667 |
| 37 | 1 | 0 | -2.271172 | -0.472451 | -2.852073 |
| 38 | 6 | 0 | -2.654004 | -0.953578 | -0.848211 |
| 39 | 8 | 0 | -3.805145 | -0.523356 | -0.947341 |
| 40 | 7 | 0 | -2.075152 | -1.295938 | 0.313096 |
| 41 | 1 | 0 | -1.095307 | -1.547347 | 0.317524 |
| 42 | 6 | 0 | -2.688558 | -1.256613 | 1.637310 |
| 43 | 6 | 0 | -3.487980 | 0.027755 | 1.947461 |
| 44 | 1 | 0 | -4.381376 | -0.013796 | 1.331223 |
| 45 | 1 | 0 | -3.815149 | -0.034431 | 2.984470 |
| 46 | 6 | 0 | -2.739535 | 1.344958 | 1.688485 |
| 47 | 1 | 0 | -2.146769 | 1.619876 | 2.560945 |
| 48 | 1 | 0 | -2.014953 | 1.198733 | 0.886509 |
| 49 | 6 | 0 | -3.697296 | 2.504508 | 1.323461 |
| 50 | 1 | 0 | -4.724660 | 2.136980 | 1.266549 |
| 51 | 6 | 0 | -3.372698 | 3.225473 | 0.021643 |
| 52 | 1 | 0 | -2.397516 | 3.707078 | 0.055121 |
| 53 | 1 | 0 | -4.122309 | 3.985003 | -0.188770 |
| 54 | 7 | 0 | -3.339126 | 2.283907 | -1.165669 |
| 55 | 1 | 0 | -3.640290 | 2.756706 | -2.014542 |
| 56 | 1 | 0 | -2.373914 | 1.912682 | -1.340108 |
| 57 | 1 | 0 | -3.920534 | 1.441177 | -1.042473 |
| 58 | 1 | 0 | -3.701628 | 3.265682 | 2.103969 |
| 59 | 1 | 0 | -3.385270 | -2.093758 | 1.741770 |
| 60 | 6 | 0 | -1.562923 | -1.509738 | 2.634924 |
| 61 | 8 | 0 | -0.404306 | -1.641145 | 2.331023 |
| 62 | 8 | 0 | -2.022517 | -1.585189 | 3.888201 |
| 63 | 1 | 0 | -1.278016 | -1.769641 | 4.480706 |

20. **Table S20:** Cartesian Coordinates for the low-energy geometry of the **AEEE** tetrapeptide calculated using the B3LYP method and the 6-311+G(*2df,2pd*) level of theory.

| **Standard Orientation** | | | | | |
| --- | --- | --- | --- | --- | --- |
| Center Number | Atomic Number | Atomic Type | Coordinates (Angstroms) | | |
|  |  |  | X | Y | Z |
| 1 | 6 | 0 | 4.681788 | -1.010717 | 1.458990 |
| 2 | 7 | 0 | 5.155825 | -2.390477 | 1.385528 |
| 3 | 1 | 0 | 5.568234 | -2.525203 | 0.467145 |
| 4 | 1 | 0 | 4.356312 | -3.026562 | 1.438666 |
| 5 | 1 | 0 | 3.891216 | -0.978922 | 2.208959 |
| 6 | 6 | 0 | 4.104071 | -0.513521 | 0.118664 |
| 7 | 8 | 0 | 4.638555 | -0.813094 | -0.945696 |
| 8 | 7 | 0 | 3.026731 | 0.301022 | 0.213574 |
| 9 | 1 | 0 | 2.623972 | 0.425151 | 1.128011 |
| 10 | 6 | 0 | 2.368638 | 0.942924 | -0.914403 |
| 11 | 6 | 0 | 2.162480 | 2.452381 | -0.659317 |
| 12 | 1 | 0 | 1.689784 | 2.609861 | 0.313293 |
| 13 | 6 | 0 | 3.466935 | 3.250714 | -0.720770 |
| 14 | 1 | 0 | 4.216172 | 2.772382 | -0.080531 |
| 15 | 1 | 0 | 3.871888 | 3.230149 | -1.734826 |
| 16 | 6 | 0 | 3.386751 | 4.739567 | -0.251210 |
| 17 | 8 | 0 | 4.182079 | 5.525581 | -0.826803 |
| 18 | 8 | 0 | 2.578566 | 4.993560 | 0.671681 |
| 19 | 1 | 0 | 1.464064 | 2.824456 | -1.409810 |
| 20 | 6 | 0 | 1.080198 | 0.234984 | -1.388069 |
| 21 | 8 | 0 | 0.617094 | 0.516903 | -2.487588 |
| 22 | 7 | 0 | 0.513253 | -0.703807 | -0.580316 |
| 23 | 6 | 0 | -0.419210 | -1.669007 | -1.152741 |
| 24 | 6 | 0 | -0.514620 | -2.933063 | -0.272580 |
| 25 | 6 | 0 | 0.772367 | -3.756980 | -0.132942 |
| 26 | 6 | 0 | 1.756809 | -3.347833 | 0.994572 |
| 27 | 8 | 0 | 2.637271 | -4.190230 | 1.279565 |
| 28 | 8 | 0 | 1.606178 | -2.206561 | 1.510325 |
| 29 | 1 | 0 | 0.518431 | -4.804848 | 0.041923 |
| 30 | 1 | 0 | 1.339407 | -3.744103 | -1.069333 |
| 31 | 1 | 0 | -0.869514 | -2.648285 | 0.719610 |
| 32 | 1 | 0 | -1.291162 | -3.553583 | -0.723475 |
| 33 | 6 | 0 | -1.853626 | -1.182583 | -1.411463 |
| 34 | 7 | 0 | -2.411791 | -0.324103 | -0.527538 |
| 35 | 6 | 0 | -3.806089 | 0.085990 | -0.640075 |
| 36 | 6 | 0 | -4.243482 | 0.918924 | 0.588979 |
| 37 | 6 | 0 | -5.607981 | 1.596316 | 0.440870 |
| 38 | 6 | 0 | -6.300801 | 2.031545 | 1.778691 |
| 39 | 8 | 0 | -7.177387 | 2.920136 | 1.647999 |
| 40 | 8 | 0 | -5.952705 | 1.426463 | 2.818515 |
| 41 | 1 | 0 | -5.540430 | 2.469877 | -0.210107 |
| 42 | 1 | 0 | -6.314460 | 0.915857 | -0.042257 |
| 43 | 1 | 0 | -3.473550 | 1.677826 | 0.754879 |
| 44 | 1 | 0 | -4.260863 | 0.287820 | 1.477970 |
| 45 | 6 | 0 | -4.731700 | -1.115214 | -0.793249 |
| 46 | 8 | 0 | -4.431696 | -2.126298 | 0.056559 |
| 47 | 1 | 0 | -5.101055 | -2.805386 | -0.097407 |
| 48 | 8 | 0 | -5.696509 | -1.161568 | -1.516143 |
| 49 | 1 | 0 | -3.943873 | 0.667728 | -1.552972 |
| 50 | 1 | 0 | -1.840893 | 0.067293 | 0.203934 |
| 51 | 8 | 0 | -2.500753 | -1.663074 | -2.336717 |
| 52 | 1 | 0 | -0.065109 | -1.951017 | -2.146750 |
| 53 | 1 | 0 | 1.015358 | -1.054424 | 0.247486 |
| 54 | 1 | 0 | 3.045335 | 0.821807 | -1.759824 |
| 55 | 6 | 0 | 5.824466 | -0.072165 | 1.871972 |
| 56 | 1 | 0 | 6.627073 | -0.114479 | 1.132775 |
| 57 | 1 | 0 | 6.226609 | -0.394203 | 2.833554 |
| 58 | 1 | 0 | 5.491410 | 0.964233 | 1.951277 |

21. **Table S21:** Cartesian Coordinates for the low-energy geometry of the **EEVA** tetrapeptide calculated using the B3LYP method and the 6-311+G(*2df,2pd*) level of theory.

| **Standard Orientation** | | | | | |
| --- | --- | --- | --- | --- | --- |
| Center Number | Atomic Number | Atomic Type | Coordinates (Angstroms) | | |
|  |  |  | X | Y | Z |
| 1 | 6 | 0 | -4.040357 | -0.410818 | -0.168444 |
| 2 | 6 | 0 | -4.976629 | 0.474058 | -1.014538 |
| 3 | 6 | 0 | -6.320354 | -0.163143 | -1.398599 |
| 4 | 6 | 0 | -7.469025 | -0.222369 | -0.341871 |
| 5 | 8 | 0 | -8.560980 | -0.637410 | -0.781132 |
| 6 | 8 | 0 | -7.201682 | 0.149054 | 0.833978 |
| 7 | 1 | 0 | -6.734125 | 0.375635 | -2.255088 |
| 8 | 1 | 0 | -6.159637 | -1.185987 | -1.754814 |
| 9 | 1 | 0 | -5.155127 | 1.402739 | -0.465856 |
| 10 | 1 | 0 | -4.436077 | 0.731633 | -1.927367 |
| 11 | 7 | 0 | -4.481520 | -0.464283 | 1.228262 |
| 12 | 1 | 0 | -4.346875 | -1.388823 | 1.616035 |
| 13 | 1 | 0 | -5.486664 | -0.235719 | 1.271300 |
| 14 | 6 | 0 | -2.590218 | 0.100953 | -0.297573 |
| 15 | 7 | 0 | -2.097674 | 0.650694 | 0.826908 |
| 16 | 6 | 0 | -0.805407 | 1.324516 | 1.019889 |
| 17 | 6 | 0 | -0.623208 | 2.508909 | 0.049903 |
| 18 | 6 | 0 | 0.490942 | 3.488351 | 0.426958 |
| 19 | 6 | 0 | 1.914016 | 3.172731 | -0.103750 |
| 20 | 8 | 0 | 2.007235 | 2.270547 | -0.984070 |
| 21 | 8 | 0 | 2.839465 | 3.862083 | 0.371840 |
| 22 | 1 | 0 | 0.242170 | 4.475678 | 0.026217 |
| 23 | 1 | 0 | 0.562503 | 3.620898 | 1.509397 |
| 24 | 1 | 0 | -1.584053 | 3.031514 | 0.049242 |
| 25 | 1 | 0 | -0.452402 | 2.147512 | -0.961285 |
| 26 | 6 | 0 | 0.351336 | 0.295430 | 1.084758 |
| 27 | 8 | 0 | 0.974054 | 0.118337 | 2.125319 |
| 28 | 7 | 0 | 0.576829 | -0.401742 | -0.047819 |
| 29 | 6 | 0 | 1.497321 | -1.522561 | -0.156516 |
| 30 | 1 | 0 | 1.519698 | -1.752535 | -1.226516 |
| 31 | 6 | 0 | 2.965918 | -1.165689 | 0.191630 |
| 32 | 8 | 0 | 3.693155 | -1.980562 | 0.758606 |
| 33 | 7 | 0 | 3.440911 | 0.009044 | -0.262511 |
| 34 | 1 | 0 | 2.829305 | 0.776519 | -0.593206 |
| 35 | 6 | 0 | 4.824137 | 0.341955 | 0.019847 |
| 36 | 6 | 0 | 5.199350 | 1.706895 | -0.572674 |
| 37 | 1 | 0 | 5.087635 | 1.698018 | -1.657299 |
| 38 | 1 | 0 | 6.236549 | 1.933561 | -0.322213 |
| 39 | 1 | 0 | 4.545085 | 2.485975 | -0.178627 |
| 40 | 6 | 0 | 5.808824 | -0.693452 | -0.503835 |
| 41 | 8 | 0 | 5.438911 | -1.265475 | -1.675790 |
| 42 | 1 | 0 | 6.152196 | -1.877272 | -1.901335 |
| 43 | 8 | 0 | 6.883714 | -0.929601 | -0.003776 |
| 44 | 1 | 0 | 4.993478 | 0.358363 | 1.098586 |
| 45 | 6 | 0 | 1.046270 | -2.816936 | 0.581238 |
| 46 | 6 | 0 | -0.476638 | -2.958997 | 0.640250 |
| 47 | 1 | 0 | -0.739354 | -3.910876 | 1.108322 |
| 48 | 1 | 0 | -0.923004 | -2.944577 | -0.357051 |
| 49 | 1 | 0 | -0.940130 | -2.164351 | 1.220597 |
| 50 | 6 | 0 | 1.657364 | -4.054082 | -0.090151 |
| 51 | 1 | 0 | 1.270562 | -4.169857 | -1.108117 |
| 52 | 1 | 0 | 1.394145 | -4.958724 | 0.464082 |
| 53 | 1 | 0 | 2.741417 | -3.983055 | -0.132473 |
| 54 | 1 | 0 | 1.430613 | -2.757253 | 1.599429 |
| 55 | 1 | 0 | -0.058672 | -0.226520 | -0.818690 |
| 56 | 1 | 0 | -0.840623 | 1.722559 | 2.033114 |
| 57 | 1 | 0 | -2.761214 | 0.605678 | 1.594474 |
| 58 | 8 | 0 | -1.969163 | -0.011819 | -1.358335 |
| 59 | 1 | 0 | -4.016747 | -1.402480 | -0.636995 |

22. **Table S22:** Cartesian Coordinates for the low-energy geometry of the **EEEI** tetrapeptide calculated using the B3LYP method and the 6-311+G(*2df,2pd*) level of theory.

| **Standard Orientation** | | | | | |
| --- | --- | --- | --- | --- | --- |
| Center Number | Atomic Number | Atomic Type | Coordinates (Angstroms) | | |
|  |  |  | X | Y | Z |
| 1 | 6 | 0 | -3.983945 | -0.370521 | -0.261349 |
| 2 | 6 | 0 | -4.801243 | 0.326687 | -1.366053 |
| 3 | 6 | 0 | -5.994167 | -0.469494 | -1.923223 |
| 4 | 6 | 0 | -7.347409 | -0.484682 | -1.144365 |
| 5 | 8 | 0 | -8.318835 | -0.938845 | -1.789309 |
| 6 | 8 | 0 | -7.349692 | -0.055594 | 0.041062 |
| 7 | 1 | 0 | -6.231226 | -0.096385 | -2.922616 |
| 8 | 1 | 0 | -5.703443 | -1.515430 | -2.070246 |
| 9 | 1 | 0 | -5.149087 | 1.287927 | -0.978201 |
| 10 | 1 | 0 | -4.112417 | 0.538567 | -2.185681 |
| 11 | 7 | 0 | -4.693670 | -0.323159 | 1.020484 |
| 12 | 1 | 0 | -4.540945 | -1.170679 | 1.551355 |
| 13 | 1 | 0 | -5.702965 | -0.220608 | 0.844895 |
| 14 | 6 | 0 | -2.570949 | 0.263907 | -0.181253 |
| 15 | 7 | 0 | -2.377473 | 1.009807 | 0.925266 |
| 16 | 6 | 0 | -1.279328 | 1.897393 | 1.334967 |
| 17 | 6 | 0 | -0.510050 | 2.599403 | 0.198216 |
| 18 | 6 | 0 | 0.430734 | 3.695828 | 0.708954 |
| 19 | 6 | 0 | 0.866432 | 4.752598 | -0.363245 |
| 20 | 8 | 0 | 2.016550 | 5.233128 | -0.206368 |
| 21 | 8 | 0 | 0.014689 | 5.049736 | -1.233369 |
| 22 | 1 | 0 | -0.071610 | 4.266613 | 1.498730 |
| 23 | 1 | 0 | 1.328143 | 3.274592 | 1.163182 |
| 24 | 1 | 0 | -1.242129 | 3.062939 | -0.463893 |
| 25 | 1 | 0 | 0.038102 | 1.883242 | -0.410213 |
| 26 | 6 | 0 | -0.386346 | 1.271857 | 2.435769 |
| 27 | 8 | 0 | -0.375034 | 1.794857 | 3.549630 |
| 28 | 7 | 0 | 0.379953 | 0.174724 | 2.205051 |
| 29 | 6 | 0 | 0.587240 | -0.655187 | 1.028929 |
| 30 | 1 | 0 | -0.033136 | -0.293742 | 0.212432 |
| 31 | 6 | 0 | 0.222573 | -2.123780 | 1.358696 |
| 32 | 1 | 0 | 0.926418 | -2.464807 | 2.121194 |
| 33 | 1 | 0 | -0.768052 | -2.136958 | 1.812113 |
| 34 | 6 | 0 | 0.258363 | -3.073367 | 0.137419 |
| 35 | 1 | 0 | -0.731632 | -3.120028 | -0.314199 |
| 36 | 1 | 0 | 0.960546 | -2.709271 | -0.607513 |
| 37 | 6 | 0 | 0.660755 | -4.485761 | 0.536068 |
| 38 | 8 | 0 | 1.906046 | -4.852753 | 0.262538 |
| 39 | 1 | 0 | 2.450006 | -4.153694 | -0.237553 |
| 40 | 8 | 0 | -0.106478 | -5.247320 | 1.096746 |
| 41 | 6 | 0 | 2.084457 | -0.615160 | 0.662934 |
| 42 | 8 | 0 | 2.934533 | -0.721815 | 1.539680 |
| 43 | 7 | 0 | 2.363153 | -0.508278 | -0.649819 |
| 44 | 1 | 0 | 1.577324 | -0.450992 | -1.274316 |
| 45 | 6 | 0 | 3.682496 | -0.754532 | -1.241951 |
| 46 | 6 | 0 | 4.740490 | 0.324714 | -0.902071 |
| 47 | 6 | 0 | 4.127348 | 1.734776 | -0.848182 |
| 48 | 6 | 0 | 5.082178 | 2.853427 | -0.427685 |
| 49 | 1 | 0 | 4.525495 | 3.785638 | -0.319836 |
| 50 | 1 | 0 | 5.883087 | 3.014380 | -1.154005 |
| 51 | 1 | 0 | 5.548960 | 2.622603 | 0.534703 |
| 52 | 1 | 0 | 3.292445 | 1.737808 | -0.151299 |
| 53 | 1 | 0 | 3.697004 | 1.983787 | -1.824897 |
| 54 | 6 | 0 | 5.885558 | 0.252575 | -1.922152 |
| 55 | 1 | 0 | 6.286304 | -0.757374 | -1.969284 |
| 56 | 1 | 0 | 5.530489 | 0.542759 | -2.918035 |
| 57 | 1 | 0 | 6.695233 | 0.933801 | -1.653063 |
| 58 | 1 | 0 | 5.144157 | 0.081924 | 0.082685 |
| 59 | 1 | 0 | 3.504334 | -0.705462 | -2.320924 |
| 60 | 6 | 0 | 4.123962 | -2.253296 | -0.981782 |
| 61 | 8 | 0 | 3.172994 | -3.085255 | -1.110024 |
| 62 | 8 | 0 | 5.312743 | -2.495025 | -0.751846 |
| 63 | 1 | 0 | 1.001144 | -0.037203 | 2.974016 |
| 64 | 1 | 0 | -1.771419 | 2.692702 | 1.894621 |
| 65 | 1 | 0 | -3.194715 | 0.981717 | 1.528150 |
| 66 | 8 | 0 | -1.747814 | 0.061836 | -1.072080 |
| 67 | 1 | 0 | -3.792678 | -1.397915 | -0.594297 |

23. **Table S23:** Cartesian Coordinates for the low-energy geometry of the **EIDG** tetrapeptide calculated using the B3LYP method and the 6-311+G(*2df,2pd*) level of theory.

| **Standard Orientation** | | | | | |
| --- | --- | --- | --- | --- | --- |
| Center Number | Atomic Number | Atomic Type | Coordinates (Angstroms) | | |
|  |  |  | X | Y | Z |
| 1 | 6 | 0 | 3.641026 | 0.449818 | 0.467123 |
| 2 | 7 | 0 | 3.242519 | 1.302024 | -0.655595 |
| 3 | 1 | 0 | 3.980591 | 1.284679 | -1.374059 |
| 4 | 1 | 0 | 3.101784 | 2.258863 | -0.360153 |
| 5 | 1 | 0 | 4.040476 | 1.031427 | 1.307239 |
| 6 | 6 | 0 | 4.727515 | -0.560328 | 0.050536 |
| 7 | 1 | 0 | 4.755294 | -1.328890 | 0.824500 |
| 8 | 1 | 0 | 4.409190 | -1.039300 | -0.880002 |
| 9 | 6 | 0 | 6.140283 | 0.014453 | -0.133144 |
| 10 | 1 | 0 | 6.354624 | 0.739374 | 0.659640 |
| 11 | 1 | 0 | 6.872200 | -0.785497 | 0.003721 |
| 12 | 6 | 0 | 6.503095 | 0.700383 | -1.488142 |
| 13 | 8 | 0 | 5.555397 | 0.963723 | -2.276580 |
| 14 | 8 | 0 | 7.720935 | 0.924084 | -1.653602 |
| 15 | 6 | 0 | 2.444304 | -0.318215 | 1.080780 |
| 16 | 8 | 0 | 2.579399 | -0.983058 | 2.100887 |
| 17 | 7 | 0 | 1.300652 | -0.195910 | 0.362465 |
| 18 | 1 | 0 | 1.439928 | 0.386988 | -0.459315 |
| 19 | 6 | 0 | -0.003369 | -0.816353 | 0.594853 |
| 20 | 6 | 0 | 0.069009 | -2.295133 | 1.041805 |
| 21 | 6 | 0 | 0.899133 | -3.133544 | 0.056177 |
| 22 | 6 | 0 | 1.286993 | -4.515086 | 0.587301 |
| 23 | 1 | 0 | 1.910011 | -5.048313 | -0.134165 |
| 24 | 1 | 0 | 0.416896 | -5.142076 | 0.793777 |
| 25 | 1 | 0 | 1.858705 | -4.420166 | 1.512292 |
| 26 | 1 | 0 | 1.812421 | -2.595254 | -0.190821 |
| 27 | 1 | 0 | 0.344031 | -3.242916 | -0.883400 |
| 28 | 6 | 0 | -1.342238 | -2.872900 | 1.205103 |
| 29 | 1 | 0 | -1.917199 | -2.312616 | 1.940723 |
| 30 | 1 | 0 | -1.884101 | -2.861327 | 0.254168 |
| 31 | 1 | 0 | -1.301548 | -3.908718 | 1.544983 |
| 32 | 1 | 0 | 0.564689 | -2.303936 | 2.010858 |
| 33 | 1 | 0 | -0.500828 | -0.793335 | -0.376072 |
| 34 | 6 | 0 | -0.821724 | 0.037279 | 1.607253 |
| 35 | 8 | 0 | -0.719686 | -0.170005 | 2.806790 |
| 36 | 7 | 0 | -1.665734 | 0.997601 | 1.125222 |
| 37 | 6 | 0 | -1.770245 | 1.492317 | -0.242637 |
| 38 | 1 | 0 | -0.792424 | 1.721602 | -0.671336 |
| 39 | 6 | 0 | -2.441103 | 0.510372 | -1.206176 |
| 40 | 8 | 0 | -2.228143 | 0.515642 | -2.409936 |
| 41 | 7 | 0 | -3.347499 | -0.335593 | -0.629359 |
| 42 | 1 | 0 | -3.448761 | -0.290186 | 0.372591 |
| 43 | 6 | 0 | -4.089247 | -1.279993 | -1.401897 |
| 44 | 1 | 0 | -3.860099 | -1.098123 | -2.453684 |
| 45 | 1 | 0 | -3.796865 | -2.316604 | -1.194949 |
| 46 | 6 | 0 | -5.585177 | -1.198598 | -1.205034 |
| 47 | 8 | 0 | -6.222302 | -2.095515 | -2.009738 |
| 48 | 1 | 0 | -7.166759 | -2.003357 | -1.823456 |
| 49 | 8 | 0 | -6.183558 | -0.487497 | -0.445280 |
| 50 | 6 | 0 | -2.639639 | 2.781759 | -0.280492 |
| 51 | 6 | 0 | -2.394162 | 3.863039 | 0.825874 |
| 52 | 8 | 0 | -2.410027 | 5.041927 | 0.438629 |
| 53 | 8 | 0 | -2.279171 | 3.422985 | 2.003747 |
| 54 | 1 | 0 | -3.689879 | 2.492871 | -0.175295 |
| 55 | 1 | 0 | -2.520389 | 3.237786 | -1.259526 |
| 56 | 1 | 0 | -1.941040 | 1.746279 | 1.797573 |

24. **Table S24:** Cartesian Coordinates for the low-energy geometry of the **PEDK** tetrapeptide calculated using the B3LYP method and the 6-311+G(*2df,2pd*) level of theory.

| **Standard Orientation** | | | | | |
| --- | --- | --- | --- | --- | --- |
| Center Number | Atomic Number | Atomic Type | Coordinates (Angstroms) | | |
|  |  |  | X | Y | Z |
| 1 | 6 | 0 | -4.953704 | -3.529357 | -0.313401 |
| 2 | 7 | 0 | -3.850469 | -2.953573 | 0.454357 |
| 3 | 1 | 0 | -3.766495 | -3.324203 | 1.391227 |
| 4 | 6 | 0 | -4.102532 | -1.511304 | 0.455220 |
| 5 | 1 | 0 | -4.983620 | -1.257308 | 1.070355 |
| 6 | 6 | 0 | -4.446048 | -1.235650 | -1.020100 |
| 7 | 6 | 0 | -5.110459 | -2.552508 | -1.498715 |
| 8 | 1 | 0 | -6.159923 | -2.412746 | -1.755534 |
| 9 | 1 | 0 | -4.609259 | -2.936552 | -2.386628 |
| 10 | 1 | 0 | -5.076224 | -0.358238 | -1.134140 |
| 11 | 1 | 0 | -3.528259 | -1.043085 | -1.573500 |
| 12 | 6 | 0 | -2.934805 | -0.735805 | 1.053996 |
| 13 | 8 | 0 | -2.137758 | -1.325345 | 1.828801 |
| 14 | 7 | 0 | -2.860458 | 0.544013 | 0.756400 |
| 15 | 1 | 0 | -3.619478 | 1.072802 | 0.158064 |
| 16 | 6 | 0 | -1.830811 | 1.402442 | 1.298107 |
| 17 | 6 | 0 | -2.259117 | 2.879791 | 1.139685 |
| 18 | 6 | 0 | -2.576891 | 3.360220 | -0.290839 |
| 19 | 6 | 0 | -4.031503 | 3.122330 | -0.782512 |
| 20 | 8 | 0 | -4.541666 | 1.978396 | -0.510544 |
| 21 | 8 | 0 | -4.577509 | 4.034997 | -1.404395 |
| 22 | 1 | 0 | -1.902868 | 2.880303 | -1.006624 |
| 23 | 1 | 0 | -2.390263 | 4.430201 | -0.360071 |
| 24 | 1 | 0 | -1.462159 | 3.501716 | 1.552637 |
| 25 | 1 | 0 | -3.132073 | 3.036076 | 1.777166 |
| 26 | 1 | 0 | -1.710968 | 1.186421 | 2.364919 |
| 27 | 6 | 0 | -0.453301 | 1.127919 | 0.644097 |
| 28 | 8 | 0 | -0.317013 | 0.469642 | -0.377605 |
| 29 | 7 | 0 | 0.595134 | 1.672257 | 1.308926 |
| 30 | 6 | 0 | 1.963037 | 1.755943 | 0.788733 |
| 31 | 6 | 0 | 2.508404 | 0.340612 | 0.490283 |
| 32 | 7 | 0 | 3.370822 | 0.262651 | -0.508583 |
| 33 | 6 | 0 | 4.025444 | -0.960553 | -0.926234 |
| 34 | 6 | 0 | 3.070277 | -2.048744 | -1.494479 |
| 35 | 1 | 0 | 3.543637 | -2.491110 | -2.371998 |
| 36 | 1 | 0 | 2.202617 | -1.501132 | -1.862172 |
| 37 | 6 | 0 | 2.636161 | -3.188420 | -0.553931 |
| 38 | 1 | 0 | 3.338240 | -4.015748 | -0.647463 |
| 39 | 6 | 0 | 1.218813 | -3.712678 | -0.826366 |
| 40 | 1 | 0 | 1.106364 | -3.965884 | -1.884328 |
| 41 | 6 | 0 | 0.082562 | -2.760649 | -0.461013 |
| 42 | 1 | 0 | 0.102171 | -1.838805 | -1.037054 |
| 43 | 7 | 0 | 0.144655 | -2.338151 | 0.970728 |
| 44 | 1 | 0 | 0.322048 | -3.132770 | 1.579162 |
| 45 | 1 | 0 | -0.769392 | -1.902077 | 1.282166 |
| 46 | 1 | 0 | 0.910231 | -1.631209 | 1.113115 |
| 47 | 1 | 0 | -0.887519 | -3.230345 | -0.617653 |
| 48 | 1 | 0 | 1.077619 | -4.651910 | -0.279865 |
| 49 | 1 | 0 | 2.706455 | -2.856817 | 0.480701 |
| 50 | 6 | 0 | 5.042482 | -1.527671 | 0.062571 |
| 51 | 8 | 0 | 5.546030 | -2.620382 | -0.048109 |
| 52 | 8 | 0 | 5.398656 | -0.665016 | 1.028241 |
| 53 | 1 | 0 | 6.084757 | -1.101064 | 1.552937 |
| 54 | 1 | 0 | 4.661398 | -0.645100 | -1.759477 |
| 55 | 1 | 0 | 3.763763 | 1.208634 | -0.921737 |
| 56 | 8 | 0 | 2.178608 | -0.622458 | 1.213412 |
| 57 | 1 | 0 | 2.568201 | 2.107689 | 1.629419 |
| 58 | 6 | 0 | 2.060605 | 2.800853 | -0.345750 |
| 59 | 6 | 0 | 3.489728 | 3.343434 | -0.640769 |
| 60 | 8 | 0 | 3.734345 | 4.505643 | -0.314048 |
| 61 | 8 | 0 | 4.285507 | 2.520612 | -1.211621 |
| 62 | 1 | 0 | 1.455176 | 3.653421 | -0.047648 |
| 63 | 1 | 0 | 1.629804 | 2.378444 | -1.254227 |
| 64 | 1 | 0 | 0.394294 | 2.213692 | 2.131514 |
| 65 | 1 | 0 | -5.890781 | -3.565074 | 0.263960 |
| 66 | 1 | 0 | -4.718934 | -4.548527 | -0.624567 |

25. **Table S25:** Cartesian Coordinates for the low-energy geometry of the **DKGH** tetrapeptide calculated using the B3LYP method and the 6-311+G(*2df,2pd*) level of theory.

| **Standard Orientation** | | | | | |
| --- | --- | --- | --- | --- | --- |
| Center Number | Atomic Number | Atomic Type | Coordinates (Angstroms) | | |
|  |  |  | X | Y | Z |
| 1 | 6 | 0 | -2.237798 | -0.892019 | -0.905883 |
| 2 | 7 | 0 | -2.137856 | 0.419566 | -0.234206 |
| 3 | 6 | 0 | -3.046219 | 1.318693 | 0.139791 |
| 4 | 6 | 0 | -2.476697 | 2.651249 | 0.692243 |
| 5 | 7 | 0 | -3.564594 | 3.618030 | 0.762223 |
| 6 | 1 | 0 | -4.426495 | 3.173747 | 1.043110 |
| 7 | 1 | 0 | -3.348904 | 4.363915 | 1.409464 |
| 8 | 1 | 0 | -2.075590 | 2.384935 | 1.681869 |
| 9 | 6 | 0 | -1.333866 | 3.300724 | -0.121588 |
| 10 | 6 | 0 | 0.060180 | 2.759844 | 0.110686 |
| 11 | 8 | 0 | 1.001454 | 3.638963 | -0.100566 |
| 12 | 8 | 0 | 0.268534 | 1.598339 | 0.453006 |
| 13 | 1 | 0 | -1.320932 | 4.363707 | 0.112826 |
| 14 | 1 | 0 | -1.541964 | 3.253820 | -1.194645 |
| 15 | 8 | 0 | -4.281832 | 1.162719 | 0.125077 |
| 16 | 1 | 0 | -1.183060 | 0.767715 | -0.128002 |
| 17 | 6 | 0 | -3.628909 | -1.527041 | -0.985869 |
| 18 | 1 | 0 | -3.517113 | -2.396837 | -1.632720 |
| 19 | 1 | 0 | -4.281987 | -0.826416 | -1.502679 |
| 20 | 6 | 0 | -4.216892 | -1.978564 | 0.351450 |
| 21 | 1 | 0 | -3.616754 | -2.797005 | 0.749160 |
| 22 | 6 | 0 | -5.667092 | -2.469572 | 0.252515 |
| 23 | 1 | 0 | -5.722031 | -3.361445 | -0.375644 |
| 24 | 6 | 0 | -6.651608 | -1.468050 | -0.336281 |
| 25 | 1 | 0 | -6.452826 | -1.264013 | -1.385863 |
| 26 | 7 | 0 | -6.563134 | -0.135508 | 0.365445 |
| 27 | 1 | 0 | -6.744518 | -0.235113 | 1.363577 |
| 28 | 1 | 0 | -7.246531 | 0.523080 | -0.005515 |
| 29 | 1 | 0 | -5.600224 | 0.305861 | 0.254318 |
| 30 | 1 | 0 | -7.677498 | -1.819679 | -0.245501 |
| 31 | 1 | 0 | -6.011070 | -2.778453 | 1.243244 |
| 32 | 1 | 0 | -4.152491 | -1.171634 | 1.084439 |
| 33 | 6 | 0 | -1.212675 | -1.939140 | -0.390359 |
| 34 | 8 | 0 | -1.328251 | -3.086753 | -0.778038 |
| 35 | 7 | 0 | -0.220326 | -1.523266 | 0.435690 |
| 36 | 6 | 0 | 0.951633 | -2.345359 | 0.680132 |
| 37 | 1 | 0 | 0.720863 | -3.360306 | 0.360552 |
| 38 | 1 | 0 | 1.183322 | -2.360599 | 1.745407 |
| 39 | 6 | 0 | 2.132399 | -1.821479 | -0.152723 |
| 40 | 8 | 0 | 1.986374 | -1.462092 | -1.302306 |
| 41 | 7 | 0 | 3.328368 | -1.753935 | 0.494000 |
| 42 | 1 | 0 | 3.452337 | -2.075440 | 1.442383 |
| 43 | 6 | 0 | 4.476021 | -1.137682 | -0.132544 |
| 44 | 1 | 0 | 4.628711 | -1.565267 | -1.128028 |
| 45 | 6 | 0 | 4.271445 | 0.350264 | -0.271205 |
| 46 | 6 | 0 | 3.348188 | 1.154884 | 0.323496 |
| 47 | 7 | 0 | 3.395242 | 2.416127 | -0.220045 |
| 48 | 6 | 0 | 4.363177 | 2.390846 | -1.110946 |
| 49 | 7 | 0 | 4.924374 | 1.158617 | -1.175836 |
| 50 | 1 | 0 | 5.698096 | 0.892398 | -1.762244 |
| 51 | 1 | 0 | 4.681034 | 3.213278 | -1.728245 |
| 52 | 1 | 0 | 1.947001 | 3.223462 | -0.085525 |
| 53 | 1 | 0 | 2.610489 | 0.894584 | 1.056771 |
| 54 | 6 | 0 | 5.709654 | -1.459985 | 0.709904 |
| 55 | 8 | 0 | 5.679868 | -1.978027 | 1.791200 |
| 56 | 8 | 0 | 6.830294 | -1.066564 | 0.083787 |
| 57 | 1 | 0 | 7.588381 | -1.260669 | 0.655284 |
| 58 | 1 | 0 | -0.106222 | -0.540345 | 0.625097 |
| 59 | 1 | 0 | -1.915697 | -0.723045 | -1.937979 |

26. **Table S26:** Cartesian Coordinates for the low-energy geometry of the **EAEE+EA** hexapeptide calculated using the B3LYP method and the 6-311+G(*2df,2pd*) level of theory.

| **Standard Orientation** | | | | | |
| --- | --- | --- | --- | --- | --- |
| Center Number | Atomic Number | Atomic Type | Coordinates (Angstroms) | | |
|  |  |  | X | Y | Z |
| 1 | 6 | 0 | 2.553524 | -0.355534 | 0.975038 |
| 2 | 7 | 0 | 1.344759 | -0.883399 | 0.367472 |
| 3 | 1 | 0 | 0.570000 | -0.258430 | 0.197069 |
| 4 | 6 | 0 | 1.254055 | -2.178376 | -0.048769 |
| 5 | 6 | 0 | -0.062016 | -2.540170 | -0.782598 |
| 6 | 7 | 0 | -1.243244 | -1.947225 | -0.140903 |
| 7 | 6 | 0 | -1.690550 | -2.260563 | 1.094701 |
| 8 | 6 | 0 | -3.020206 | -1.603340 | 1.544782 |
| 9 | 7 | 0 | -3.522228 | -0.572531 | 0.650213 |
| 10 | 6 | 0 | -3.097753 | 0.714098 | 0.747477 |
| 11 | 6 | 0 | -3.765225 | 1.732692 | -0.210898 |
| 12 | 7 | 0 | -2.774404 | 2.645465 | -0.778994 |
| 13 | 6 | 0 | -1.628008 | 2.215746 | -1.318474 |
| 14 | 6 | 0 | -0.680730 | 3.242467 | -1.972638 |
| 15 | 7 | 0 | 0.555260 | 3.272678 | -1.185295 |
| 16 | 1 | 0 | 1.264544 | 3.812079 | -1.666990 |
| 17 | 1 | 0 | 0.939531 | 2.341146 | -1.068321 |
| 18 | 1 | 0 | -0.516971 | 2.800973 | -2.969719 |
| 19 | 6 | 0 | -1.210304 | 4.664199 | -2.220341 |
| 20 | 6 | 0 | -1.394999 | 5.599977 | -1.001420 |
| 21 | 1 | 0 | -0.683271 | 6.424335 | -1.036588 |
| 22 | 1 | 0 | -1.186281 | 5.044065 | -0.085237 |
| 23 | 6 | 0 | -2.829548 | 6.174763 | -0.890874 |
| 24 | 8 | 0 | -3.749917 | 5.305452 | -0.906809 |
| 25 | 8 | 0 | -2.961819 | 7.414779 | -0.791621 |
| 26 | 1 | 0 | -2.157956 | 4.578956 | -2.755759 |
| 27 | 1 | 0 | -0.509082 | 5.131440 | -2.921394 |
| 28 | 8 | 0 | -1.287602 | 1.025744 | -1.342824 |
| 29 | 1 | 0 | -3.059765 | 3.639447 | -0.827251 |
| 30 | 1 | 0 | -4.383684 | 2.355831 | 0.439758 |
| 31 | 6 | 0 | -4.661540 | 1.145529 | -1.307058 |
| 32 | 1 | 0 | -5.095259 | 1.967820 | -1.878190 |
| 33 | 1 | 0 | -5.472821 | 0.545232 | -0.895364 |
| 34 | 1 | 0 | -4.077606 | 0.524803 | -1.986399 |
| 35 | 8 | 0 | -2.256957 | 1.083507 | 1.555776 |
| 36 | 1 | 0 | -4.434204 | -0.786994 | 0.232149 |
| 37 | 6 | 0 | -4.072626 | -2.710537 | 1.738829 |
| 38 | 6 | 0 | -5.395674 | -2.272976 | 2.388827 |
| 39 | 6 | 0 | -6.544018 | -1.813254 | 1.441031 |
| 40 | 8 | 0 | -7.699959 | -1.883493 | 1.916376 |
| 41 | 8 | 0 | -6.211922 | -1.403196 | 0.294656 |
| 42 | 1 | 0 | -5.798756 | -3.091564 | 2.988870 |
| 43 | 1 | 0 | -5.217208 | -1.450948 | 3.091039 |
| 44 | 1 | 0 | -4.283526 | -3.168222 | 0.769369 |
| 45 | 1 | 0 | -3.593103 | -3.472733 | 2.352506 |
| 46 | 1 | 0 | -2.789139 | -1.150181 | 2.511592 |
| 47 | 8 | 0 | -1.121910 | -3.042443 | 1.847757 |
| 48 | 1 | 0 | -1.712574 | -1.195694 | -0.625659 |
| 49 | 6 | 0 | -0.199086 | -4.046850 | -1.096604 |
| 50 | 6 | 0 | 0.594222 | -4.491717 | -2.330637 |
| 51 | 6 | 0 | 0.352705 | -5.950705 | -2.843864 |
| 52 | 8 | 0 | 1.033420 | -6.266397 | -3.856883 |
| 53 | 8 | 0 | -0.477864 | -6.660454 | -2.229973 |
| 54 | 1 | 0 | 0.366989 | -3.830931 | -3.175863 |
| 55 | 1 | 0 | 1.664872 | -4.384965 | -2.151183 |
| 56 | 1 | 0 | -1.256049 | -4.252482 | -1.266326 |
| 57 | 1 | 0 | 0.093501 | -4.631023 | -0.227644 |
| 58 | 1 | 0 | 0.011398 | -2.009911 | -1.736662 |
| 59 | 8 | 0 | 2.176279 | -2.969683 | 0.080582 |
| 60 | 1 | 0 | 3.043813 | -1.201709 | 1.455756 |
| 61 | 6 | 0 | 3.504046 | 0.248310 | -0.086069 |
| 62 | 8 | 0 | 3.185516 | 1.240395 | -0.730857 |
| 63 | 7 | 0 | 4.691861 | -0.390716 | -0.189988 |
| 64 | 6 | 0 | 5.876889 | 0.031398 | -0.933891 |
| 65 | 1 | 0 | 6.522995 | -0.851474 | -0.947899 |
| 66 | 6 | 0 | 6.749885 | 1.118286 | -0.204414 |
| 67 | 8 | 0 | 7.695900 | 1.571944 | -0.869051 |
| 68 | 8 | 0 | 6.466444 | 1.398041 | 0.996447 |
| 69 | 6 | 0 | 5.577230 | 0.395820 | -2.389975 |
| 70 | 1 | 0 | 5.068799 | -0.430449 | -2.890423 |
| 71 | 1 | 0 | 6.517479 | 0.607435 | -2.894951 |
| 72 | 1 | 0 | 4.935897 | 1.271060 | -2.445372 |
| 73 | 1 | 0 | 4.798863 | -1.196564 | 0.400856 |
| 74 | 6 | 0 | 2.212758 | 0.711513 | 2.025302 |
| 75 | 6 | 0 | 3.434786 | 1.417938 | 2.645498 |
| 76 | 1 | 0 | 3.059599 | 2.193057 | 3.315453 |
| 77 | 1 | 0 | 4.025310 | 1.903201 | 1.869639 |
| 78 | 6 | 0 | 4.314818 | 0.509449 | 3.497398 |
| 79 | 8 | 0 | 3.942836 | 0.088564 | 4.577000 |
| 80 | 8 | 0 | 5.511605 | 0.195040 | 3.021392 |
| 81 | 1 | 0 | 5.781796 | 0.663822 | 2.150865 |
| 82 | 1 | 0 | 1.617113 | 0.242732 | 2.808388 |
| 83 | 1 | 0 | 1.592516 | 1.475566 | 1.554474 |

27. **Table S27:** Cartesian Coordinates for the low-energy geometry of the **EA+EEEA** hexapeptide calculated using the B3LYP method and the 6-311+G(*2df,2pd*) level of theory.

| **Standard Orientation** | | | | | |
| --- | --- | --- | --- | --- | --- |
| Center Number | Atomic Number | Atomic Type | Coordinates (Angstroms) | | |
|  |  |  | X | Y | Z |
| 1 | 6 | 0 | -2.197851 | 1.774344 | -0.220429 |
| 2 | 6 | 0 | -2.967658 | 3.120039 | -0.145732 |
| 3 | 6 | 0 | -2.634530 | 4.068983 | -1.303902 |
| 4 | 6 | 0 | -3.505991 | 5.360512 | -1.442577 |
| 5 | 8 | 0 | -3.248389 | 6.055937 | -2.461777 |
| 6 | 8 | 0 | -4.363283 | 5.587173 | -0.557461 |
| 7 | 1 | 0 | -1.591153 | 4.384520 | -1.239364 |
| 8 | 1 | 0 | -2.723959 | 3.534924 | -2.257013 |
| 9 | 1 | 0 | -4.034223 | 2.901413 | -0.163213 |
| 10 | 1 | 0 | -2.774200 | 3.606920 | 0.806655 |
| 11 | 7 | 0 | -3.021346 | 0.574728 | 0.014234 |
| 12 | 1 | 0 | -3.089036 | -0.116561 | -0.727269 |
| 13 | 6 | 0 | -3.770456 | 0.331268 | 1.111214 |
| 14 | 6 | 0 | -4.486749 | -1.052181 | 1.204172 |
| 15 | 7 | 0 | -5.319719 | -1.548287 | 0.092106 |
| 16 | 6 | 0 | -4.884488 | -1.882855 | -1.122941 |
| 17 | 6 | 0 | -5.797546 | -2.688420 | -2.071931 |
| 18 | 7 | 0 | -5.910392 | -1.917163 | -3.319588 |
| 19 | 1 | 0 | -6.182226 | -2.526512 | -4.081698 |
| 20 | 1 | 0 | -4.998955 | -1.531001 | -3.541262 |
| 21 | 1 | 0 | -5.192909 | -3.597751 | -2.225514 |
| 22 | 6 | 0 | -7.166736 | -3.172482 | -1.587587 |
| 23 | 6 | 0 | -8.257181 | -2.119744 | -1.290552 |
| 24 | 1 | 0 | -9.098239 | -2.225400 | -1.974968 |
| 25 | 1 | 0 | -7.852441 | -1.116514 | -1.439988 |
| 26 | 6 | 0 | -8.770535 | -2.219086 | 0.168580 |
| 27 | 8 | 0 | -7.856882 | -2.190206 | 1.046213 |
| 28 | 8 | 0 | -9.999180 | -2.326006 | 0.353476 |
| 29 | 1 | 0 | -7.003697 | -3.778680 | -0.694530 |
| 30 | 1 | 0 | -7.532679 | -3.862777 | -2.357106 |
| 31 | 8 | 0 | -3.750660 | -1.627809 | -1.561320 |
| 32 | 1 | 0 | -6.290445 | -1.797364 | 0.385569 |
| 33 | 1 | 0 | -5.195360 | -0.895274 | 2.012907 |
| 34 | 6 | 0 | -3.469918 | -2.115122 | 1.651112 |
| 35 | 1 | 0 | -3.991989 | -3.029237 | 1.938862 |
| 36 | 1 | 0 | -2.905689 | -1.751330 | 2.509871 |
| 37 | 1 | 0 | -2.773645 | -2.352182 | 0.847578 |
| 38 | 8 | 0 | -3.837738 | 1.066807 | 2.084716 |
| 39 | 6 | 0 | -0.851602 | 1.773039 | 0.558647 |
| 40 | 8 | 0 | -0.194909 | 2.793638 | 0.672870 |
| 41 | 7 | 0 | -0.428866 | 0.559429 | 1.011617 |
| 42 | 6 | 0 | 0.933255 | 0.250896 | 1.442946 |
| 43 | 6 | 0 | 1.328963 | 0.958569 | 2.764239 |
| 44 | 6 | 0 | 2.501909 | 0.345243 | 3.548899 |
| 45 | 6 | 0 | 2.438778 | 0.477255 | 5.118525 |
| 46 | 8 | 0 | 1.303924 | 0.489014 | 5.638578 |
| 47 | 8 | 0 | 3.561515 | 0.506762 | 5.682720 |
| 48 | 1 | 0 | 2.560417 | -0.733513 | 3.358847 |
| 49 | 1 | 0 | 3.471986 | 0.764499 | 3.267475 |
| 50 | 1 | 0 | 0.458429 | 0.895123 | 3.417224 |
| 51 | 1 | 0 | 1.496476 | 2.017864 | 2.571818 |
| 52 | 6 | 0 | 1.894830 | 0.324005 | 0.209658 |
| 53 | 8 | 0 | 1.462605 | 0.194257 | -0.920837 |
| 54 | 7 | 0 | 3.217954 | 0.457936 | 0.519264 |
| 55 | 6 | 0 | 4.396895 | 0.357651 | -0.342386 |
| 56 | 1 | 0 | 5.151679 | 0.977802 | 0.149753 |
| 57 | 6 | 0 | 4.939691 | -1.095849 | -0.404365 |
| 58 | 8 | 0 | 4.298347 | -2.003735 | -0.905404 |
| 59 | 7 | 0 | 6.180738 | -1.238453 | 0.138827 |
| 60 | 6 | 0 | 7.051868 | -2.411698 | 0.043904 |
| 61 | 1 | 0 | 7.861272 | -2.208475 | 0.752767 |
| 62 | 6 | 0 | 7.798745 | -2.557426 | -1.332675 |
| 63 | 8 | 0 | 8.357736 | -3.645209 | -1.532507 |
| 64 | 8 | 0 | 7.832061 | -1.539533 | -2.088448 |
| 65 | 6 | 0 | 6.368307 | -3.702015 | 0.496781 |
| 66 | 1 | 0 | 5.959013 | -3.580957 | 1.501694 |
| 67 | 1 | 0 | 7.103334 | -4.504016 | 0.495719 |
| 68 | 1 | 0 | 5.554431 | -3.965196 | -0.172615 |
| 69 | 1 | 0 | 6.643379 | -0.382004 | 0.389589 |
| 70 | 6 | 0 | 4.202382 | 0.911641 | -1.761631 |
| 71 | 6 | 0 | 5.469146 | 0.822688 | -2.636149 |
| 72 | 1 | 0 | 5.220436 | 1.242111 | -3.612654 |
| 73 | 1 | 0 | 5.758348 | -0.217087 | -2.783514 |
| 74 | 6 | 0 | 6.650931 | 1.632225 | -2.111837 |
| 75 | 8 | 0 | 6.631372 | 2.847534 | -2.077063 |
| 76 | 8 | 0 | 7.710022 | 0.954467 | -1.686222 |
| 77 | 1 | 0 | 7.664678 | -0.066581 | -1.825196 |
| 78 | 1 | 0 | 3.878112 | 1.949074 | -1.682165 |
| 79 | 1 | 0 | 3.402763 | 0.357867 | -2.243707 |
| 80 | 1 | 0 | 3.424760 | 0.525037 | 1.502506 |
| 81 | 1 | 0 | 0.914109 | -0.820726 | 1.670711 |
| 82 | 1 | 0 | -0.997900 | -0.221511 | 0.737528 |
| 83 | 1 | 0 | -1.862570 | 1.652560 | -1.254714 |

28. **Table S28:** Cartesian Coordinates for the low-energy geometry of the **EGED+EA** hexapeptide calculated using the B3LYP method and the 6-311+G(*2df,2pd*) level of theory.

| **Standard Orientation** | | | | | |
| --- | --- | --- | --- | --- | --- |
| Center Number | Atomic Number | Atomic Type | Coordinates (Angstroms) | | |
|  |  |  | X | Y | Z |
| 1 | 6 | 0 | -2.697149 | 0.842304 | 0.106045 |
| 2 | 7 | 0 | -3.933658 | 0.080246 | 0.033533 |
| 3 | 1 | 0 | -4.755123 | 0.584948 | -0.332130 |
| 4 | 6 | 0 | -4.002351 | -1.111177 | 0.643192 |
| 5 | 6 | 0 | -5.317239 | -1.926591 | 0.650356 |
| 6 | 7 | 0 | -6.555887 | -1.275647 | 0.248770 |
| 7 | 6 | 0 | -7.050125 | -1.189319 | -1.007322 |
| 8 | 6 | 0 | -8.524927 | -0.731934 | -1.098884 |
| 9 | 1 | 0 | -8.579540 | -0.280820 | -2.089499 |
| 10 | 7 | 0 | -9.372239 | -1.953463 | -1.139767 |
| 11 | 1 | 0 | -10.336448 | -1.650036 | -1.239507 |
| 12 | 1 | 0 | -9.335057 | -2.373708 | -0.214255 |
| 13 | 6 | 0 | -9.014197 | 0.317381 | -0.067773 |
| 14 | 1 | 0 | -9.821477 | 0.890042 | -0.527967 |
| 15 | 1 | 0 | -8.203265 | 1.018334 | 0.145168 |
| 16 | 6 | 0 | -9.565431 | -0.287694 | 1.251278 |
| 17 | 8 | 0 | -10.750115 | -0.025558 | 1.554582 |
| 18 | 8 | 0 | -8.766131 | -1.026677 | 1.902237 |
| 19 | 8 | 0 | -6.431561 | -1.470209 | -2.028485 |
| 20 | 1 | 0 | -7.250291 | -1.115399 | 0.997960 |
| 21 | 1 | 0 | -5.451628 | -2.137119 | 1.713546 |
| 22 | 6 | 0 | -5.072265 | -3.275504 | -0.048190 |
| 23 | 1 | 0 | -4.928253 | -3.134191 | -1.115881 |
| 24 | 1 | 0 | -5.937735 | -3.924535 | 0.099746 |
| 25 | 1 | 0 | -4.189521 | -3.744191 | 0.383131 |
| 26 | 8 | 0 | -3.035268 | -1.610460 | 1.228335 |
| 27 | 1 | 0 | -2.433961 | 0.983422 | 1.160705 |
| 28 | 6 | 0 | -2.851751 | 2.224128 | -0.555047 |
| 29 | 6 | 0 | -3.788510 | 3.220370 | 0.149167 |
| 30 | 6 | 0 | -5.275208 | 3.221966 | -0.315940 |
| 31 | 8 | 0 | -5.866158 | 4.322289 | -0.238605 |
| 32 | 8 | 0 | -5.738626 | 2.119844 | -0.710205 |
| 33 | 1 | 0 | -3.789568 | 3.035364 | 1.229499 |
| 34 | 1 | 0 | -3.408990 | 4.235275 | 0.018121 |
| 35 | 1 | 0 | -3.177835 | 2.071714 | -1.585070 |
| 36 | 1 | 0 | -1.848319 | 2.655755 | -0.598822 |
| 37 | 6 | 0 | -1.506438 | 0.122915 | -0.570346 |
| 38 | 8 | 0 | -1.583688 | -0.407713 | -1.667967 |
| 39 | 7 | 0 | -0.327171 | 0.242899 | 0.096775 |
| 40 | 1 | 0 | -0.297417 | 0.561142 | 1.053290 |
| 41 | 6 | 0 | 0.913618 | -0.297253 | -0.404194 |
| 42 | 6 | 0 | 2.071274 | 0.092722 | 0.527479 |
| 43 | 8 | 0 | 1.848973 | 0.589684 | 1.631439 |
| 44 | 7 | 0 | 3.300540 | -0.163800 | 0.049115 |
| 45 | 6 | 0 | 4.542590 | -0.028305 | 0.813237 |
| 46 | 6 | 0 | 5.610084 | -0.957526 | 0.171994 |
| 47 | 7 | 0 | 6.629250 | -1.210232 | 0.924620 |
| 48 | 6 | 0 | 7.808834 | -1.980264 | 0.494443 |
| 49 | 6 | 0 | 7.598707 | -3.342223 | -0.201195 |
| 50 | 8 | 0 | 8.293919 | -4.303858 | 0.089716 |
| 51 | 8 | 0 | 6.677134 | -3.442714 | -1.147639 |
| 52 | 1 | 0 | 6.130731 | -2.566348 | -1.208054 |
| 53 | 1 | 0 | 8.332491 | -2.250379 | 1.411908 |
| 54 | 6 | 0 | 8.836258 | -1.137155 | -0.318024 |
| 55 | 1 | 0 | 9.565854 | -1.816886 | -0.761142 |
| 56 | 1 | 0 | 8.308034 | -0.625074 | -1.157998 |
| 57 | 6 | 0 | 9.625204 | -0.112688 | 0.493459 |
| 58 | 8 | 0 | 10.494523 | 0.571117 | -0.394405 |
| 59 | 1 | 0 | 9.935358 | 1.016260 | -1.084006 |
| 60 | 8 | 0 | 8.685510 | 0.916188 | 0.944048 |
| 61 | 1 | 0 | 7.912089 | 0.431644 | 1.271127 |
| 62 | 8 | 0 | 5.333554 | -1.352898 | -1.026877 |
| 63 | 1 | 0 | 4.357638 | -0.374046 | 1.828852 |
| 64 | 6 | 0 | 4.999846 | 1.440461 | 0.911275 |
| 65 | 1 | 0 | 4.147149 | 2.028050 | 1.254423 |
| 66 | 1 | 0 | 5.766936 | 1.502968 | 1.681370 |
| 67 | 6 | 0 | 5.558410 | 2.005846 | -0.400519 |
| 68 | 1 | 0 | 4.958351 | 1.715715 | -1.261394 |
| 69 | 1 | 0 | 6.568494 | 1.600664 | -0.562033 |
| 70 | 6 | 0 | 5.712172 | 3.493539 | -0.454390 |
| 71 | 8 | 0 | 5.656100 | 4.183327 | -1.450512 |
| 72 | 8 | 0 | 5.978114 | 4.061370 | 0.753792 |
| 73 | 1 | 0 | 6.124571 | 4.999049 | 0.570042 |
| 74 | 1 | 0 | 3.438624 | -0.664370 | -0.822656 |
| 75 | 1 | 0 | 0.870926 | -1.390357 | -0.469160 |
| 76 | 1 | 0 | 1.102654 | 0.063239 | -1.416869 |

29. **Table S29:** Cartesian Coordinates for the low-energy geometry of the **EG+EDEA** hexapeptide calculated using the B3LYP method and the 6-311+G(*2df,2pd*) level of theory.

| **Standard Orientation** | | | | | |
| --- | --- | --- | --- | --- | --- |
| Center Number | Atomic Number | Atomic Type | Coordinates (Angstroms) | | |
|  |  |  | X | Y | Z |
| 1 | 6 | 0 | -1.614630 | 1.860259 | -0.634929 |
| 2 | 7 | 0 | -2.856774 | 1.840114 | 0.118708 |
| 3 | 1 | 0 | -3.197298 | 2.796183 | 0.304915 |
| 4 | 6 | 0 | -3.748117 | 0.835701 | 0.064703 |
| 5 | 6 | 0 | -5.044768 | 1.141752 | 0.827920 |
| 6 | 7 | 0 | -6.080815 | 0.177554 | 0.525411 |
| 7 | 6 | 0 | -7.230455 | 0.114529 | 1.218705 |
| 8 | 6 | 0 | -8.224906 | -0.997223 | 0.826441 |
| 9 | 7 | 0 | -7.676992 | -1.938325 | -0.149098 |
| 10 | 1 | 0 | -8.457999 | -2.413778 | -0.610823 |
| 11 | 1 | 0 | -7.096236 | -2.635656 | 0.299752 |
| 12 | 1 | 0 | -8.459528 | -1.473192 | 1.790006 |
| 13 | 6 | 0 | -9.538691 | -0.377822 | 0.303927 |
| 14 | 6 | 0 | -10.824546 | -1.169121 | 0.591945 |
| 15 | 6 | 0 | -11.161903 | -2.447645 | -0.237342 |
| 16 | 8 | 0 | -12.296410 | -2.932015 | -0.007056 |
| 17 | 8 | 0 | -10.297511 | -2.871870 | -1.045803 |
| 18 | 1 | 0 | -10.846153 | -1.461352 | 1.647238 |
| 19 | 1 | 0 | -11.682499 | -0.501605 | 0.464997 |
| 20 | 1 | 0 | -9.430532 | -0.221714 | -0.772647 |
| 21 | 1 | 0 | -9.637849 | 0.603654 | 0.770922 |
| 22 | 8 | 0 | -7.500654 | 0.883886 | 2.144674 |
| 23 | 1 | 0 | -5.909169 | -0.508126 | -0.198517 |
| 24 | 1 | 0 | -5.375250 | 2.157119 | 0.595758 |
| 25 | 1 | 0 | -4.830040 | 1.128007 | 1.898940 |
| 26 | 8 | 0 | -3.579430 | -0.240477 | -0.502723 |
| 27 | 1 | 0 | -1.799875 | 1.450995 | -1.632579 |
| 28 | 6 | 0 | -1.116813 | 3.325465 | -0.754610 |
| 29 | 6 | 0 | -2.186477 | 4.405495 | -1.108947 |
| 30 | 8 | 0 | -3.179871 | 4.464489 | -0.324372 |
| 31 | 8 | 0 | -1.950734 | 5.138844 | -2.088666 |
| 32 | 1 | 0 | -0.696447 | 3.608570 | 0.214113 |
| 33 | 1 | 0 | -0.317532 | 3.359352 | -1.491212 |
| 34 | 6 | 0 | -0.479978 | 1.039932 | 0.036081 |
| 35 | 8 | 0 | -0.342208 | 0.989973 | 1.249074 |
| 36 | 7 | 0 | 0.387357 | 0.487796 | -0.846734 |
| 37 | 1 | 0 | 0.182627 | 0.521938 | -1.833447 |
| 38 | 6 | 0 | 1.660720 | -0.163698 | -0.520188 |
| 39 | 1 | 0 | 2.057039 | 0.287439 | 0.385733 |
| 40 | 6 | 0 | 2.560093 | 0.106652 | -1.740819 |
| 41 | 8 | 0 | 2.065800 | 0.057664 | -2.870098 |
| 42 | 7 | 0 | 3.869607 | 0.421621 | -1.597854 |
| 43 | 6 | 0 | 4.771294 | 0.430313 | -0.446311 |
| 44 | 1 | 0 | 4.239163 | 0.029237 | 0.410602 |
| 45 | 6 | 0 | 5.965427 | -0.485195 | -0.786570 |
| 46 | 8 | 0 | 6.693822 | -0.206056 | -1.743719 |
| 47 | 7 | 0 | 6.131400 | -1.509367 | 0.057469 |
| 48 | 6 | 0 | 7.259214 | -2.433934 | 0.034125 |
| 49 | 1 | 0 | 7.057027 | -3.130341 | 0.852571 |
| 50 | 6 | 0 | 8.631615 | -1.778112 | 0.428827 |
| 51 | 8 | 0 | 9.651227 | -2.417056 | 0.117379 |
| 52 | 8 | 0 | 8.601107 | -0.697786 | 1.090650 |
| 53 | 6 | 0 | 7.332416 | -3.249486 | -1.260953 |
| 54 | 1 | 0 | 6.389826 | -3.776575 | -1.419506 |
| 55 | 1 | 0 | 8.148754 | -3.966285 | -1.190950 |
| 56 | 1 | 0 | 7.515022 | -2.594054 | -2.110384 |
| 57 | 1 | 0 | 5.309256 | -1.752318 | 0.624214 |
| 58 | 6 | 0 | 5.287210 | 1.864015 | -0.185749 |
| 59 | 6 | 0 | 6.577040 | 1.986005 | 0.650172 |
| 60 | 1 | 0 | 6.895790 | 3.029858 | 0.599337 |
| 61 | 1 | 0 | 7.363746 | 1.375526 | 0.211001 |
| 62 | 6 | 0 | 6.433394 | 1.685218 | 2.139967 |
| 63 | 8 | 0 | 5.820041 | 2.438317 | 2.875885 |
| 64 | 8 | 0 | 7.034720 | 0.606625 | 2.602820 |
| 65 | 1 | 0 | 7.567946 | 0.063214 | 1.910895 |
| 66 | 1 | 0 | 4.486737 | 2.444814 | 0.275446 |
| 67 | 1 | 0 | 5.496724 | 2.313799 | -1.158599 |
| 68 | 1 | 0 | 4.344257 | 0.509321 | -2.487309 |
| 69 | 6 | 0 | 1.474521 | -1.681097 | -0.288624 |
| 70 | 6 | 0 | 2.690560 | -2.492061 | 0.243161 |
| 71 | 8 | 0 | 2.746761 | -3.687383 | -0.103235 |
| 72 | 8 | 0 | 3.487882 | -1.876544 | 1.010279 |
| 73 | 1 | 0 | 0.680813 | -1.780303 | 0.455169 |
| 74 | 1 | 0 | 1.115524 | -2.143998 | -1.207231 |

30. **Table S30:** Cartesian Coordinates for the low-energy geometry of the **EAQG+PK** hexapeptide calculated using the B3LYP method and the 6-311+G(*2df,2pd*) level of theory.

| **Standard Orientation** | | | | | |
| --- | --- | --- | --- | --- | --- |
| Center Number | Atomic Number | Atomic Type | Coordinates (Angstroms) | | |
|  |  |  | X | Y | Z |
| 1 | 6 | 0 | -3.263688 | -2.177616 | -1.651357 |
| 2 | 6 | 0 | -3.648617 | -3.641872 | -1.360900 |
| 3 | 6 | 0 | -2.864984 | -3.983572 | -0.088109 |
| 4 | 6 | 0 | -1.534185 | -3.255359 | -0.280135 |
| 5 | 7 | 0 | -1.918001 | -2.058347 | -1.063533 |
| 6 | 6 | 0 | -1.121002 | -1.016998 | -1.385202 |
| 7 | 6 | 0 | 0.325917 | -1.049189 | -0.908368 |
| 8 | 7 | 0 | 1.142049 | -1.817421 | -1.849140 |
| 9 | 6 | 0 | 2.501451 | -1.795591 | -1.995308 |
| 10 | 6 | 0 | 3.289658 | -0.565463 | -1.518708 |
| 11 | 7 | 0 | 4.521372 | -1.025652 | -0.888628 |
| 12 | 6 | 0 | 4.838968 | -0.986181 | 0.449229 |
| 13 | 6 | 0 | 4.319117 | 0.221447 | 1.251488 |
| 14 | 7 | 0 | 3.310459 | -0.113256 | 2.256743 |
| 15 | 6 | 0 | 2.404982 | -1.105992 | 2.133231 |
| 16 | 6 | 0 | 1.335613 | -1.175055 | 3.240839 |
| 17 | 7 | 0 | 0.320580 | -2.144200 | 2.813898 |
| 18 | 1 | 0 | -0.231374 | -2.439562 | 3.609379 |
| 19 | 1 | 0 | 0.788860 | -2.956214 | 2.428525 |
| 20 | 1 | 0 | 1.888453 | -1.532176 | 4.122062 |
| 21 | 6 | 0 | 0.674761 | 0.156830 | 3.653486 |
| 22 | 6 | 0 | -0.044927 | 0.939777 | 2.551908 |
| 23 | 1 | 0 | -0.844190 | 1.533685 | 3.003054 |
| 24 | 1 | 0 | -0.545314 | 0.261612 | 1.862442 |
| 25 | 6 | 0 | 0.804783 | 1.943823 | 1.746002 |
| 26 | 8 | 0 | 1.897271 | 2.328484 | 2.212864 |
| 27 | 8 | 0 | 0.302319 | 2.328989 | 0.643864 |
| 28 | 1 | 0 | 1.412126 | 0.804690 | 4.129128 |
| 29 | 1 | 0 | -0.043477 | -0.098434 | 4.437217 |
| 30 | 8 | 0 | 2.446765 | -1.960781 | 1.252773 |
| 31 | 1 | 0 | 2.986737 | 0.714294 | 2.751316 |
| 32 | 1 | 0 | 3.869388 | 0.934421 | 0.562016 |
| 33 | 6 | 0 | 5.501402 | 0.903915 | 1.940500 |
| 34 | 1 | 0 | 5.154288 | 1.765190 | 2.511950 |
| 35 | 1 | 0 | 6.226438 | 1.248901 | 1.202573 |
| 36 | 1 | 0 | 5.998578 | 0.201931 | 2.605815 |
| 37 | 8 | 0 | 5.639860 | -1.761261 | 0.931956 |
| 38 | 1 | 0 | 4.911990 | -1.841523 | -1.346172 |
| 39 | 1 | 0 | 2.709818 | -0.015107 | -0.787623 |
| 40 | 6 | 0 | 3.566761 | 0.324085 | -2.756658 |
| 41 | 6 | 0 | 3.889795 | 1.792623 | -2.428068 |
| 42 | 6 | 0 | 2.646164 | 2.493261 | -1.911021 |
| 43 | 7 | 0 | 2.659259 | 3.008479 | -0.682104 |
| 44 | 1 | 0 | 3.467663 | 2.895255 | -0.095347 |
| 45 | 1 | 0 | 1.764054 | 3.094366 | -0.172318 |
| 46 | 8 | 0 | 1.630913 | 2.525417 | -2.630812 |
| 47 | 1 | 0 | 4.713766 | 1.862797 | -1.717411 |
| 48 | 1 | 0 | 4.199219 | 2.295407 | -3.345512 |
| 49 | 1 | 0 | 2.697671 | 0.313210 | -3.414693 |
| 50 | 1 | 0 | 4.392668 | -0.121909 | -3.309905 |
| 51 | 8 | 0 | 3.081237 | -2.696976 | -2.576878 |
| 52 | 1 | 0 | 0.757328 | -2.692460 | -2.175537 |
| 53 | 1 | 0 | 0.401359 | -1.458339 | 0.098692 |
| 54 | 1 | 0 | 0.654484 | -0.015645 | -0.879669 |
| 55 | 8 | 0 | -1.556555 | -0.069057 | -2.048195 |
| 56 | 1 | 0 | -0.833138 | -3.874373 | -0.845564 |
| 57 | 1 | 0 | -1.061908 | -2.966089 | 0.656367 |
| 58 | 1 | 0 | -3.379121 | -3.597400 | 0.793010 |
| 59 | 1 | 0 | -2.726478 | -5.053813 | 0.051385 |
| 60 | 1 | 0 | -3.319169 | -4.271159 | -2.188582 |
| 61 | 1 | 0 | -4.724362 | -3.763844 | -1.261476 |
| 62 | 1 | 0 | -3.214454 | -1.974102 | -2.718978 |
| 63 | 6 | 0 | -4.317554 | -1.188135 | -1.129522 |
| 64 | 8 | 0 | -5.320720 | -0.984326 | -1.794141 |
| 65 | 7 | 0 | -4.095817 | -0.608124 | 0.071714 |
| 66 | 1 | 0 | -3.252950 | -0.803574 | 0.587807 |
| 67 | 6 | 0 | -4.965065 | 0.429874 | 0.592685 |
| 68 | 6 | 0 | -4.784133 | 1.781821 | -0.143253 |
| 69 | 1 | 0 | -5.099946 | 1.609067 | -1.172650 |
| 70 | 1 | 0 | -5.476024 | 2.506707 | 0.287702 |
| 71 | 6 | 0 | -3.346251 | 2.298675 | -0.117871 |
| 72 | 1 | 0 | -3.059642 | 2.607096 | 0.890175 |
| 73 | 1 | 0 | -2.682484 | 1.475140 | -0.375640 |
| 74 | 6 | 0 | -3.094314 | 3.462303 | -1.083328 |
| 75 | 1 | 0 | -3.591107 | 3.270752 | -2.040779 |
| 76 | 6 | 0 | -1.609678 | 3.724401 | -1.320136 |
| 77 | 1 | 0 | -1.046399 | 3.697499 | -0.389952 |
| 78 | 1 | 0 | -1.448202 | 4.682952 | -1.811252 |
| 79 | 7 | 0 | -0.991833 | 2.680861 | -2.200595 |
| 80 | 1 | 0 | -1.344966 | 2.750039 | -3.152335 |
| 81 | 1 | 0 | 0.064583 | 2.751126 | -2.253800 |
| 82 | 1 | 0 | -1.188390 | 1.709138 | -1.901665 |
| 83 | 1 | 0 | -3.540697 | 4.379509 | -0.693268 |
| 84 | 1 | 0 | -6.001878 | 0.117749 | 0.456005 |
| 85 | 6 | 0 | -4.711822 | 0.548718 | 2.083713 |
| 86 | 8 | 0 | -3.839110 | -0.020446 | 2.687091 |
| 87 | 8 | 0 | -5.586225 | 1.387944 | 2.670460 |
| 88 | 1 | 0 | -5.365222 | 1.429732 | 3.612351 |

31. **Table S31:** Cartesian Coordinates for the low-energy geometry of the **EA+QGPK** hexapeptide calculated using the B3LYP method and the 6-311+G(*2df,2pd*) level of theory.

| **Standard Orientation** | | | | | |
| --- | --- | --- | --- | --- | --- |
| Center Number | Atomic Number | Atomic Type | Coordinates (Angstroms) | | |
|  |  |  | X | Y | Z |
| 1 | 6 | 0 | 5.764620 | -0.582945 | -0.543221 |
| 2 | 7 | 0 | 5.804794 | -0.876775 | -1.971979 |
| 3 | 1 | 0 | 6.672601 | -1.328538 | -2.228128 |
| 4 | 1 | 0 | 5.723838 | -0.011750 | -2.493197 |
| 5 | 1 | 0 | 6.720557 | -0.168853 | -0.180931 |
| 6 | 6 | 0 | 4.817630 | 0.612627 | -0.331047 |
| 7 | 8 | 0 | 4.718369 | 1.461155 | -1.220648 |
| 8 | 7 | 0 | 4.232103 | 0.728836 | 0.876811 |
| 9 | 6 | 0 | 3.485774 | 1.900054 | 1.357050 |
| 10 | 1 | 0 | 3.308578 | 1.703494 | 2.412184 |
| 11 | 6 | 0 | 2.063417 | 2.001399 | 0.772886 |
| 12 | 8 | 0 | 1.097415 | 2.041593 | 1.528941 |
| 13 | 7 | 0 | 1.968499 | 2.004974 | -0.573150 |
| 14 | 6 | 0 | 0.697214 | 1.878211 | -1.269105 |
| 15 | 1 | 0 | 0.923991 | 1.845840 | -2.336826 |
| 16 | 6 | 0 | 0.066389 | 0.521635 | -0.876546 |
| 17 | 6 | 0 | 0.675870 | -0.669058 | -1.631857 |
| 18 | 6 | 0 | -0.095426 | -1.011314 | -2.892059 |
| 19 | 7 | 0 | -0.293336 | -2.331794 | -3.102164 |
| 20 | 1 | 0 | -0.299830 | -2.949381 | -2.298010 |
| 21 | 1 | 0 | -0.846540 | -2.591902 | -3.902588 |
| 22 | 8 | 0 | -0.499242 | -0.144665 | -3.668726 |
| 23 | 1 | 0 | 1.704038 | -0.448033 | -1.930479 |
| 24 | 1 | 0 | 0.718897 | -1.541803 | -0.987452 |
| 25 | 1 | 0 | -1.006057 | 0.509826 | -1.017980 |
| 26 | 1 | 0 | 0.224064 | 0.389073 | 0.187297 |
| 27 | 6 | 0 | -0.135903 | 3.173017 | -1.092585 |
| 28 | 8 | 0 | 0.337275 | 4.157405 | -0.561844 |
| 29 | 7 | 0 | -1.385284 | 3.262969 | -1.666309 |
| 30 | 1 | 0 | -1.812818 | 4.146379 | -1.433565 |
| 31 | 6 | 0 | -2.272834 | 2.242482 | -2.162545 |
| 32 | 1 | 0 | -2.959905 | 2.702129 | -2.879306 |
| 33 | 1 | 0 | -1.724613 | 1.485167 | -2.720102 |
| 34 | 6 | 0 | -3.120049 | 1.615837 | -1.044857 |
| 35 | 8 | 0 | -3.174589 | 2.154350 | 0.066923 |
| 36 | 7 | 0 | -3.801886 | 0.489001 | -1.314559 |
| 37 | 6 | 0 | -4.727608 | -0.081279 | -0.315052 |
| 38 | 6 | 0 | -5.561191 | -1.086527 | -1.138607 |
| 39 | 6 | 0 | -4.670660 | -1.448206 | -2.336215 |
| 40 | 6 | 0 | -3.928309 | -0.147999 | -2.642666 |
| 41 | 1 | 0 | -4.517671 | 0.491768 | -3.305806 |
| 42 | 1 | 0 | -2.953049 | -0.305193 | -3.098333 |
| 43 | 1 | 0 | -3.961847 | -2.233449 | -2.072326 |
| 44 | 1 | 0 | -5.243441 | -1.800461 | -3.191862 |
| 45 | 1 | 0 | -6.471579 | -0.594006 | -1.482224 |
| 46 | 1 | 0 | -5.857583 | -1.951932 | -0.550111 |
| 47 | 1 | 0 | -5.345400 | 0.713985 | 0.095632 |
| 48 | 6 | 0 | -4.045587 | -0.706428 | 0.927858 |
| 49 | 8 | 0 | -4.433329 | -0.345851 | 2.050041 |
| 50 | 7 | 0 | -3.105783 | -1.619266 | 0.699854 |
| 51 | 1 | 0 | -2.802499 | -1.828014 | -0.248358 |
| 52 | 6 | 0 | -2.324238 | -2.433939 | 1.643779 |
| 53 | 6 | 0 | -1.908159 | -1.724077 | 2.932468 |
| 54 | 1 | 0 | -2.807897 | -1.487572 | 3.498289 |
| 55 | 1 | 0 | -1.334635 | -2.446755 | 3.511191 |
| 56 | 6 | 0 | -1.073576 | -0.458956 | 2.705222 |
| 57 | 1 | 0 | -0.037538 | -0.734248 | 2.519856 |
| 58 | 1 | 0 | -1.408106 | 0.027227 | 1.787419 |
| 59 | 6 | 0 | -1.148656 | 0.555816 | 3.866002 |
| 60 | 1 | 0 | -1.833025 | 0.199239 | 4.641593 |
| 61 | 6 | 0 | -1.574681 | 1.946886 | 3.422573 |
| 62 | 1 | 0 | -0.875055 | 2.348174 | 2.693036 |
| 63 | 1 | 0 | -1.635021 | 2.630807 | 4.266873 |
| 64 | 7 | 0 | -2.948667 | 1.928583 | 2.767128 |
| 65 | 1 | 0 | -3.558832 | 2.636431 | 3.164428 |
| 66 | 1 | 0 | -2.911213 | 2.100764 | 1.741607 |
| 67 | 1 | 0 | -3.442902 | 1.018891 | 2.831711 |
| 68 | 1 | 0 | -0.176114 | 0.667098 | 4.345356 |
| 69 | 1 | 0 | -2.926689 | -3.305579 | 1.920410 |
| 70 | 6 | 0 | -1.111289 | -2.985637 | 0.821889 |
| 71 | 8 | 0 | -1.243680 | -2.981245 | -0.423407 |
| 72 | 8 | 0 | -0.130817 | -3.373214 | 1.494326 |
| 73 | 1 | 0 | 2.835541 | 1.995830 | -1.098825 |
| 74 | 6 | 0 | 4.264080 | 3.213374 | 1.242338 |
| 75 | 1 | 0 | 5.212665 | 3.128174 | 1.773005 |
| 76 | 1 | 0 | 3.682979 | 4.013380 | 1.700156 |
| 77 | 1 | 0 | 4.467584 | 3.479456 | 0.209399 |
| 78 | 1 | 0 | 4.047155 | -0.135116 | 1.380716 |
| 79 | 6 | 0 | 5.541419 | -1.874133 | 0.270843 |
| 80 | 6 | 0 | 4.343897 | -2.741674 | -0.130058 |
| 81 | 1 | 0 | 4.611595 | -3.798967 | -0.110406 |
| 82 | 1 | 0 | 4.066128 | -2.540361 | -1.166745 |
| 83 | 6 | 0 | 3.096610 | -2.601464 | 0.721784 |
| 84 | 8 | 0 | 2.962558 | -1.761145 | 1.598818 |
| 85 | 8 | 0 | 2.187675 | -3.490488 | 0.389072 |
| 86 | 1 | 0 | 1.292774 | -3.413847 | 0.880228 |
| 87 | 1 | 0 | 5.509841 | -1.646459 | 1.337014 |
| 88 | 1 | 0 | 6.448362 | -2.465965 | 0.129131 |

32. **Table S32:** Cartesian Coordinates for the low-energy geometry of the **AEEE+VA** hexapeptide calculated using the B3LYP method and the 6-311+G(*2df,2pd*) level of theory.

| **Standard Orientation** | | | | | |
| --- | --- | --- | --- | --- | --- |
| Center Number | Atomic Number | Atomic Type | Coordinates (Angstroms) | | |
|  |  |  | X | Y | Z |
| 1 | 6 | 0 | 3.925358 | -1.821712 | -0.081309 |
| 2 | 7 | 0 | 3.363588 | -0.571002 | 0.405847 |
| 3 | 1 | 0 | 3.045178 | -0.515263 | 1.365998 |
| 4 | 6 | 0 | 3.259188 | 0.564869 | -0.338370 |
| 5 | 6 | 0 | 2.736267 | 1.777873 | 0.474836 |
| 6 | 7 | 0 | 1.337941 | 1.553394 | 0.876831 |
| 7 | 6 | 0 | 1.021842 | 0.920826 | 2.023351 |
| 8 | 6 | 0 | -0.450747 | 0.934649 | 2.471358 |
| 9 | 7 | 0 | -1.410043 | 1.010591 | 1.377116 |
| 10 | 6 | 0 | -2.555690 | 1.748314 | 1.498661 |
| 11 | 6 | 0 | -3.730353 | 1.412657 | 0.555613 |
| 12 | 7 | 0 | -3.682409 | 0.054269 | 0.014740 |
| 13 | 6 | 0 | -4.335560 | -0.990597 | 0.586152 |
| 14 | 6 | 0 | -4.392381 | -2.295585 | -0.229236 |
| 15 | 7 | 0 | -4.328509 | -3.475880 | 0.632716 |
| 16 | 1 | 0 | -4.856683 | -3.271002 | 1.475785 |
| 17 | 1 | 0 | -3.366863 | -3.622852 | 0.929958 |
| 18 | 1 | 0 | -3.541471 | -2.340215 | -0.909733 |
| 19 | 6 | 0 | -5.692673 | -2.304763 | -1.044534 |
| 20 | 1 | 0 | -6.555215 | -2.305145 | -0.374908 |
| 21 | 1 | 0 | -5.726854 | -3.208666 | -1.652527 |
| 22 | 1 | 0 | -5.771135 | -1.433365 | -1.696278 |
| 23 | 8 | 0 | -4.872597 | -0.913466 | 1.687171 |
| 24 | 1 | 0 | -3.338885 | -0.069275 | -0.924669 |
| 25 | 6 | 0 | -3.936309 | 2.460708 | -0.554291 |
| 26 | 1 | 0 | -3.090538 | 2.433356 | -1.250145 |
| 27 | 6 | 0 | -5.256067 | 2.282124 | -1.309433 |
| 28 | 1 | 0 | -5.227397 | 1.418373 | -1.975843 |
| 29 | 1 | 0 | -6.055435 | 2.077774 | -0.587668 |
| 30 | 6 | 0 | -5.749054 | 3.511471 | -2.150276 |
| 31 | 8 | 0 | -6.595840 | 3.228092 | -3.033054 |
| 32 | 8 | 0 | -5.292239 | 4.632040 | -1.835458 |
| 33 | 1 | 0 | -3.924047 | 3.450730 | -0.101405 |
| 34 | 1 | 0 | -4.594946 | 1.445431 | 1.217496 |
| 35 | 8 | 0 | -2.683292 | 2.641257 | 2.323291 |
| 36 | 1 | 0 | -1.430422 | 0.190004 | 0.787479 |
| 37 | 6 | 0 | -0.753341 | -0.207010 | 3.477697 |
| 38 | 6 | 0 | -1.443758 | -1.474163 | 2.954255 |
| 39 | 6 | 0 | -0.830371 | -2.186819 | 1.746920 |
| 40 | 8 | 0 | -1.185112 | -3.402963 | 1.588250 |
| 41 | 8 | 0 | -0.087485 | -1.545112 | 0.976422 |
| 42 | 1 | 0 | -1.511040 | -2.199894 | 3.766851 |
| 43 | 1 | 0 | -2.480396 | -1.254460 | 2.686226 |
| 44 | 1 | 0 | 0.193539 | -0.462673 | 3.950358 |
| 45 | 1 | 0 | -1.399659 | 0.206437 | 4.255380 |
| 46 | 1 | 0 | -0.557041 | 1.878896 | 3.010116 |
| 47 | 8 | 0 | 1.887055 | 0.435261 | 2.757623 |
| 48 | 1 | 0 | 0.594064 | 1.915491 | 0.301942 |
| 49 | 6 | 0 | 2.957791 | 3.123849 | -0.215341 |
| 50 | 6 | 0 | 4.438302 | 3.519636 | -0.271620 |
| 51 | 6 | 0 | 4.746254 | 5.044508 | -0.467204 |
| 52 | 8 | 0 | 5.943511 | 5.296439 | -0.753398 |
| 53 | 8 | 0 | 3.803705 | 5.847542 | -0.287104 |
| 54 | 1 | 0 | 4.923101 | 3.248352 | 0.674645 |
| 55 | 1 | 0 | 4.962747 | 2.965531 | -1.048825 |
| 56 | 1 | 0 | 2.424230 | 3.896267 | 0.340272 |
| 57 | 1 | 0 | 2.536421 | 3.102609 | -1.222466 |
| 58 | 1 | 0 | 3.297430 | 1.780630 | 1.410276 |
| 59 | 8 | 0 | 3.566246 | 0.614256 | -1.517164 |
| 60 | 1 | 0 | 3.746594 | -2.538223 | 0.727204 |
| 61 | 6 | 0 | 3.157774 | -2.402300 | -1.302236 |
| 62 | 8 | 0 | 3.745314 | -2.960432 | -2.225876 |
| 63 | 7 | 0 | 1.818611 | -2.323966 | -1.224062 |
| 64 | 1 | 0 | 1.365962 | -1.928857 | -0.403117 |
| 65 | 6 | 0 | 0.957283 | -2.960718 | -2.214515 |
| 66 | 6 | 0 | 0.676974 | -2.056018 | -3.409610 |
| 67 | 1 | 0 | 0.145649 | -1.158665 | -3.097469 |
| 68 | 1 | 0 | 0.057067 | -2.574574 | -4.141998 |
| 69 | 1 | 0 | 1.617609 | -1.764659 | -3.874446 |
| 70 | 6 | 0 | -0.320832 | -3.421848 | -1.497238 |
| 71 | 8 | 0 | -0.067415 | -4.215735 | -0.477611 |
| 72 | 1 | 0 | -0.635797 | -3.941746 | 0.378561 |
| 73 | 8 | 0 | -1.427081 | -3.064629 | -1.852921 |
| 74 | 1 | 0 | 1.482496 | -3.856311 | -2.556538 |
| 75 | 6 | 0 | 5.457335 | -1.784574 | -0.335079 |
| 76 | 6 | 0 | 6.188615 | -0.870249 | 0.651319 |
| 77 | 1 | 0 | 7.266054 | -0.924749 | 0.476881 |
| 78 | 1 | 0 | 6.005360 | -1.171353 | 1.687221 |
| 79 | 1 | 0 | 5.890322 | 0.170357 | 0.548311 |
| 80 | 6 | 0 | 6.045356 | -3.200337 | -0.266881 |
| 81 | 1 | 0 | 5.937749 | -3.614550 | 0.741665 |
| 82 | 1 | 0 | 7.113275 | -3.181438 | -0.500760 |
| 83 | 1 | 0 | 5.551719 | -3.864475 | -0.972074 |
| 84 | 1 | 0 | 5.607628 | -1.398828 | -1.343357 |

33. **Table S33:** Cartesian Coordinates for the low-energy geometry of the **AE+EEVA** hexapeptide calculated using the B3LYP method and the 6-311+G(*2df,2pd*) level of theory.

| **Standard Orientation** | | | | | |
| --- | --- | --- | --- | --- | --- |
| **Center Number** | **Atomic Number** | Atomic Type | Coordinates (Angstroms) | | |
|  |  |  | X | Y | Z |
| 1 | 6 | 0 | -1.822220 | 2.245320 | -0.976400 |
| 2 | 6 | 0 | -2.005476 | 3.756150 | -0.731984 |
| 3 | 6 | 0 | -3.306959 | 4.366185 | -1.277988 |
| 4 | 6 | 0 | -4.528540 | 4.393071 | -0.308543 |
| 5 | 8 | 0 | -5.358745 | 5.304084 | -0.506246 |
| 6 | 8 | 0 | -4.572367 | 3.490514 | 0.574954 |
| 7 | 1 | 0 | -3.128851 | 5.397180 | -1.588096 |
| 8 | 1 | 0 | -3.625466 | 3.831429 | -2.179844 |
| 9 | 1 | 0 | -1.950185 | 3.942480 | 0.343406 |
| 10 | 1 | 0 | -1.143812 | 4.243978 | -1.186154 |
| 11 | 7 | 0 | -2.917607 | 1.505821 | -0.385604 |
| 12 | 1 | 0 | -3.586020 | 2.080919 | 0.158310 |
| 13 | 6 | 0 | -3.233825 | 0.253084 | -0.780848 |
| 14 | 6 | 0 | -4.472884 | -0.348423 | -0.085572 |
| 15 | 7 | 0 | -4.165993 | -1.654801 | 0.478486 |
| 16 | 6 | 0 | -3.155944 | -1.875898 | 1.318819 |
| 17 | 6 | 0 | -3.023590 | -3.323191 | 1.827059 |
| 18 | 7 | 0 | -1.662817 | -3.846395 | 1.646733 |
| 19 | 1 | 0 | -1.006197 | -3.159807 | 2.008571 |
| 20 | 1 | 0 | -1.457394 | -3.909439 | 0.655405 |
| 21 | 1 | 0 | -3.705910 | -3.958770 | 1.261630 |
| 22 | 6 | 0 | -3.402821 | -3.379893 | 3.307276 |
| 23 | 1 | 0 | -2.741382 | -2.734320 | 3.887712 |
| 24 | 1 | 0 | -3.305883 | -4.402343 | 3.675193 |
| 25 | 1 | 0 | -4.429561 | -3.043608 | 3.458860 |
| 26 | 8 | 0 | -2.347717 | -1.017027 | 1.699690 |
| 27 | 1 | 0 | -4.832866 | -2.413474 | 0.210036 |
| 28 | 6 | 0 | -5.627940 | -0.485516 | -1.091522 |
| 29 | 1 | 0 | -5.289491 | -1.121996 | -1.911229 |
| 30 | 6 | 0 | -6.963053 | -1.012091 | -0.527633 |
| 31 | 1 | 0 | -7.105914 | -0.636192 | 0.491619 |
| 32 | 1 | 0 | -7.786507 | -0.598994 | -1.110858 |
| 33 | 6 | 0 | -7.186653 | -2.553628 | -0.478961 |
| 34 | 8 | 0 | -8.366276 | -2.938374 | -0.606793 |
| 35 | 8 | 0 | -6.167704 | -3.283817 | -0.291926 |
| 36 | 1 | 0 | -5.796273 | 0.510793 | -1.506507 |
| 37 | 1 | 0 | -4.775784 | 0.332411 | 0.715051 |
| 38 | 8 | 0 | -2.598921 | -0.376169 | -1.621434 |
| 39 | 6 | 0 | -0.415992 | 1.842346 | -0.480301 |
| 40 | 7 | 0 | -0.337123 | 0.952614 | 0.521291 |
| 41 | 6 | 0 | 0.869737 | 0.572711 | 1.279711 |
| 42 | 6 | 0 | 1.510486 | 1.789633 | 1.980964 |
| 43 | 6 | 0 | 2.579665 | 1.446530 | 3.029526 |
| 44 | 6 | 0 | 4.061169 | 1.673278 | 2.626405 |
| 45 | 8 | 0 | 4.301443 | 1.969858 | 1.422520 |
| 46 | 8 | 0 | 4.901597 | 1.527855 | 3.542657 |
| 47 | 1 | 0 | 2.416335 | 2.033231 | 3.936107 |
| 48 | 1 | 0 | 2.504354 | 0.401958 | 3.343872 |
| 49 | 1 | 0 | 0.682391 | 2.320257 | 2.457215 |
| 50 | 1 | 0 | 1.942749 | 2.457631 | 1.239524 |
| 51 | 6 | 0 | 1.870658 | -0.296730 | 0.478640 |
| 52 | 8 | 0 | 2.180600 | -1.416630 | 0.870926 |
| 53 | 7 | 0 | 2.371391 | 0.263574 | -0.642471 |
| 54 | 6 | 0 | 3.208367 | -0.432254 | -1.605215 |
| 55 | 1 | 0 | 3.521360 | 0.348244 | -2.307306 |
| 56 | 6 | 0 | 4.536356 | -0.959993 | -1.006477 |
| 57 | 8 | 0 | 5.024453 | -2.028438 | -1.374223 |
| 58 | 7 | 0 | 5.204444 | -0.139570 | -0.168914 |
| 59 | 1 | 0 | 4.764582 | 0.673416 | 0.293037 |
| 60 | 6 | 0 | 6.470941 | -0.599469 | 0.365619 |
| 61 | 6 | 0 | 7.123853 | 0.462490 | 1.261666 |
| 62 | 1 | 0 | 7.317989 | 1.374045 | 0.695871 |
| 63 | 1 | 0 | 8.067274 | 0.072471 | 1.647708 |
| 64 | 1 | 0 | 6.468897 | 0.726723 | 2.092669 |
| 65 | 6 | 0 | 7.470610 | -0.982080 | -0.716030 |
| 66 | 8 | 0 | 7.419228 | -0.197106 | -1.820828 |
| 67 | 1 | 0 | 8.104576 | -0.539070 | -2.409708 |
| 68 | 8 | 0 | 8.329372 | -1.825331 | -0.591276 |
| 69 | 1 | 0 | 6.327884 | -1.517570 | 0.940135 |
| 70 | 6 | 0 | 2.477543 | -1.524611 | -2.437384 |
| 71 | 6 | 0 | 1.016950 | -1.161853 | -2.715993 |
| 72 | 1 | 0 | 0.560795 | -1.922556 | -3.353241 |
| 73 | 1 | 0 | 0.933674 | -0.204305 | -3.236743 |
| 74 | 1 | 0 | 0.413431 | -1.092972 | -1.815169 |
| 75 | 6 | 0 | 3.209760 | -1.767491 | -3.765114 |
| 76 | 1 | 0 | 3.176609 | -0.868005 | -4.389790 |
| 77 | 1 | 0 | 2.721639 | -2.569552 | -4.325024 |
| 78 | 1 | 0 | 4.248630 | -2.044115 | -3.602481 |
| 79 | 1 | 0 | 2.506770 | -2.446972 | -1.856505 |
| 80 | 1 | 0 | 1.979565 | 1.167041 | -0.901232 |
| 81 | 1 | 0 | 0.501784 | -0.108176 | 2.043903 |
| 82 | 1 | 0 | -1.195970 | 0.505153 | 0.828391 |
| 83 | 8 | 0 | 0.566557 | 2.373308 | -1.014991 |
| 84 | 1 | 0 | -1.797247 | 2.056294 | -2.053515 |

34. **Table S34:** Cartesian Coordinates for the low-energy geometry of the **EEEI+DG** hexapeptide calculated using the B3LYP method and the 6-311+G(*2df,2pd*) level of theory.

| **Standard Orientation** | | | | | |
| --- | --- | --- | --- | --- | --- |
| Center Number | Atomic Number | Atomic Type | Coordinates (Angstroms) | | |
|  |  |  | X | Y | Z |
| 1 | 6 | 0 | -4.725172 | -0.843764 | -1.883601 |
| 2 | 1 | 0 | -5.397045 | -1.004642 | -2.724440 |
| 3 | 7 | 0 | -4.419480 | 0.600763 | -1.987217 |
| 4 | 1 | 0 | -4.777841 | 1.022959 | -2.829487 |
| 5 | 6 | 0 | -3.673769 | 1.507724 | -1.292455 |
| 6 | 6 | 0 | -3.056892 | 1.120444 | 0.064852 |
| 7 | 7 | 0 | -1.586569 | 1.191550 | 0.002940 |
| 8 | 6 | 0 | -0.738587 | 2.135776 | 0.471341 |
| 9 | 6 | 0 | 0.754774 | 1.709649 | 0.530330 |
| 10 | 7 | 0 | 1.017801 | 1.590912 | 1.977262 |
| 11 | 6 | 0 | 1.641994 | 0.667451 | 2.742757 |
| 12 | 6 | 0 | 2.137384 | -0.661803 | 2.134389 |
| 13 | 7 | 0 | 3.149403 | -0.382208 | 1.102056 |
| 14 | 6 | 0 | 3.561081 | -1.058419 | 0.021411 |
| 15 | 6 | 0 | 4.883278 | -0.516558 | -0.580703 |
| 16 | 6 | 0 | 5.969977 | -1.600609 | -0.423371 |
| 17 | 6 | 0 | 7.246231 | -1.388170 | -1.267065 |
| 18 | 6 | 0 | 8.522789 | -0.933524 | -0.493994 |
| 19 | 8 | 0 | 9.617538 | -1.369063 | -0.936206 |
| 20 | 8 | 0 | 8.338851 | -0.158164 | 0.476071 |
| 21 | 1 | 0 | 7.496407 | -2.302139 | -1.808704 |
| 22 | 1 | 0 | 7.063098 | -0.622559 | -2.028444 |
| 23 | 1 | 0 | 6.247675 | -1.642511 | 0.633152 |
| 24 | 1 | 0 | 5.513537 | -2.555805 | -0.684058 |
| 25 | 7 | 0 | 5.287779 | 0.746543 | 0.022523 |
| 26 | 1 | 0 | 4.998696 | 1.548977 | -0.538960 |
| 27 | 1 | 0 | 6.290506 | 0.760357 | 0.192424 |
| 28 | 1 | 0 | 4.664373 | -0.404448 | -1.648986 |
| 29 | 8 | 0 | 3.012376 | -2.041045 | -0.500405 |
| 30 | 1 | 0 | 3.825005 | 0.362151 | 1.299250 |
| 31 | 6 | 0 | 0.981591 | -1.593568 | 1.727992 |
| 32 | 6 | 0 | 0.013002 | -1.914303 | 2.871540 |
| 33 | 6 | 0 | -1.190626 | -2.796461 | 2.422416 |
| 34 | 8 | 0 | -1.447136 | -3.800681 | 3.122219 |
| 35 | 8 | 0 | -1.808678 | -2.397457 | 1.395441 |
| 36 | 1 | 0 | 0.530892 | -2.420352 | 3.688163 |
| 37 | 1 | 0 | -0.404435 | -0.988895 | 3.279363 |
| 38 | 1 | 0 | 1.418811 | -2.519166 | 1.358300 |
| 39 | 1 | 0 | 0.419158 | -1.164243 | 0.907012 |
| 40 | 1 | 0 | 2.656667 | -1.127050 | 2.976073 |
| 41 | 8 | 0 | 1.805144 | 0.866744 | 3.948922 |
| 42 | 1 | 0 | 0.744192 | 2.412560 | 2.499276 |
| 43 | 1 | 0 | 0.898436 | 0.742353 | 0.061094 |
| 44 | 6 | 0 | 1.718078 | 2.738487 | -0.082921 |
| 45 | 1 | 0 | 1.492703 | 3.714933 | 0.354945 |
| 46 | 1 | 0 | 2.723266 | 2.466233 | 0.236802 |
| 47 | 6 | 0 | 1.731619 | 2.840722 | -1.609752 |
| 48 | 1 | 0 | 1.721110 | 1.833660 | -2.039489 |
| 49 | 1 | 0 | 0.853086 | 3.365623 | -1.984062 |
| 50 | 6 | 0 | 3.023472 | 3.544649 | -2.140934 |
| 51 | 8 | 0 | 2.863387 | 4.469637 | -2.972046 |
| 52 | 8 | 0 | 4.112501 | 3.096880 | -1.692576 |
| 53 | 8 | 0 | -1.054730 | 3.223334 | 0.944676 |
| 54 | 1 | 0 | -1.156632 | 0.365809 | -0.399453 |
| 55 | 6 | 0 | -3.690173 | 1.963951 | 1.205023 |
| 56 | 6 | 0 | -3.232676 | 1.429366 | 2.573957 |
| 57 | 6 | 0 | -3.473550 | 2.403789 | 3.729192 |
| 58 | 1 | 0 | -3.093025 | 1.992496 | 4.667294 |
| 59 | 1 | 0 | -4.534877 | 2.621233 | 3.871592 |
| 60 | 1 | 0 | -2.956286 | 3.348080 | 3.545053 |
| 61 | 1 | 0 | -2.169153 | 1.199794 | 2.539177 |
| 62 | 1 | 0 | -3.738745 | 0.480757 | 2.776257 |
| 63 | 6 | 0 | -5.221614 | 2.007051 | 1.108380 |
| 64 | 1 | 0 | -5.552528 | 2.435338 | 0.163050 |
| 65 | 1 | 0 | -5.685368 | 1.025026 | 1.207006 |
| 66 | 1 | 0 | -5.623603 | 2.638370 | 1.903734 |
| 67 | 1 | 0 | -3.311004 | 2.975911 | 1.085139 |
| 68 | 1 | 0 | -3.270914 | 0.073253 | 0.272282 |
| 69 | 8 | 0 | -3.541897 | 2.646424 | -1.727726 |
| 70 | 6 | 0 | -3.495730 | -1.700552 | -2.260632 |
| 71 | 8 | 0 | -3.240024 | -1.881763 | -3.451944 |
| 72 | 7 | 0 | -2.760822 | -2.253920 | -1.273220 |
| 73 | 1 | 0 | -2.929950 | -2.100416 | -0.283722 |
| 74 | 6 | 0 | -1.591560 | -3.047405 | -1.545814 |
| 75 | 1 | 0 | -1.537991 | -3.862698 | -0.827691 |
| 76 | 1 | 0 | -1.662323 | -3.451784 | -2.556080 |
| 77 | 6 | 0 | -0.290603 | -2.265451 | -1.441258 |
| 78 | 8 | 0 | 0.759680 | -3.075015 | -1.390992 |
| 79 | 1 | 0 | 1.584632 | -2.577842 | -1.122431 |
| 80 | 8 | 0 | -0.218062 | -1.056892 | -1.410847 |
| 81 | 6 | 0 | -5.557850 | -1.269423 | -0.673971 |
| 82 | 6 | 0 | -7.087962 | -0.927146 | -0.810508 |
| 83 | 8 | 0 | -7.714032 | -0.858971 | 0.271494 |
| 84 | 8 | 0 | -7.536509 | -0.809410 | -1.977173 |
| 85 | 1 | 0 | -5.188540 | -0.885376 | 0.273010 |
| 86 | 1 | 0 | -5.507086 | -2.359293 | -0.594726 |

35. **Table S35:** Cartesian Coordinates for the low-energy geometry of the **EE+EIDG** hexapeptide calculated using the B3LYP method and the 6-311+G(*2df,2pd*) level of theory.

| **Standard Orientation** | | | | | |
| --- | --- | --- | --- | --- | --- |
| Center Number | Atomic Number | Atomic Type | Coordinates (Angstroms) | | |
|  |  |  | X | Y | Z |
| 1 | 6 | 0 | 0.828394 | 0.685469 | 0.882502 |
| 2 | 7 | 0 | 2.089496 | 0.876074 | 0.177631 |
| 3 | 1 | 0 | 1.975100 | 1.090131 | -0.803237 |
| 4 | 6 | 0 | 3.320741 | 0.698983 | 0.715604 |
| 5 | 6 | 0 | 4.500644 | 0.960905 | -0.261236 |
| 6 | 7 | 0 | 5.396381 | -0.202611 | -0.287483 |
| 7 | 6 | 0 | 6.496610 | -0.487896 | 0.444865 |
| 8 | 6 | 0 | 7.129774 | -1.870942 | 0.120377 |
| 9 | 6 | 0 | 8.512405 | -1.678678 | -0.532156 |
| 10 | 6 | 0 | 9.531868 | -2.811408 | -0.319135 |
| 11 | 6 | 0 | 9.456880 | -4.109040 | -1.182846 |
| 12 | 8 | 0 | 10.452428 | -4.866686 | -1.076920 |
| 13 | 8 | 0 | 8.434260 | -4.288521 | -1.893328 |
| 14 | 1 | 0 | 10.538677 | -2.412808 | -0.474237 |
| 15 | 1 | 0 | 9.511157 | -3.130076 | 0.728951 |
| 16 | 1 | 0 | 8.363541 | -1.511470 | -1.602553 |
| 17 | 1 | 0 | 8.931924 | -0.762145 | -0.114427 |
| 18 | 7 | 0 | 6.276326 | -2.719928 | -0.717998 |
| 19 | 1 | 0 | 5.628147 | -3.252284 | -0.151465 |
| 20 | 1 | 0 | 6.869487 | -3.374877 | -1.237894 |
| 21 | 1 | 0 | 7.298110 | -2.305161 | 1.114664 |
| 22 | 8 | 0 | 7.019622 | 0.223324 | 1.292947 |
| 23 | 1 | 0 | 5.151870 | -0.962768 | -0.910624 |
| 24 | 6 | 0 | 5.222774 | 2.302973 | -0.015853 |
| 25 | 6 | 0 | 4.387614 | 3.543780 | -0.350988 |
| 26 | 6 | 0 | 5.222868 | 4.850751 | -0.588556 |
| 27 | 8 | 0 | 4.730669 | 5.908726 | -0.121264 |
| 28 | 8 | 0 | 6.274961 | 4.724757 | -1.261448 |
| 29 | 1 | 0 | 3.840260 | 3.373740 | -1.287274 |
| 30 | 1 | 0 | 3.639755 | 3.745571 | 0.415855 |
| 31 | 1 | 0 | 6.109392 | 2.318877 | -0.650017 |
| 32 | 1 | 0 | 5.572773 | 2.337285 | 1.014102 |
| 33 | 1 | 0 | 4.083165 | 1.010921 | -1.271149 |
| 34 | 8 | 0 | 3.498455 | 0.353235 | 1.875693 |
| 35 | 6 | 0 | 0.579557 | 1.763085 | 1.985820 |
| 36 | 1 | 0 | 1.453129 | 2.413865 | 1.977041 |
| 37 | 1 | 0 | -0.274337 | 2.383185 | 1.705420 |
| 38 | 6 | 0 | 0.383325 | 1.247730 | 3.419102 |
| 39 | 1 | 0 | 1.121660 | 0.467508 | 3.622426 |
| 40 | 1 | 0 | 0.616250 | 2.054214 | 4.117404 |
| 41 | 6 | 0 | -1.006152 | 0.700368 | 3.837029 |
| 42 | 8 | 0 | -1.805732 | 0.343447 | 2.917129 |
| 43 | 8 | 0 | -1.222333 | 0.642338 | 5.062942 |
| 44 | 6 | 0 | -0.275763 | 0.671670 | -0.215109 |
| 45 | 8 | 0 | 0.021980 | 0.887744 | -1.390927 |
| 46 | 7 | 0 | -1.511362 | 0.451946 | 0.265764 |
| 47 | 1 | 0 | -1.593873 | 0.376687 | 1.311422 |
| 48 | 6 | 0 | -2.787146 | 0.500917 | -0.452080 |
| 49 | 6 | 0 | -2.849548 | 1.523977 | -1.618269 |
| 50 | 6 | 0 | -2.550968 | 2.944004 | -1.109669 |
| 51 | 6 | 0 | -2.229114 | 3.946893 | -2.219605 |
| 52 | 1 | 0 | -1.988910 | 4.927271 | -1.801789 |
| 53 | 1 | 0 | -3.063226 | 4.080566 | -2.913163 |
| 54 | 1 | 0 | -1.362825 | 3.613483 | -2.793554 |
| 55 | 1 | 0 | -1.702613 | 2.909350 | -0.428495 |
| 56 | 1 | 0 | -3.404451 | 3.299995 | -0.521625 |
| 57 | 6 | 0 | -4.211425 | 1.468944 | -2.324169 |
| 58 | 1 | 0 | -4.399591 | 0.484264 | -2.752436 |
| 59 | 1 | 0 | -5.025145 | 1.705606 | -1.635128 |
| 60 | 1 | 0 | -4.244868 | 2.188684 | -3.144829 |
| 61 | 1 | 0 | -2.073694 | 1.240112 | -2.326094 |
| 62 | 1 | 0 | -3.513032 | 0.821077 | 0.295858 |
| 63 | 6 | 0 | -3.183011 | -0.902491 | -0.968174 |
| 64 | 8 | 0 | -2.554245 | -1.437961 | -1.871567 |
| 65 | 7 | 0 | -4.268567 | -1.532197 | -0.427933 |
| 66 | 6 | 0 | -5.201535 | -1.138031 | 0.627147 |
| 67 | 1 | 0 | -4.688854 | -0.483198 | 1.327203 |
| 68 | 6 | 0 | -6.370261 | -0.293722 | 0.069216 |
| 69 | 8 | 0 | -6.522803 | 0.887988 | 0.368778 |
| 70 | 7 | 0 | -7.180078 | -0.944766 | -0.798761 |
| 71 | 1 | 0 | -7.007028 | -1.970046 | -0.906277 |
| 72 | 6 | 0 | -8.196065 | -0.252856 | -1.516027 |
| 73 | 1 | 0 | -7.905937 | 0.791905 | -1.654812 |
| 74 | 1 | 0 | -8.324312 | -0.687336 | -2.511039 |
| 75 | 6 | 0 | -9.569527 | -0.228074 | -0.868571 |
| 76 | 8 | 0 | -10.457933 | 0.449992 | -1.670217 |
| 77 | 1 | 0 | -11.296651 | 0.436191 | -1.190198 |
| 78 | 8 | 0 | -9.918243 | -0.706219 | 0.173144 |
| 79 | 6 | 0 | -5.669024 | -2.379912 | 1.419092 |
| 80 | 6 | 0 | -6.293365 | -3.596977 | 0.666456 |
| 81 | 8 | 0 | -6.573450 | -4.573739 | 1.374337 |
| 82 | 8 | 0 | -6.459450 | -3.493624 | -0.593516 |
| 83 | 1 | 0 | -6.395096 | -2.061535 | 2.170230 |
| 84 | 1 | 0 | -4.810479 | -2.758909 | 1.974729 |
| 85 | 1 | 0 | -4.511149 | -2.401934 | -0.888244 |
| 86 | 1 | 0 | 0.837510 | -0.296508 | 1.361863 |

36. **Table S36:** Cartesian Coordinates for the low-energy geometry of the **PEDK+GH** hexapeptide calculated using the B3LYP method and the 6-311+G(*2df,2pd*) level of theory.

| **Standard Orientation** | | | | | |
| --- | --- | --- | --- | --- | --- |
| Center Number | Atomic Number | Atomic Type | Coordinates (Angstroms) | | |
|  |  |  | X | Y | Z |
| 1 | 6 | 0 | 4.910310 | -5.205473 | -0.390965 |
| 2 | 7 | 0 | 4.253992 | -4.017952 | -0.936771 |
| 3 | 1 | 0 | 3.929185 | -4.138740 | -1.886978 |
| 4 | 6 | 0 | 5.268730 | -2.956404 | -0.859409 |
| 5 | 1 | 0 | 6.070918 | -3.137273 | -1.596177 |
| 6 | 6 | 0 | 5.853232 | -3.152755 | 0.550402 |
| 7 | 6 | 0 | 5.638459 | -4.660069 | 0.851928 |
| 8 | 1 | 0 | 6.574986 | -5.187940 | 1.027891 |
| 9 | 1 | 0 | 5.023412 | -4.788913 | 1.742375 |
| 10 | 1 | 0 | 6.897797 | -2.857922 | 0.599558 |
| 11 | 1 | 0 | 5.315269 | -2.530544 | 1.262698 |
| 12 | 6 | 0 | 4.688534 | -1.588550 | -1.209984 |
| 13 | 8 | 0 | 3.713897 | -1.571172 | -2.042360 |
| 14 | 7 | 0 | 5.259229 | -0.540934 | -0.688836 |
| 15 | 6 | 0 | 4.775081 | 0.754864 | -1.117500 |
| 16 | 6 | 0 | 5.786115 | 1.851600 | -0.706715 |
| 17 | 6 | 0 | 6.123236 | 1.930912 | 0.797422 |
| 18 | 6 | 0 | 7.303849 | 1.082189 | 1.269756 |
| 19 | 8 | 0 | 7.334392 | -0.175301 | 0.854491 |
| 20 | 1 | 0 | 6.520279 | -0.398966 | 0.261664 |
| 21 | 8 | 0 | 8.157238 | 1.538961 | 1.999421 |
| 22 | 1 | 0 | 5.251014 | 1.638648 | 1.388171 |
| 23 | 1 | 0 | 6.365287 | 2.955233 | 1.071470 |
| 24 | 1 | 0 | 5.392939 | 2.819241 | -1.024307 |
| 25 | 1 | 0 | 6.699483 | 1.688827 | -1.282379 |
| 26 | 1 | 0 | 4.680500 | 0.778526 | -2.210270 |
| 27 | 6 | 0 | 3.384743 | 1.108662 | -0.550632 |
| 28 | 8 | 0 | 2.962975 | 0.694350 | 0.521446 |
| 29 | 7 | 0 | 2.670554 | 1.979987 | -1.313236 |
| 30 | 6 | 0 | 1.482385 | 2.700975 | -0.857270 |
| 31 | 6 | 0 | 0.367765 | 1.721297 | -0.417189 |
| 32 | 7 | 0 | -0.284338 | 2.037138 | 0.662787 |
| 33 | 6 | 0 | -1.303046 | 1.176860 | 1.256684 |
| 34 | 6 | 0 | -0.760740 | -0.217063 | 1.721836 |
| 35 | 1 | 0 | -1.161568 | -0.417888 | 2.717900 |
| 36 | 1 | 0 | 0.310769 | -0.069132 | 1.852260 |
| 37 | 6 | 0 | -1.034647 | -1.442473 | 0.832773 |
| 38 | 1 | 0 | -2.004107 | -1.875112 | 1.091492 |
| 39 | 6 | 0 | 0.033407 | -2.540310 | 0.936509 |
| 40 | 1 | 0 | 0.181025 | -2.831302 | 1.980484 |
| 41 | 6 | 0 | 1.402468 | -2.169716 | 0.370095 |
| 42 | 1 | 0 | 1.869757 | -1.350544 | 0.911468 |
| 43 | 7 | 0 | 1.339594 | -1.723968 | -1.053548 |
| 44 | 1 | 0 | 0.806651 | -2.378094 | -1.619683 |
| 45 | 1 | 0 | 2.344304 | -1.667800 | -1.474608 |
| 46 | 1 | 0 | 0.905149 | -0.777432 | -1.139812 |
| 47 | 1 | 0 | 2.082605 | -3.019781 | 0.403166 |
| 48 | 1 | 0 | -0.331845 | -3.436981 | 0.424053 |
| 49 | 1 | 0 | -1.122913 | -1.140216 | -0.207162 |
| 50 | 6 | 0 | -2.618790 | 1.072269 | 0.480456 |
| 51 | 8 | 0 | -3.711735 | 1.090259 | 1.071551 |
| 52 | 7 | 0 | -2.546087 | 0.977415 | -0.855252 |
| 53 | 6 | 0 | -3.730914 | 0.990266 | -1.673185 |
| 54 | 1 | 0 | -3.428800 | 0.857832 | -2.712120 |
| 55 | 1 | 0 | -4.243543 | 1.952761 | -1.594165 |
| 56 | 6 | 0 | -4.698858 | -0.141670 | -1.331775 |
| 57 | 8 | 0 | -4.367892 | -1.303203 | -1.204562 |
| 58 | 7 | 0 | -6.014963 | 0.236846 | -1.204123 |
| 59 | 1 | 0 | -6.282595 | 1.207356 | -1.239035 |
| 60 | 6 | 0 | -7.000236 | -0.723700 | -0.809720 |
| 61 | 1 | 0 | -6.790525 | -1.664103 | -1.331149 |
| 62 | 6 | 0 | -6.935417 | -1.006973 | 0.675889 |
| 63 | 6 | 0 | -6.045130 | -0.543727 | 1.595466 |
| 64 | 7 | 0 | -6.365192 | -1.162023 | 2.785262 |
| 65 | 6 | 0 | -7.405705 | -1.975305 | 2.622525 |
| 66 | 7 | 0 | -7.763895 | -1.894556 | 1.342647 |
| 67 | 1 | 0 | -8.551790 | -2.366343 | 0.921977 |
| 68 | 1 | 0 | -7.875844 | -2.575973 | 3.379350 |
| 69 | 1 | 0 | -5.869156 | -1.023665 | 3.653186 |
| 70 | 1 | 0 | -5.194028 | 0.127446 | 1.480468 |
| 71 | 6 | 0 | -8.392981 | -0.260152 | -1.234319 |
| 72 | 8 | 0 | -8.650587 | 0.787926 | -1.743149 |
| 73 | 8 | 0 | -9.317923 | -1.211519 | -0.948283 |
| 74 | 1 | 0 | -10.183646 | -0.889458 | -1.243766 |
| 75 | 1 | 0 | -1.605038 | 0.867024 | -1.261442 |
| 76 | 1 | 0 | -1.610777 | 1.694020 | 2.165346 |
| 77 | 8 | 0 | 0.132406 | 0.738460 | -1.204627 |
| 78 | 1 | 0 | 1.089024 | 3.193167 | -1.750686 |
| 79 | 6 | 0 | 1.884369 | 3.788025 | 0.168658 |
| 80 | 6 | 0 | 0.904703 | 4.948337 | 0.344964 |
| 81 | 8 | 0 | 1.197647 | 6.082818 | 0.043411 |
| 82 | 8 | 0 | -0.276069 | 4.636557 | 0.867475 |
| 83 | 1 | 0 | -0.328995 | 3.617637 | 0.932426 |
| 84 | 1 | 0 | 2.817057 | 4.234692 | -0.165343 |
| 85 | 1 | 0 | 2.057218 | 3.308991 | 1.132590 |
| 86 | 1 | 0 | 3.054180 | 2.241115 | -2.204680 |
| 87 | 1 | 0 | 5.642416 | -5.635933 | -1.091778 |
| 88 | 1 | 0 | 4.181360 | -5.982726 | -0.155691 |

37. **Table S37:** Cartesian Coordinates for the low-energy geometry of the **PE+DKGH** hexapeptide calculated using the B3LYP method and the 6-311+G(*2df,2pd*) level of theory.

| **Standard Orientation** | | | | | |
| --- | --- | --- | --- | --- | --- |
| Center Number | Atomic Number | Atomic Type | Coordinates (Angstroms) | | |
|  |  |  | X | Y | Z |
| 1 | 6 | 0 | -0.228737 | -3.244215 | -1.304250 |
| 2 | 7 | 0 | -0.546330 | -1.856825 | -0.938784 |
| 3 | 6 | 0 | -1.615423 | -1.217251 | -1.389880 |
| 4 | 6 | 0 | -1.850064 | 0.240550 | -0.919998 |
| 5 | 7 | 0 | -3.208925 | 0.223911 | -0.366754 |
| 6 | 1 | 0 | -3.848999 | -0.208726 | -1.018878 |
| 7 | 6 | 0 | -3.578390 | 0.016980 | 0.928461 |
| 8 | 6 | 0 | -2.997434 | 0.874530 | 2.063778 |
| 9 | 7 | 0 | -2.334217 | 2.079113 | 1.611390 |
| 10 | 6 | 0 | -3.067873 | 3.121372 | 1.151808 |
| 11 | 6 | 0 | -2.341402 | 4.403706 | 0.760280 |
| 12 | 7 | 0 | -2.638475 | 4.693403 | -0.646606 |
| 13 | 6 | 0 | -1.786315 | 5.829888 | -0.994217 |
| 14 | 6 | 0 | -0.442378 | 5.479614 | -0.327246 |
| 15 | 6 | 0 | -0.806723 | 4.482502 | 0.806689 |
| 16 | 1 | 0 | -0.454299 | 4.809808 | 1.783293 |
| 17 | 1 | 0 | -0.349273 | 3.516371 | 0.606083 |
| 18 | 1 | 0 | 0.056578 | 6.371013 | 0.050455 |
| 19 | 1 | 0 | 0.223811 | 5.002496 | -1.042480 |
| 20 | 1 | 0 | -2.173248 | 6.776868 | -0.588482 |
| 21 | 1 | 0 | -1.708125 | 5.935886 | -2.076039 |
| 22 | 1 | 0 | -3.625848 | 4.862606 | -0.787905 |
| 23 | 1 | 0 | -2.757041 | 5.178624 | 1.426029 |
| 24 | 8 | 0 | -4.286530 | 3.057424 | 1.033222 |
| 25 | 1 | 0 | -1.332014 | 2.139769 | 1.780672 |
| 26 | 6 | 0 | -2.153344 | 0.090676 | 3.090779 |
| 27 | 6 | 0 | -0.951173 | -0.703725 | 2.578792 |
| 28 | 6 | 0 | 0.191080 | 0.079321 | 1.902620 |
| 29 | 8 | 0 | 0.307356 | 1.292000 | 2.114652 |
| 30 | 8 | 0 | 0.942386 | -0.619049 | 1.139481 |
| 31 | 1 | 0 | -1.273803 | -1.493358 | 1.899440 |
| 32 | 1 | 0 | -0.497304 | -1.225359 | 3.426443 |
| 33 | 1 | 0 | -2.830397 | -0.601058 | 3.593629 |
| 34 | 1 | 0 | -1.825063 | 0.811873 | 3.839329 |
| 35 | 1 | 0 | -3.913961 | 1.179957 | 2.574881 |
| 36 | 8 | 0 | -4.482514 | -0.790068 | 1.190572 |
| 37 | 1 | 0 | -1.149642 | 0.508987 | -0.137499 |
| 38 | 6 | 0 | -1.736579 | 1.252635 | -2.076296 |
| 39 | 6 | 0 | -0.306263 | 1.294780 | -2.583580 |
| 40 | 8 | 0 | 0.260175 | 2.487254 | -2.468168 |
| 41 | 1 | 0 | 1.255001 | 2.421301 | -2.594088 |
| 42 | 8 | 0 | 0.254937 | 0.305554 | -3.004843 |
| 43 | 1 | 0 | -2.058845 | 2.232141 | -1.728250 |
| 44 | 1 | 0 | -2.383411 | 0.931673 | -2.894672 |
| 45 | 8 | 0 | -2.504074 | -1.757416 | -2.082186 |
| 46 | 1 | 0 | 0.087141 | -1.331632 | -0.328085 |
| 47 | 6 | 0 | -1.353425 | -4.256317 | -0.947917 |
| 48 | 1 | 0 | -0.918382 | -5.251808 | -1.035757 |
| 49 | 1 | 0 | -2.109230 | -4.177886 | -1.725947 |
| 50 | 6 | 0 | -1.978711 | -4.051136 | 0.434509 |
| 51 | 1 | 0 | -1.304558 | -4.419816 | 1.209322 |
| 52 | 6 | 0 | -3.350386 | -4.703555 | 0.653878 |
| 53 | 1 | 0 | -3.265087 | -5.792532 | 0.631388 |
| 54 | 6 | 0 | -4.441332 | -4.346134 | -0.355624 |
| 55 | 1 | 0 | -4.271096 | -4.831429 | -1.314532 |
| 56 | 7 | 0 | -4.542401 | -2.873788 | -0.649806 |
| 57 | 1 | 0 | -4.583436 | -2.233663 | 0.184641 |
| 58 | 1 | 0 | -5.383615 | -2.688473 | -1.192048 |
| 59 | 1 | 0 | -3.732715 | -2.534446 | -1.231924 |
| 60 | 1 | 0 | -5.412441 | -4.669406 | 0.014693 |
| 61 | 1 | 0 | -3.698360 | -4.445951 | 1.656522 |
| 62 | 1 | 0 | -2.079620 | -2.983988 | 0.631859 |
| 63 | 6 | 0 | 1.128401 | -3.755814 | -0.775317 |
| 64 | 8 | 0 | 1.599611 | -4.737491 | -1.328953 |
| 65 | 7 | 0 | 1.705834 | -3.124019 | 0.270363 |
| 66 | 6 | 0 | 3.043268 | -3.465206 | 0.720956 |
| 67 | 1 | 0 | 3.356366 | -4.377411 | 0.223035 |
| 68 | 1 | 0 | 3.038270 | -3.653326 | 1.797499 |
| 69 | 6 | 0 | 4.132373 | -2.417486 | 0.459997 |
| 70 | 8 | 0 | 5.297163 | -2.765992 | 0.364443 |
| 71 | 7 | 0 | 3.733006 | -1.119999 | 0.366123 |
| 72 | 1 | 0 | 2.791150 | -0.839128 | 0.640155 |
| 73 | 6 | 0 | 4.708774 | -0.075246 | 0.206901 |
| 74 | 1 | 0 | 5.596219 | -0.532080 | -0.243373 |
| 75 | 6 | 0 | 4.162228 | 1.003702 | -0.690189 |
| 76 | 6 | 0 | 3.098946 | 0.972746 | -1.546970 |
| 77 | 7 | 0 | 2.946170 | 2.195937 | -2.159080 |
| 78 | 6 | 0 | 3.909161 | 2.957113 | -1.691832 |
| 79 | 7 | 0 | 4.676709 | 2.277759 | -0.803659 |
| 80 | 1 | 0 | 5.463836 | 2.629787 | -0.284645 |
| 81 | 1 | 0 | 4.087657 | 3.986843 | -1.949898 |
| 82 | 1 | 0 | 2.429973 | 0.160278 | -1.759358 |
| 83 | 6 | 0 | 5.190393 | 0.445210 | 1.571547 |
| 84 | 8 | 0 | 4.819477 | 0.067922 | 2.642006 |
| 85 | 8 | 0 | 6.159908 | 1.393016 | 1.421360 |
| 86 | 1 | 0 | 6.447541 | 1.647405 | 2.310575 |
| 87 | 1 | 0 | 1.259452 | -2.324316 | 0.719709 |
| 88 | 1 | 0 | -0.107990 | -3.277037 | -2.386901 |

38.
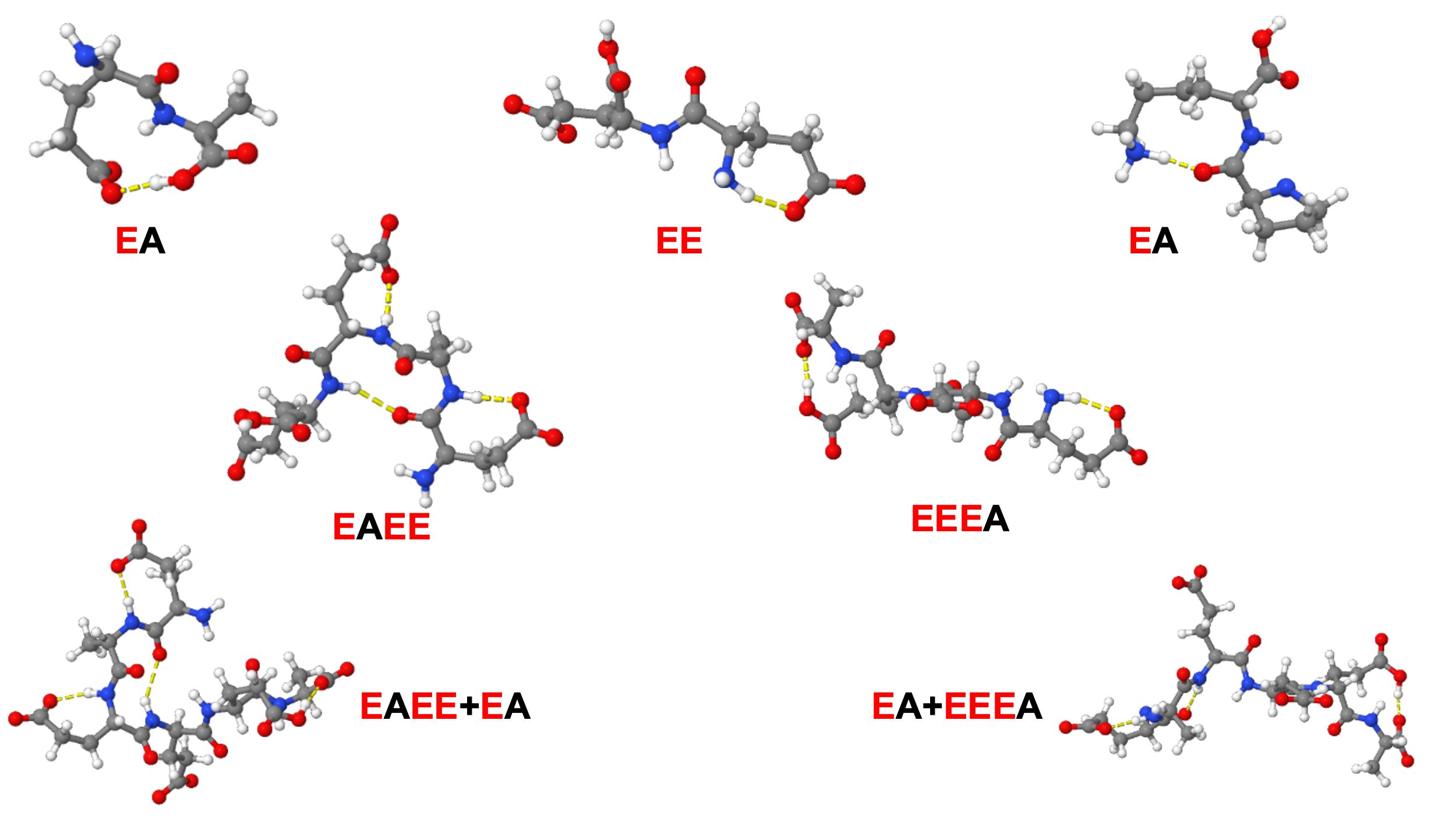

**Figure S1**: All structures optimized at the 6-311+G(2df,pd) basis set used in the buildup of the βI E-hook hexamers.

39.
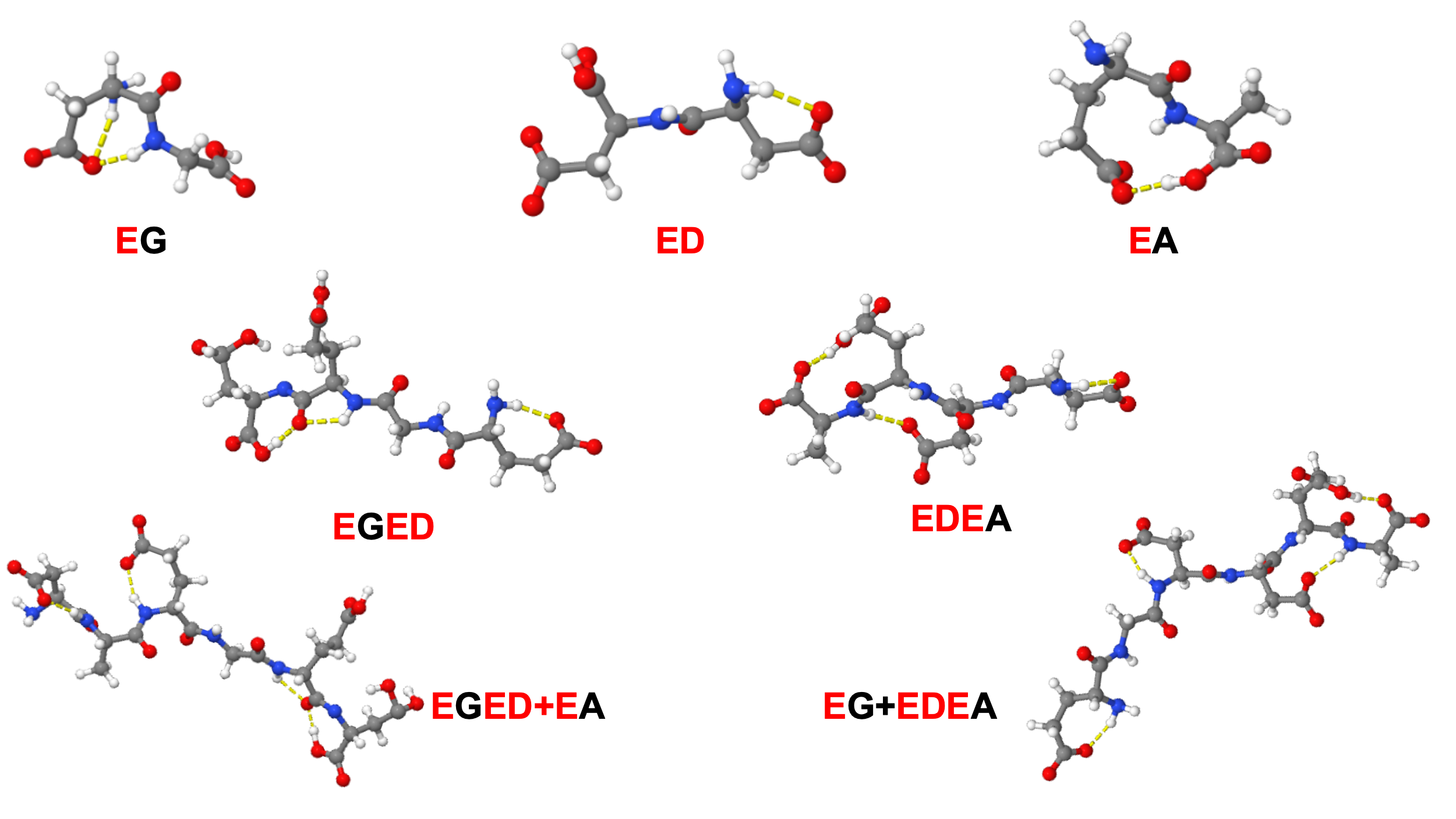

**Figure S2:** All structures optimized at the 6-311+G(2df,pd) basis set used in the buildup of the βII E-hook hexamers.

40.
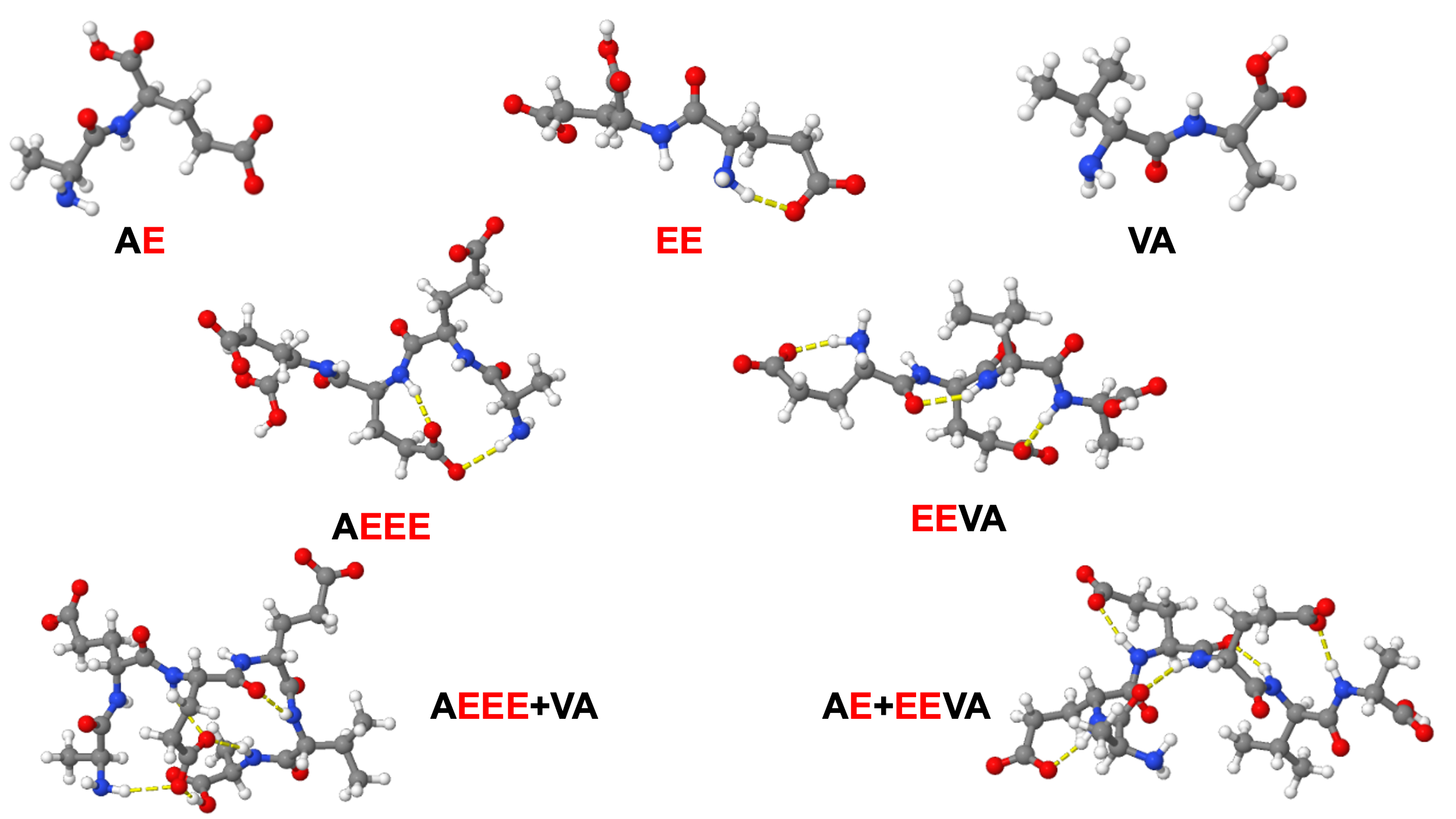

**Figure S3:** All structures optimized at the 6-311+G(2df,pd) basis set used in the buildup of the βIV E-hook hexamers.

41.
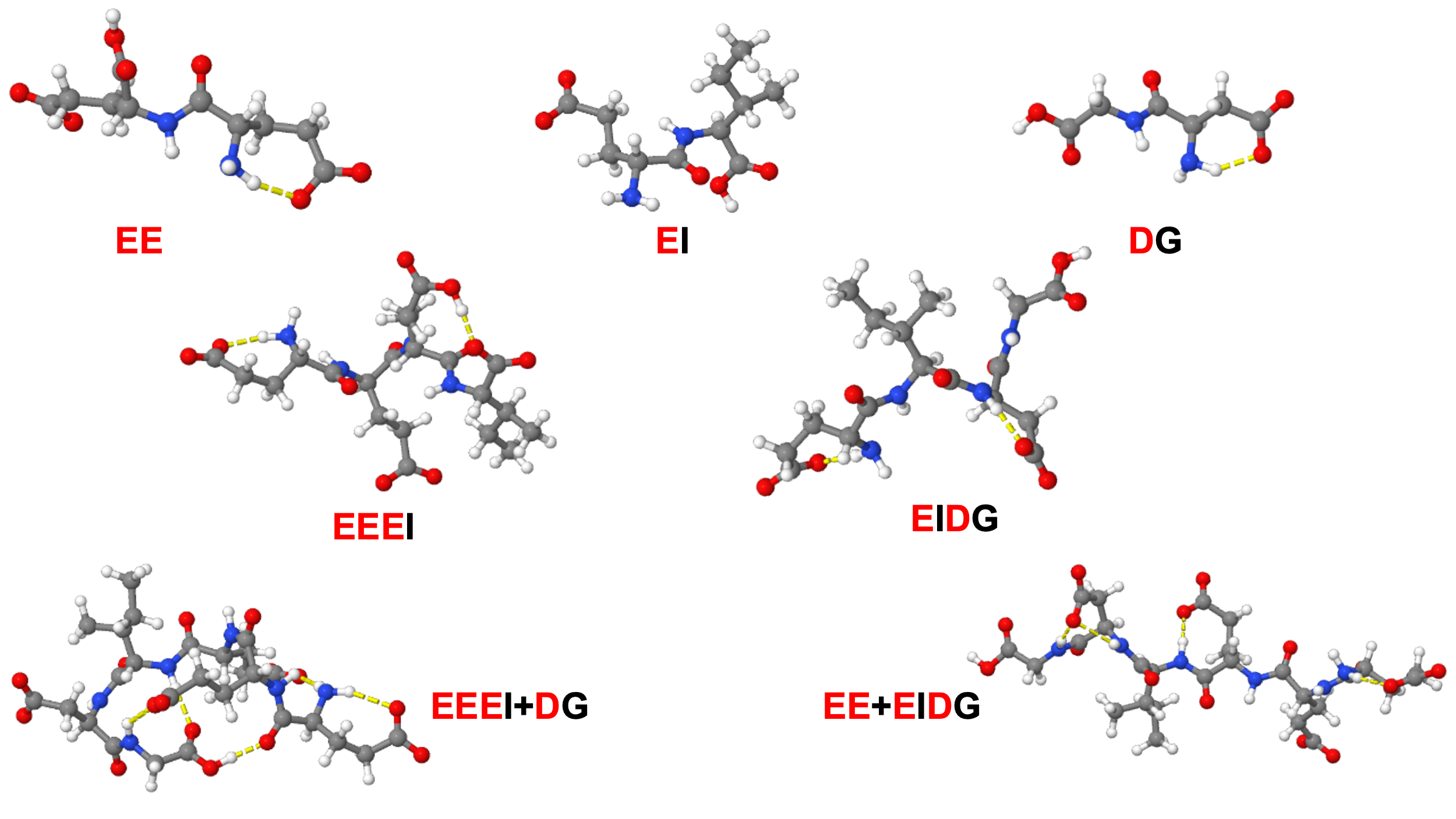

**Figure S4:** All structures optimized at the 6-311+G(2df,pd) basis set used in the buildup of the βV E-hook hexamers.

42.
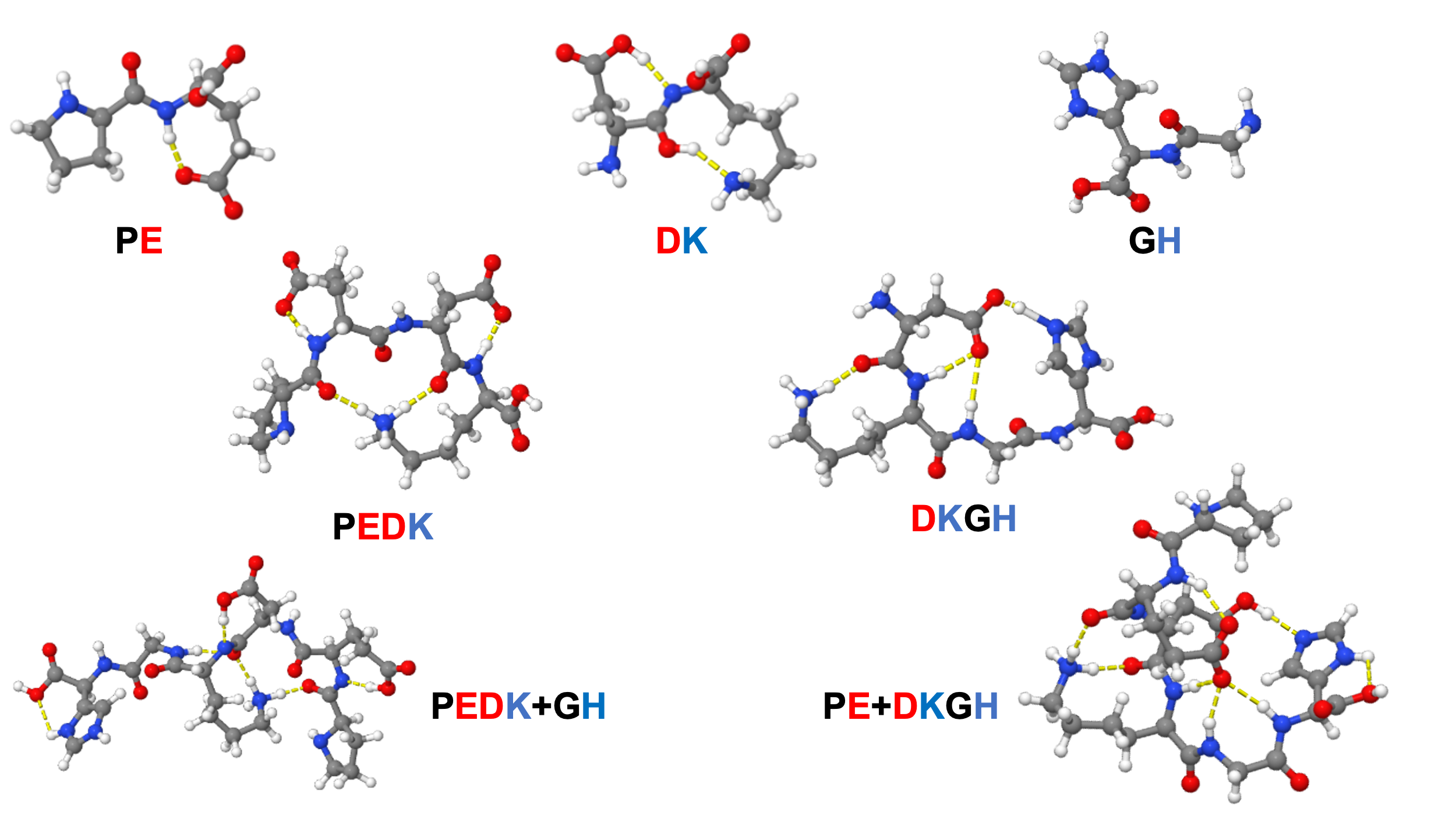

**Figure S5:** All structures optimized at the 6-311+G(2df,pd) basis set used in the buildup of the βVI E-hook hexamers.

43. Python code used in the dynamic time warping analysis of the experimental and computed Raman spectra:

import os

import numpy as np

import matplotlib.pyplot as plt

from scipy.ndimage import gaussian_filter1d

from scipy.signal import find_peaks

from scipy.stats import pearsonr

def calculate_peak_metrics(results):

    matched_exp = results["matched_exp_peak_indices"]

    exp_peaks = results["exp_peaks"]

    assignments = results["dtw_peak_assignments"]

    match_fraction = len(matched_exp) / len(exp_peaks) if len(exp_peaks) > 0 else np.nan

    shifts = np.array([item["shift"] for item in assignments])

    peak_rms = np.sqrt(np.mean(shifts**2)) if len(shifts) > 0 else np.nan

    return match_fraction, peak_rms

###############################################################################

\# GRAPH / ANALYSIS OPTIONS

###############################################################################

GRAPH = {

    "figsize": (32, 16),

    "dpi": 300,

    "save_dpi": 1200,

    "spectrum_linewidth": 2.5,

    "match_linewidth": 1.5,

    "warped_match_linewidth": 1.0,

    "marker_size": 100,

    "axis_fontsize": 32,

    "tick_fontsize": 26,

    "legend_fontsize": 24,

    "matched_color": "black",

    "unmatched_color": "red",

    "match_line_color": "black",

    "match_line_alpha": 0.7,

    "overall_original_title": "PEDKGH Experiment vs PE+DKGH Simulation",

    "overall_warped_title": "DTW-Warped Spectra: PEDKGH vs PE+DKGH",

}

FINGERPRINT_REGION = (200, 2000)

EXP_SMOOTH_SIGMA = 4

SIM_SMOOTH_SIGMA = 2

DTW_WINDOW = 50

EXP_PEAK_SNAP_TOLERANCE = 40

EXP_PEAK_HEIGHT = 0.002

EXP_PEAK_PROMINENCE = 0.002

EXP_PEAK_DISTANCE = 10

SIM_PEAK_HEIGHT = 0.002

SIM_PEAK_PROMINENCE = 0.002

SIM_PEAK_DISTANCE = 5

WAVENUMBER_WEIGHT = 2.0

INTENSITY_WEIGHT = 100.0

COVERAGE_BONUS = 10.0

OCCUPANCY_PENALTY_WEIGHT = 15.0

SAVE_ORIGINAL_FIGURE = True

SAVE_WARPED_FIGURE = True

ORIGINAL_FIGURE_NAME = "Originals.svg"

WARPED_FIGURE_NAME = "Warped.svg"

PRINT_FIRST_PASS_ATTEMPTS = True

PRINT_SECOND_PASS_ATTEMPTS = True

NEWFILENAME = input("Select new figure name: ")

ORIGINAL_FIGURE_NAME = NEWFILENAME + ORIGINAL_FIGURE_NAME

WARPED_FIGURE_NAME = NEWFILENAME + WARPED_FIGURE_NAME

###############################################################################

\# LIST TXT FILES AND SELECT INPUTS

###############################################################################

script_dir = os.path.dirname(os.path.abspath(__file__))

txt_files = [

    file for file in os.listdir(script_dir)

    if file.lower().endswith(".txt")

]

if len(txt_files) < 2:

    raise RuntimeError("Need at least two .txt files in this directory.")

print("\nAvailable .txt files:\n")

for i, file in enumerate(txt_files):

    print(f"{i}: {file}")

exp_choice = int(input("\nEnter number for experimental spectrum: "))

sim_choice = int(input("Enter number for simulated spectrum: "))

exp_file = os.path.join(script_dir, txt_files[exp_choice])

sim_file = os.path.join(script_dir, txt_files[sim_choice])

print(f"\nExperimental file: {txt_files[exp_choice]}")

print(f"Simulated file:    {txt_files[sim_choice]}")

###############################################################################

\# LOAD DATA

###############################################################################

exp_data = np.loadtxt(exp_file)

sim_data = np.loadtxt(sim_file)

exp_x_raw = exp_data[:, 0]

exp_y_raw = exp_data[:, 1]

sim_x_raw = sim_data[:, 0]

sim_y_raw = sim_data[:, 1]

###############################################################################

\# REGION PREPROCESSING FUNCTION

###############################################################################

def prepare_region(exp_x_raw, exp_y_raw, sim_x_raw, sim_y_raw, xmin, xmax):

    """

    Restrict, interpolate, vector normalize, and smooth one spectral region.

    The experimental and simulated spectra are normalized separately within the

    requested region.

    """

    exp_mask = (exp_x_raw >= xmin) & (exp_x_raw <= xmax)

    sim_mask = (sim_x_raw >= xmin) & (sim_x_raw <= xmax)

    exp_x_region = exp_x_raw[exp_mask]

    exp_y_region = exp_y_raw[exp_mask]

    sim_x_region = sim_x_raw[sim_mask]

    sim_y_region = sim_y_raw[sim_mask]

    if len(exp_x_region) == 0:

        raise ValueError(f"No experimental data found in {xmin}-{xmax} cm^-1.")

    if len(sim_x_region) == 0:

        raise ValueError(f"No simulated data found in {xmin}-{xmax} cm^-1.")

    sim_y_interp_region = np.interp(

        exp_x_region,

        sim_x_region,

        sim_y_region

    )

    # Remove offsets before vector normalization.

    exp_y_region = exp_y_region - np.min(exp_y_region)

    sim_y_interp_region = sim_y_interp_region - np.min(sim_y_interp_region)

    exp_norm = np.linalg.norm(exp_y_region)

    sim_norm = np.linalg.norm(sim_y_interp_region)

    if exp_norm == 0:

        raise ValueError(f"Experimental spectrum norm is zero in {xmin}-{xmax} cm^-1.")

    if sim_norm == 0:

        raise ValueError(f"Simulated spectrum norm is zero in {xmin}-{xmax} cm^-1.")

    exp_y_region = exp_y_region / exp_norm

    sim_y_interp_region = sim_y_interp_region / sim_norm

    exp_y_region = gaussian_filter1d(exp_y_region, sigma=EXP_SMOOTH_SIGMA)

    sim_y_interp_region = gaussian_filter1d(sim_y_interp_region, sigma=SIM_SMOOTH_SIGMA)

    return exp_x_region, exp_y_region, sim_y_interp_region

###############################################################################

\# DTW FUNCTIONS

###############################################################################

def dtw_distance(y1, y2, window=None):

    n = len(y1)

    m = len(y2)

    if window is None:

        window = max(n, m)

    else:

        window = max(window, abs(n - m))

    dtw = np.full((n + 1, m + 1), np.inf)

    dtw[0, 0] = 0

    for i in range(1, n + 1):

        j_start = max(1, i - window)

        j_end = min(m, i + window)

        for j in range(j_start, j_end + 1):

            cost = abs(y1[i - 1] - y2[j - 1])

            dtw[i, j] = cost + min(

                dtw[i - 1, j],

                dtw[i, j - 1],

                dtw[i - 1, j - 1]

            )

    return dtw[n, m], dtw

def get_dtw_path(dtw_matrix):

    i = dtw_matrix.shape[0] - 1

    j = dtw_matrix.shape[1] - 1

    path = []

    while i > 0 and j > 0:

        path.append((i - 1, j - 1))

        steps = [

            dtw_matrix[i - 1, j],

            dtw_matrix[i, j - 1],

            dtw_matrix[i - 1, j - 1]

        ]

        best_step = np.argmin(steps)

        if best_step == 0:

            i -= 1

        elif best_step == 1:

            j -= 1

        else:

            i -= 1

            j -= 1

    path.reverse()

    return np.array(path)

###############################################################################

\# REGION ANALYSIS FUNCTION

###############################################################################

def analyze_region(region_name, exp_x_region, exp_y_region, sim_y_interp_region):

    """

    Run DTW, peak finding, and DTW-derived peak assignment for one region.

    """

    dtw_score, dtw_matrix = dtw_distance(

        exp_y_region,

        sim_y_interp_region,

        window=DTW_WINDOW

    )

    dtw_score_norm = dtw_score / len(exp_y_region)

    dtw_path = get_dtw_path(dtw_matrix)

    exp_path_indices = dtw_path[:, 0]

    sim_path_indices = dtw_path[:, 1]

    warped_x_exp = exp_x_region[exp_path_indices]

    warped_exp_y = exp_y_region[exp_path_indices]

    warped_sim_y = sim_y_interp_region[sim_path_indices]

    warped_sim_original_x = exp_x_region[sim_path_indices]

    pearson_original, _ = pearsonr(exp_y_region, sim_y_interp_region)

    pearson_warped, _ = pearsonr(warped_exp_y, warped_sim_y)

    exp_peaks, _ = find_peaks(

        exp_y_region,

        height=EXP_PEAK_HEIGHT,

        prominence=EXP_PEAK_PROMINENCE,

        distance=EXP_PEAK_DISTANCE

    )

    sim_peaks, _ = find_peaks(

        sim_y_interp_region,

        height=SIM_PEAK_HEIGHT,

        prominence=SIM_PEAK_PROMINENCE,

        distance=SIM_PEAK_DISTANCE

    )

    exp_peak_positions = exp_x_region[exp_peaks]

    exp_peak_intensities = exp_y_region[exp_peaks]

    sim_peak_positions = exp_x_region[sim_peaks]

    sim_peak_intensities = sim_y_interp_region[sim_peaks]

    if len(exp_peaks) == 0:

        raise RuntimeError(

            f"No experimental peaks detected in {region_name}. "

            "Lower experimental height/prominence thresholds."

        )

    if len(sim_peaks) == 0:

        raise RuntimeError(

            f"No simulated peaks detected in {region_name}. "

            "Lower simulated height/prominence thresholds."

        )

    ###########################################################################

    # PASS 1: BUILD CANDIDATES AND INITIAL DISTANCE-ONLY ASSIGNMENTS

    ###########################################################################

    candidate_records = []

    initial_matched_exp_peak_indices = set()

    for sim_peak_index in sim_peaks:

        matching_rows = dtw_path[dtw_path[:, 1] == sim_peak_index]

        if len(matching_rows) == 0:

            continue

        aligned_exp_indices = matching_rows[:, 0]

        aligned_exp_position = np.mean(exp_x_region[aligned_exp_indices])

        exp_peak_distances = np.abs(exp_peak_positions - aligned_exp_position)

        candidate_mask = exp_peak_distances <= EXP_PEAK_SNAP_TOLERANCE

        candidate_indices = np.where(candidate_mask)[0]

        if len(candidate_indices) == 0:

            continue

        candidate_distances = exp_peak_distances[candidate_indices]

        if PRINT_FIRST_PASS_ATTEMPTS:

            print("\n==============================================================")

            print(

                f"{region_name.upper()} FIRST PASS MATCH ATTEMPTS "

                f"FOR SIMULATED PEAK {exp_x_region[sim_peak_index]:.2f} cm^-1"

            )

            print("==============================================================")

            print("Exp Peak      DTW/Wavenumber Distance")

            print("-" * 45)

            for i, candidate_index in enumerate(candidate_indices):

                candidate_position = exp_peak_positions[candidate_index]

                candidate_distance = candidate_distances[i]

                print(

                    f"{candidate_position:8.2f}      "

                    f"{candidate_distance:12.6f}"

                )

        initial_best_local_index = np.argmin(candidate_distances)

        initial_best_exp_index = candidate_indices[initial_best_local_index]

        initial_best_position = exp_peak_positions[initial_best_exp_index]

        initial_best_distance = candidate_distances[initial_best_local_index]

        if PRINT_FIRST_PASS_ATTEMPTS:

            print("\nFIRST PASS CHOICE")

            print(f"Simulated Peak:    {exp_x_region[sim_peak_index]:.2f} cm^-1")

            print(f"Experimental Peak: {initial_best_position:.2f} cm^-1")

            print(f"Distance:          {initial_best_distance:.6f}")

        initial_matched_exp_peak_indices.add(

            exp_peaks[initial_best_exp_index]

        )

        candidate_records.append(

            {

                "sim_peak_index": sim_peak_index,

                "aligned_exp_indices": aligned_exp_indices,

                "candidate_indices": candidate_indices,

                "candidate_distances": candidate_distances,

                "num_aligned_points": len(aligned_exp_indices)

            }

        )

    ###########################################################################

    # DETERMINE WHICH EXPERIMENTAL PEAKS WERE LEFT UNCOVERED IN PASS 1

    ###########################################################################

    initial_unmatched_exp_peak_indices = set(exp_peaks) - initial_matched_exp_peak_indices

    ###########################################################################

    # PASS 2: SOFT MANY-TO-ONE ASSIGNMENT

    ###########################################################################

    dtw_peak_assignments = []

    exp_assignment_counts = {}

    combined_min_intensity = min(np.min(exp_y_region), np.min(sim_y_interp_region))

    combined_max_intensity = max(np.max(exp_y_region), np.max(sim_y_interp_region))

    intensity_span = combined_max_intensity - combined_min_intensity

    if intensity_span == 0:

        raise ValueError(f"Intensity span is zero in {region_name}; cannot scale intensity differences.")

    for record in candidate_records:

        sim_peak_index = record["sim_peak_index"]

        candidate_indices = record["candidate_indices"]

        candidate_distances = record["candidate_distances"]

        sim_intensity = sim_y_interp_region[sim_peak_index]

        candidate_exp_intensities = exp_peak_intensities[candidate_indices]

        wavenumber_term = WAVENUMBER_WEIGHT * candidate_distances

        raw_intensity_differences = np.abs(

            candidate_exp_intensities - sim_intensity

        )

        scaled_intensity_differences = (

            raw_intensity_differences / intensity_span

        )

        intensity_term = INTENSITY_WEIGHT * scaled_intensity_differences

        coverage_term = np.zeros_like(candidate_distances)

        for i, candidate_index in enumerate(candidate_indices):

            true_exp_peak_index = exp_peaks[candidate_index]

            if true_exp_peak_index in initial_unmatched_exp_peak_indices:

                coverage_term[i] = -COVERAGE_BONUS

        occupancy_term = np.zeros_like(candidate_distances)

        for i, candidate_index in enumerate(candidate_indices):

            true_exp_peak_index = exp_peaks[candidate_index]

            current_assignment_count = exp_assignment_counts.get(

                true_exp_peak_index,

                0

            )

            occupancy_term[i] = (

                OCCUPANCY_PENALTY_WEIGHT * current_assignment_count

            )

        candidate_scores = (

            wavenumber_term

            + intensity_term

            + coverage_term

            + occupancy_term

        )

        if PRINT_SECOND_PASS_ATTEMPTS:

            print("\n==============================================================")

            print(f"{region_name.upper()} SIMULATED PEAK: {exp_x_region[sim_peak_index]:.2f} cm^-1")

            print("==============================================================")

            print(

                "Exp Peak     "

                "Wave Term    "

                "Intensity Δ  "

                "Scaled Int Δ "

                "Intensity Term   "

                "Coverage Term   "

                "Occupancy Term   "

                "Final Score"

            )

            print("-" * 130)

            for i, candidate_index in enumerate(candidate_indices):

                exp_peak_position = exp_peak_positions[candidate_index]

                print(

                    f"{exp_peak_position:8.2f}     "

                    f"{wavenumber_term[i]:10.3f}   "

                    f"{raw_intensity_differences[i]:10.6f}   "

                    f"{scaled_intensity_differences[i]:10.6f}   "

                    f"{intensity_term[i]:14.6f}   "

                    f"{coverage_term[i]:14.3f}   "

                    f"{occupancy_term[i]:14.3f}   "

                    f"{candidate_scores[i]:11.6f}"

                )

        best_candidate_local_index = np.argmin(candidate_scores)

        nearest_exp_peak_index = candidate_indices[best_candidate_local_index]

        matched_exp_position = exp_peak_positions[nearest_exp_peak_index]

        matched_exp_intensity = exp_peak_intensities[nearest_exp_peak_index]

        sim_position = exp_x_region[sim_peak_index]

        shift = sim_position - matched_exp_position

        matched_true_exp_peak_index = exp_peaks[nearest_exp_peak_index]

        exp_assignment_counts[matched_true_exp_peak_index] = (

            exp_assignment_counts.get(matched_true_exp_peak_index, 0) + 1

        )

        dtw_peak_assignments.append(

            {

                "sim_peak": sim_position,

                "sim_peak_index": sim_peak_index,

                "sim_intensity": sim_intensity,

                "matched_exp_peak": matched_exp_position,

                "matched_exp_peak_index": matched_true_exp_peak_index,

                "matched_exp_intensity": matched_exp_intensity,

                "shift": shift,

                "num_aligned_points": record["num_aligned_points"],

                "assignment_score": candidate_scores[best_candidate_local_index],

                "wavenumber_term": wavenumber_term[best_candidate_local_index],

                "raw_intensity_difference": raw_intensity_differences[best_candidate_local_index],

                "scaled_intensity_difference": scaled_intensity_differences[best_candidate_local_index],

                "intensity_term": intensity_term[best_candidate_local_index],

                "coverage_term": coverage_term[best_candidate_local_index],

                "occupancy_term": occupancy_term[best_candidate_local_index]

            }

        )

    ###########################################################################

    # MATCHED / UNMATCHED PEAKS

    ###########################################################################

    matched_exp_peak_indices = set()

    matched_sim_peak_indices = set()

    for item in dtw_peak_assignments:

        matched_exp_peak_indices.add(

            item["matched_exp_peak_index"]

        )

        matched_sim_peak_indices.add(

            item["sim_peak_index"]

        )

    unmatched_exp_peak_indices = [

        peak_index

        for peak_index in exp_peaks

        if peak_index not in matched_exp_peak_indices

    ]

    unmatched_sim_peak_indices = [

        peak_index

        for peak_index in sim_peaks

        if peak_index not in matched_sim_peak_indices

    ]

    print(f"\n================ {region_name.upper()} DTW RESULTS ================\n")

    print(f"Region analyzed:          {exp_x_region[0]:.2f}–{exp_x_region[-1]:.2f} cm^-1")

    print(f"DTW window:               {DTW_WINDOW} data points")

    print(f"Raw DTW score:            {dtw_score:.4f}")

    print(f"Normalized DTW score:     {dtw_score_norm:.6f}")

    print(f"Original Pearson r:       {pearson_original:.4f}")

    print(f"DTW-warped Pearson r:     {pearson_warped:.4f}")

    print(f"\n========== {region_name.upper()} DTW PEAK SHIFT ASSIGNMENTS ==========\n")

    print("Sim Peak   Matched Exp Peak   Shift   Sim Int   Exp Int   Points")

    print("------------------------------------------------------------------")

    for item in dtw_peak_assignments:

        print(

            f"{item['sim_peak']:8.2f}   "

            f"{item['matched_exp_peak']:16.2f}   "

            f"{item['shift']:7.2f}   "

            f"{item['sim_intensity']:7.3f}   "

            f"{item['matched_exp_intensity']:7.3f}   "

            f"{item['num_aligned_points']:6d}"

        )

    print(f"\n================ {region_name.upper()} PEAK COUNTS ================\n")

    print(f"Total experimental peaks found: {len(exp_peaks)}")

    print(f"Total simulated peaks found:    {len(sim_peaks)}")

    return {

        "region_name": region_name,

        "x": exp_x_region,

        "exp_y": exp_y_region,

        "sim_y": sim_y_interp_region,

        "dtw_score": dtw_score,

        "dtw_score_norm": dtw_score_norm,

        "dtw_path": dtw_path,

        "exp_path_indices": exp_path_indices,

        "sim_path_indices": sim_path_indices,

        "warped_x_exp": warped_x_exp,

        "warped_exp_y": warped_exp_y,

        "warped_sim_y": warped_sim_y,

        "warped_sim_original_x": warped_sim_original_x,

        "pearson_original": pearson_original,

        "pearson_warped": pearson_warped,

        "exp_peaks": exp_peaks,

        "sim_peaks": sim_peaks,

        "exp_peak_positions": exp_peak_positions,

        "exp_peak_intensities": exp_peak_intensities,

        "sim_peak_positions": sim_peak_positions,

        "sim_peak_intensities": sim_peak_intensities,

        "dtw_peak_assignments": dtw_peak_assignments,

        "matched_exp_peak_indices": matched_exp_peak_indices,

        "matched_sim_peak_indices": matched_sim_peak_indices,

        "unmatched_exp_peak_indices": unmatched_exp_peak_indices,

        "unmatched_sim_peak_indices": unmatched_sim_peak_indices,

    }

###############################################################################

\# PREPARE REGION

###############################################################################

fingerprint_xmin, fingerprint_xmax = FINGERPRINT_REGION

fp_exp_x, fp_exp_y, fp_sim_y = prepare_region(

    exp_x_raw,

    exp_y_raw,

    sim_x_raw,

    sim_y_raw,

    fingerprint_xmin,

    fingerprint_xmax

)

###############################################################################

\# ANALYZE REGION

###############################################################################

fingerprint_results = analyze_region(

    "fingerprint",

    fp_exp_x,

    fp_exp_y,

    fp_sim_y

)

###############################################################################

\# SCORING METRICS

###############################################################################

fp_match_fraction, fp_peak_rms = calculate_peak_metrics(fingerprint_results)

print("\n================ SCORING SUMMARY ================\n")

print("Fingerprint Region")

print(f"Peak match fraction:     {fp_match_fraction:.4f}")

print(f"Pearson original:        {fingerprint_results['pearson_original']:.4f}")

print(f"Pearson DTW-warped:      {fingerprint_results['pearson_warped']:.4f}")

print(f"Peak shift RMSD:         {fp_peak_rms:.4f} cm^-1")

###############################################################################

\# PLOTTING HELPERS

###############################################################################

def get_marker_arrays(results):

    x = results["x"]

    exp_y = results["exp_y"]

    sim_y = results["sim_y"]

    matched_exp_indices = list(results["matched_exp_peak_indices"])

    matched_sim_indices = list(results["matched_sim_peak_indices"])

    unmatched_exp_indices = results["unmatched_exp_peak_indices"]

    unmatched_sim_indices = results["unmatched_sim_peak_indices"]

    return {

        "matched_exp_positions": x[matched_exp_indices],

        "matched_exp_intensities": exp_y[matched_exp_indices],

        "unmatched_exp_positions": x[unmatched_exp_indices],

        "unmatched_exp_intensities": exp_y[unmatched_exp_indices],

        "matched_sim_positions": x[matched_sim_indices],

        "matched_sim_intensities": sim_y[matched_sim_indices],

        "unmatched_sim_positions": x[unmatched_sim_indices],

        "unmatched_sim_intensities": sim_y[unmatched_sim_indices],

    }

def plot_region_on_axis(ax, results, include_legend=False):

    x = results["x"]

    exp_y = results["exp_y"]

    sim_y = results["sim_y"]

    markers = get_marker_arrays(results)

    ax.plot(

        x,

        exp_y,

        linewidth=GRAPH["spectrum_linewidth"],

        label="Experimental"

    )

    ax.plot(

        x,

        sim_y,

        linewidth=GRAPH["spectrum_linewidth"],

        label="Simulated"

    )

    ax.scatter(

        markers["matched_exp_positions"],

        markers["matched_exp_intensities"],

        marker="o",

        s=GRAPH["marker_size"],

        color=GRAPH["matched_color"],

        edgecolors=GRAPH["matched_color"],

        linewidths=1.5,

        zorder=5,

        label="Matched Experimental Peaks"

    )

    ax.scatter(

        markers["unmatched_exp_positions"],

        markers["unmatched_exp_intensities"],

        marker="o",

        s=GRAPH["marker_size"],

        color=GRAPH["unmatched_color"],

        edgecolors=GRAPH["unmatched_color"],

        linewidths=1.5,

        zorder=5,

        label="Unmatched Experimental Peaks"

    )

    ax.scatter(

        markers["matched_sim_positions"],

        markers["matched_sim_intensities"],

        marker="x",

        s=GRAPH["marker_size"],

        color=GRAPH["matched_color"],

        linewidths=2.0,

        zorder=5,

        label="Matched Simulated Peaks"

    )

    ax.scatter(

        markers["unmatched_sim_positions"],

        markers["unmatched_sim_intensities"],

        marker="x",

        s=GRAPH["marker_size"],

        color=GRAPH["unmatched_color"],

        linewidths=2.0,

        zorder=5,

        label="Unmatched Simulated Peaks"

    )

    for item in results["dtw_peak_assignments"]:

        sim_peak = item["sim_peak"]

        exp_peak = item["matched_exp_peak"]

        sim_y_value = np.interp(sim_peak, x, sim_y)

        exp_y_value = np.interp(exp_peak, x, exp_y)

        ax.plot(

            [exp_peak, sim_peak],

            [exp_y_value, sim_y_value],

            linestyle="--",

            linewidth=GRAPH["match_linewidth"],

            color=GRAPH["match_line_color"],

            alpha=GRAPH["match_line_alpha"]

        )

    ax.tick_params(axis="both", labelsize=GRAPH["tick_fontsize"])

    for tick in ax.get_xticklabels():

        tick.set_fontweight("bold")

    for tick in ax.get_yticklabels():

        tick.set_fontweight("bold")

    if include_legend:

        legend = ax.legend(fontsize=GRAPH["legend_fontsize"])

        for text in legend.get_texts():

            text.set_fontweight("bold")

###############################################################################

\# PLOT 1: ORIGINAL FINGERPRINT SPECTRA

###############################################################################

fig, ax1 = plt.subplots(

    figsize=GRAPH["figsize"],

    dpi=GRAPH["dpi"]

)

plot_region_on_axis(ax1, fingerprint_results, include_legend=True)

ax1.set_xlim(FINGERPRINT_REGION[0], FINGERPRINT_REGION[1])

###############################################################################

\# REMOVE Y-AXIS TICKS / NUMBERS BUT KEEP LABEL

###############################################################################

ax1.set_yticks([])

ax1.tick_params(axis="y", left=False, labelleft=False)

ax1.spines["top"].set_visible(False)

ax1.spines["right"].set_visible(False)

ax1.spines["left"].set_visible(False)

ax1.spines["bottom"].set_visible(True)

fig.supxlabel(

    "Raman Shift (cm$^{-1}$)",

    fontsize=GRAPH["axis_fontsize"],

    fontweight="bold"

)

ax1.set_ylabel(

    "Vector-Normalized Intensity",

    fontsize=GRAPH["axis_fontsize"],

    fontweight="bold"

)

fig.suptitle(

    GRAPH["overall_original_title"],

    fontsize=GRAPH["axis_fontsize"],

    fontweight="bold"

)

if SAVE_ORIGINAL_FIGURE:

    plt.savefig(ORIGINAL_FIGURE_NAME, dpi=GRAPH["save_dpi"], bbox_inches="tight")

plt.tight_layout()

#plt.show()

###############################################################################

\# PLOT 2: DTW-WARPED FINGERPRINT SPECTRA

###############################################################################

fig, ax1 = plt.subplots(

    figsize=GRAPH["figsize"],

    dpi=GRAPH["dpi"]

)

results = fingerprint_results

alignment_index = np.arange(len(results["warped_exp_y"]))

ax1.plot(

    alignment_index,

    results["warped_exp_y"],

    linewidth=GRAPH["spectrum_linewidth"],

    label="Experimental DTW Path"

)

ax1.plot(

    alignment_index,

    results["warped_sim_y"],

    linewidth=GRAPH["spectrum_linewidth"],

    label="Simulated Warped by DTW"

)

for item in results["dtw_peak_assignments"]:

    exp_idx = item["matched_exp_peak_index"]

    sim_idx = item["sim_peak_index"]

    exp_path_matches = np.where(results["exp_path_indices"] == exp_idx)[0]

    sim_path_matches = np.where(results["sim_path_indices"] == sim_idx)[0]

    if len(exp_path_matches) == 0 or len(sim_path_matches) == 0:

        continue

    exp_warped_idx = exp_path_matches[len(exp_path_matches) // 2]

    sim_warped_idx = sim_path_matches[len(sim_path_matches) // 2]

    ax1.scatter(

        exp_warped_idx,

        results["exp_y"][exp_idx],

        marker="o",

        s=GRAPH["marker_size"],

        color=GRAPH["matched_color"],

        edgecolors=GRAPH["matched_color"],

        linewidths=1.5,

        zorder=5

    )

    ax1.scatter(

        sim_warped_idx,

        results["sim_y"][sim_idx],

        marker="x",

        s=GRAPH["marker_size"],

        color=GRAPH["matched_color"],

        linewidths=1.5,

        zorder=5

    )

    ax1.plot(

        [exp_warped_idx, sim_warped_idx],

        [results["exp_y"][exp_idx], results["sim_y"][sim_idx]],

        linestyle="--",

        linewidth=GRAPH["warped_match_linewidth"],

        color=GRAPH["match_line_color"],

        alpha=GRAPH["match_line_alpha"]

    )

ax1.tick_params(axis="both", labelsize=GRAPH["tick_fontsize"])

for tick in ax1.get_xticklabels():

    tick.set_fontweight("bold")

for tick in ax1.get_yticklabels():

    tick.set_fontweight("bold")

###############################################################################

\# MATCH ORIGINAL FIGURE FORMATTING

###############################################################################

ax1.set_yticks([])

ax1.tick_params(axis="y", left=False, labelleft=False)

ax1.spines["top"].set_visible(False)

ax1.spines["right"].set_visible(False)

ax1.spines["left"].set_visible(False)

ax1.spines["bottom"].set_visible(True)

ax1.set_ylabel(

    "Vector-Normalized Intensity",

    fontsize=GRAPH["axis_fontsize"],

    fontweight="bold"

)

fig.supxlabel(

    "DTW Alignment Index",

    fontsize=GRAPH["axis_fontsize"],

    fontweight="bold"

)

fig.suptitle(

    GRAPH["overall_warped_title"],

    fontsize=GRAPH["axis_fontsize"],

    fontweight="bold"

)

legend = ax1.legend(fontsize=GRAPH["legend_fontsize"])

for text in legend.get_texts():

    text.set_fontweight("bold")

if SAVE_WARPED_FIGURE:

    plt.savefig(WARPED_FIGURE_NAME, dpi=GRAPH["save_dpi"], bbox_inches="tight")

plt.tight_layout()

#plt.show()
